# Type I Interferon Signaling Drives Microglial Dysfunction and Senescence in Human iPSC Models of Down Syndrome and Alzheimer’s Disease

**DOI:** 10.1101/2021.12.22.473858

**Authors:** Mengmeng Jin, Ranjie Xu, Le Wang, Mahabub Maraj Alam, Ziyuan Ma, Sining Zhu, Alessandra C. Martini, Azadeh Jadali, Matteo Bernabucci, Ping Xie, Kelvin Kwan, Zhiping P. Pang, Elizabeth Head, Ying Liu, Ronald P. Hart, Peng Jiang

## Abstract

Microglia are critical for brain development and play a central role in Alzheimer’s disease (AD) etiology. Down syndrome (DS), also known as trisomy 21, is the most common genetic origin of intellectual disability and the most common risk factor for AD. Surprisingly, little information is available on the impact of trisomy of human chromosome 21 (Hsa21) on microglia in DS brain development and AD in DS (DSAD). Using our new induced pluripotent stem cell (iPSC)-based human microglia-containing cerebral organoid and chimeric mouse brain models, here we report that DS microglia exhibit enhanced synaptic pruning function during brain development. Consequently, electrophysiological recordings demonstrate that DS microglial mouse chimeras show impaired synaptic functions, as compared to control microglial chimeras. Upon being exposed to human brain tissue-derived soluble pathological tau, DS microglia display dystrophic phenotypes in chimeric mouse brains, recapitulating microglial responses seen in human AD and DSAD brain tissues. Further flow cytometry, single-cell RNA-sequencing, and immunohistological analyses of chimeric mouse brains demonstrate that DS microglia undergo cellular senescence and exhibit elevated type I interferon signaling after being challenged by pathological tau. Mechanistically, we find that shRNA-mediated knockdown of Hsa21encoded type I interferon receptor genes, *IFNARs*, rescues the defective DS microglial phenotypes both during brain development and in response to pathological tau. Our findings provide first *in vivo* evidence supporting a paradigm shifting theory that human microglia respond to pathological tau by exhibiting accelerated senescence and dystrophic phenotypes. Our results further suggest that targeting IFNARs may improve microglial functions during DS brain development and prevent human microglial senescence in DS individuals with AD.

## Introduction

Down syndrome (DS), caused by trisomy of human chromosome 21 (Hsa21), is both the most common genetic cause of abnormal brain development and learning disability in children, and the most common risk factor for Alzheimer’s disease (AD) (Lott and Head, 2019; Parker et al., 2010; Wiseman et al., 2015). As the resident macrophages of the central nervous system (CNS), microglia play critical roles in the maintenance of CNS homeostasis, modulation of neuronal development, and remodeling of neuronal synapses, thereby critically regulating synaptic plasticity and learning and memory process (Bar and Barak, 2019; Casano and Peri, 2015; Li and Barres, 2018; Morris et al., 2013; Parkhurst et al., 2013; Wang et al., 2020). Moreover, microglial dysfunction is a central mechanism in AD etiology, and many AD risk genes are highly and sometimes exclusively expressed by microglia (Gosselin et al., 2017; Hansen et al., 2018; Holtman et al., 2017). Therefore, targeting microglia has enormous potential for improving DS brain development and treating AD. Unfortunately, the precise contributions of microglia to brain development in DS and AD in DS (henceforth referred to as DSAD) as well as underlying molecular mechanisms are poorly understood, which hampers the development of therapeutics.

Limited information is available on the influences of trisomy 21 on microglia during brain development and degeneration. A recent study using a mouse model of DS reports that microglia are activated during brain development, as indicated by altered morphology as well as expression of activation markers and pro-inflammatory cytokines (Pinto et al., 2020). Consistently, a study using post- mortem human brain tissues from fetuses to young adults with DS also shows elevated microglial activation and expression of brain inflammatory cytokines, as compared to the brain tissues from control subjects (Flores-Aguilar et al., 2020). Despite these characterizations at cellular levels, how trisomy of Hsa21 genes alter microglial development and functions remains largely unknown. The association between DS and AD is largely due to overexpression of the amyloid precursor protein (APP), whose gene is located on Hsa21 (Doran et al., 2017; Head et al., 2003; Lott and Head, 2019; Prasher et al., 1998). As early as 35 - 40 years of age, tau pathological changes are observed in the hippocampus in adults with DS (Head et al., 2003; Lott and Head, 2019). While the aggregation of amyloid-beta (Aβ) precedes that of tau, tau protein pathology commences in humans much sooner than was previously thought (Braak and Del Tredici, 2015). Contrary to the marked microglial activation reported in amyloidogenic mouse models (Jimenez et al., 2008; Meyer-Luehmann and Prinz, 2015), in brain tissue derived from AD patients, brain regions particularly relevant in AD development, such as the hippocampal formation, exhibit low and late Aβ pathology, whereas hyperphosphorylated tau (p-tau) accumulates starting in the early stages of the disease (Braak and Del Tredici, 2015; Sanchez-Mejias et al., 2016). Intriguingly, studies using human brain tissue derived from both AD and DSAD patients showed that degenerating neuronal structures positive for p-tau invariably colocalized with severely dystrophic and senescent rather than activated microglial cells (Shahidehpour et al., 2021; Streit et al., 2009; Xue and Streit, 2011). In DSAD, microglial phenotypes shift with age and the presence of AD pathology, showing the presence of higher numbers of dystrophic microglia (Martini et al., 2020). Thus, the preferential accumulation of p-tau over Aβ plaques in specific brain regions could induce a totally different microglial response than merely activation and inflammation.

Modeling the DS-related cellular phenotypes and elucidation of the molecular mechanisms underlying DS disease pathogenesis is challenging. This is because functional DS human brain tissue is relatively inaccessible, and DS mouse models often show variations and discrepancies in modeling DS-related phenotypes due to the trisomy of different subsets of Hsa21 orthologous genes in different mouse models (Belichenko et al., 2015; Das and Reeves, 2011; Xu et al., 2019). In addition, none of the mouse models of DS reliably reproduces Aβ or tau pathology even in aged animals (Choong et al., 2015). It is also important to note that rodent microglia are not able to fully mirror the properties of human microglia. Transcriptomic profiling of human and murine microglia reveals species-specific expression patterns in hundreds of genes, including genes involved in brain development, immune function, and phagocytic function (Galatro et al., 2017; Geirsdottir et al., 2019; Gosselin et al., 2017; Jiang et al., 2020). A limited overlap was also observed in microglial genes regulated during aging and neurodegeneration between mice and humans, indicating that human and mouse microglia age differently under normal and diseased conditions (Friedman et al., 2018; Galatro et al., 2017).

Furthermore, previous studies show that a tissue environment-dependent transcriptional network specifies human microglia identity, and *in vitro* environments drastically alter the human microglia transcriptome (Gosselin et al., 2017). These findings argue for the utilization of species-specific and *in vivo* research tools to investigate microglial functions in human brain development, aging, and neurodegeneration (Smith and Dragunow, 2014).

Recent advances in stem cell technology have led to the efficient generation of microglia from human induced pluripotent stem cells (hiPSCs) (Abud et al., 2017; Brownjohn et al., 2018; Douvaras et al., 2017; Haenseler et al., 2017; Jiang et al., 2020; Muffat et al., 2016; Pandya et al., 2017), providing an unlimited source of human microglia to study their pathophysiology. Very recently, we have developed new hiPSC-based microglia-containing cerebral organoid and chimeric mouse brain models (Jiang et al., 2020; Xu et al., 2021; Xu et al., 2020), in which hiPSC-derived microglia undergo maturation and develop appropriate functions. With DS hiPSC established in our previous studies (Chen et al., 2014; Xu et al., 2019), these new models provide unprecedented opportunities to investigate the role of human microglia in DS brain development and DSAD. In this study, we demonstrate that DS microglia show defective development and functions compared to control microglia. Moreover, in human-mouse microglial chimeric brains, we discover that DSAD human brain tissue-derived pathological tau induces senescence in DS microglia, recapitulating microglial responses in human AD and DSAD brain tissue. Importantly, inhibiting the expression of Hsa21-encoded type I interferon receptor genes, *IFNARs*, improves the defective DS microglia functions during brain development and prevents DS microglial senescence in response to pathological tau.

## Results

### Generation and characterization of DS hiPSC-derived primitive macrophage progenitors

Microglia arise from the embryonic yolk sac (YS) as *Myb*-independent primitive macrophage progenitors (PMPs) and migrate into the brain to complete differentiation (Li and Barres, 2018). Using a published protocol (Haenseler et al., 2017), we first derived PMPs from the control (Cont) and DS hiPSCs that were generated and fully characterized in our previous studies (Chen et al., 2014; Xu et al., 2019), including two DS hiPSC lines (DS1 and DS2) and isogenic disomic (Di)-DS3 and trisomic (Tri)- DS3 (Table S1). Then, we confirmed the identity of these hiPSC-derived PMPs by immunostaining and bulk RNA sequencing (RNA-seq) (Fig. 1A). We found that over 94% of Cont and DS hiPSC-derived PMPs expressed CD235, a YS primitive hematopoietic progenitor marker, and CD43, a hematopoietic progenitor-like cell marker. These PMPs were also highly proliferative, as indicated by expressing Ki67, a cell proliferation marker (Fig. 1B-C). These DS PMPs exhibited trisomy of Hsa21, as demonstrated by fluorescence *in situ* hybridization assay and gene copy number assay for Hsa21 genes, *OLIG2, IFNAR1,* and *IFNAR2* (Fig. 1B, S1A).

**Fig 1.**
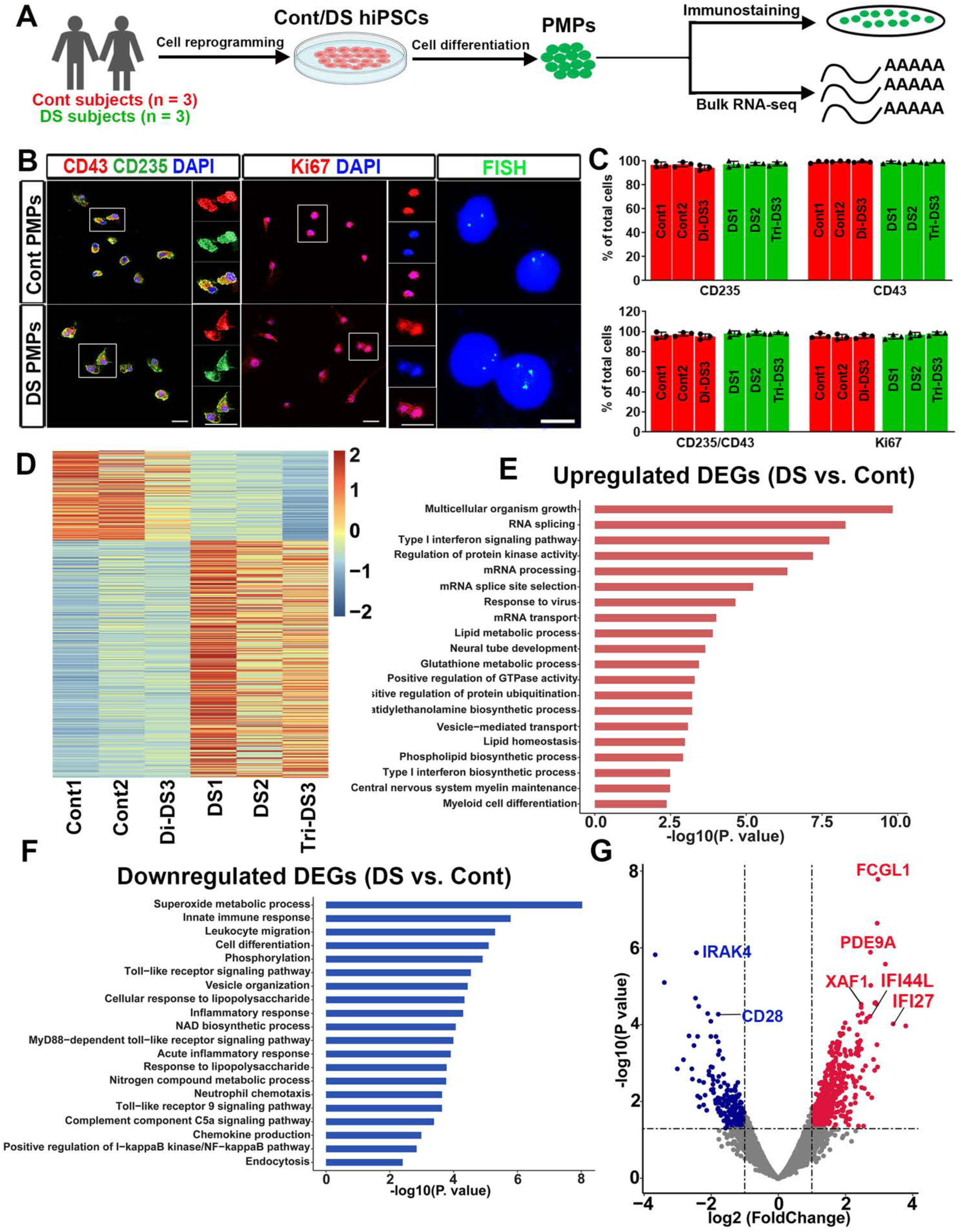
Generation and characterization of DS hiPSC-derived PMPs. (A) Schematic representation of the generation and characterization of PMPs. (B) Representative images of CD235^+^, CD43^+^, Ki67^+^ cells, and fluorescence in situ hybridization (FISH) analysis in PMPs. Scale bars: 20 μm,10 μm, and 5 μm in the original and enlarged images. (C) Quantification of CD235^+^, CD43^+^, CD235^+^/CD43^+^, and Ki67^+^ PMPs derived from the three pairs of Cont and DS hiPSC lines (n=3, each experiment was repeated three times). Data are presented as mean ± SEM. (D) The heatmap showing all DEGs between Cont and DS PMPs. (E-F) GO analyses of the upregulated and downregulated DEGs in the PMPs. (G) A volcano plot illustrating downregulated (blue) and upregulated (red) DEGs in PMPs.

Derivation of PMPs from both Cont and DS hiPSCs with high purity allowed us to compare their gene expression profiles without further purification steps. We performed RNA-seq using PMPs differentiated from the three pairs of iPSC cells. A total of 775 differentially expressed genes (DEGs) between Cont vs. DS PMPs were identified, including 562 upregulated genes and 213 downregulated genes (Fig. 1D). Gene ontology (GO) analyses of the upregulated DEGs in DS PMPs showed significant enrichment of biological processes particularly related to type I interferon (IFN) signaling pathway, neural tube development, and cell differentiation (Fig. 1E, Table S2). Of note, the type I IFN signaling pathway activated in DS PMPs is consistent with previous studies that showed Hsa21 constantly activates interferon signaling (Fig. 1E, Table S2) (Sullivan et al., 2016; Waugh et al., 2019). In addition, we found that the downregulated genes in DS PMPs were enriched in pathways relevant to immune signaling, such as innate immune response, Toll-like receptor signaling pathway, and NF- κB pathway (Fig. 1F, Table S2). Moreover, the volcano plot showed that the expression of *XAF1*, *IFI27*, and *IFI44L* involved in the type I IFN signaling pathway were upregulated in DS PMPs (Fig. 1G).

Consistently, qPCR analysis showed that the expression of Hsa21 genes that encode type I IFN receptors (IFNARs), *IFNAR*1 and *IFNAR*2, were increased in DS PMPs compared to Cont PMPs (Fig. S1B). Furthermore, expression of *IRAK4* and *FCGL1* – which are involved in T cell activation (Borowski et al., 2007; Suzuki et al., 2006; Wang et al., 2019) – was also altered in DS PMPs compared to Cont PMPs (Fig. 1G). Taken together, these findings suggest that aberrant development of myeloid cell lineage in DS can be detected as early as the PMP stage.

Interestingly, we found that 22 DEGs are Hsa21 genes. Notably, among these 22 genes, the tetratricopeptide repeat domain 3 (TTC3) is an E3 ligase that interacts with Akt, and DS neural cells exhibit elevated TTC3 expression (Suizu et al., 2009), which is consistent with our observations. The regulator of calcineurin 1 (RCAN1) was reported to contribute to neurotrophin receptor trafficking and developmental abnormalities in DS (Patel et al., 2015). Here, we observed that DS PMPs had higher expression of this gene. Intersectin-1 (ITSN1) is involved in endocytic functions (Keating et al., 2006). DS individuals show elevated expression of ITSN1 in the brain, which has been shown to contribute to the development of AD in DS (Hunter et al., 2011; Keating et al., 2006; Ling et al., 2014). Our PMPs also showed higher expression of *ITSN1*. Additionally, *PWP2* and *PDE9A* were found to express at higher levels in our DS PMPs. PWP2 contributes to ribosome biogenesis that is upregulated in lymphoblastoid cell lines derived from DS individuals between the ages of 24 and 41 (Sultan et al., 2007). PDE9A participates in the cyclic guanosine 3’,5’-monophosphate (cGMP) catabolic process, expression of which is upregulated in the cerebellum of the Ts65Dn mouse model of DS (Spellman et al., 2013). Altogether, our results indicate that these Cont or DS hiPSC-derived PMPs exhibit myeloid cell lineage characteristics and that DS PMPs showed higher expression of many Hsa21 genes compared to Cont PMPs.

### Abnormal development and function of DS microglia in cerebral organoids, human brain tissues, and human-mouse microglial chimeras

We recently developed a novel microglia-containing cerebral organoid model that provides a convenient tool to study human microglial functions in neurodevelopmental disorders (Xu et al., 2021). As shown in Fig. 2A, here we used PMPs derived from the three pairs of Cont and DS hiPSCs and primitive neural progenitor cells (pNPCs) derived from Cont hiPSC cells (Xu et al., 2019). Then, hiPSC-derived pNPCs and PMPs were mixed to develop organoids, as described in our recent study (Xu et al., 2021). By week 4, both Cont and DS organoids contained cells that expressed CD45, which is a marker for all nucleated cells of the hemopoietic lineage (Carson et al., 1998) and labels microglia in the organoids (Fig. 2A-B). Similar percentages of CD45^+^ microglia were seen in both Cont and DS organoids (Fig. 2C). Moreover, we found cells in organoids that expressed TMEM119 – a highly specific marker expressed only by microglia and not by other macrophages (Bennett et al., 2016) (Fig. 2D). These results demonstrate that both Cont and DS PMPs can differentiate into microglia in cerebral organoids at a similar efficiency. Moreover, we stained the Cont and DS organoids with MAP2 and observed many MAP2^+^ neurons in the organoids (Fig. S2A), indicating successful neuronal differentiation in the Cont and DS organoids at week 8.

**Fig 2.**
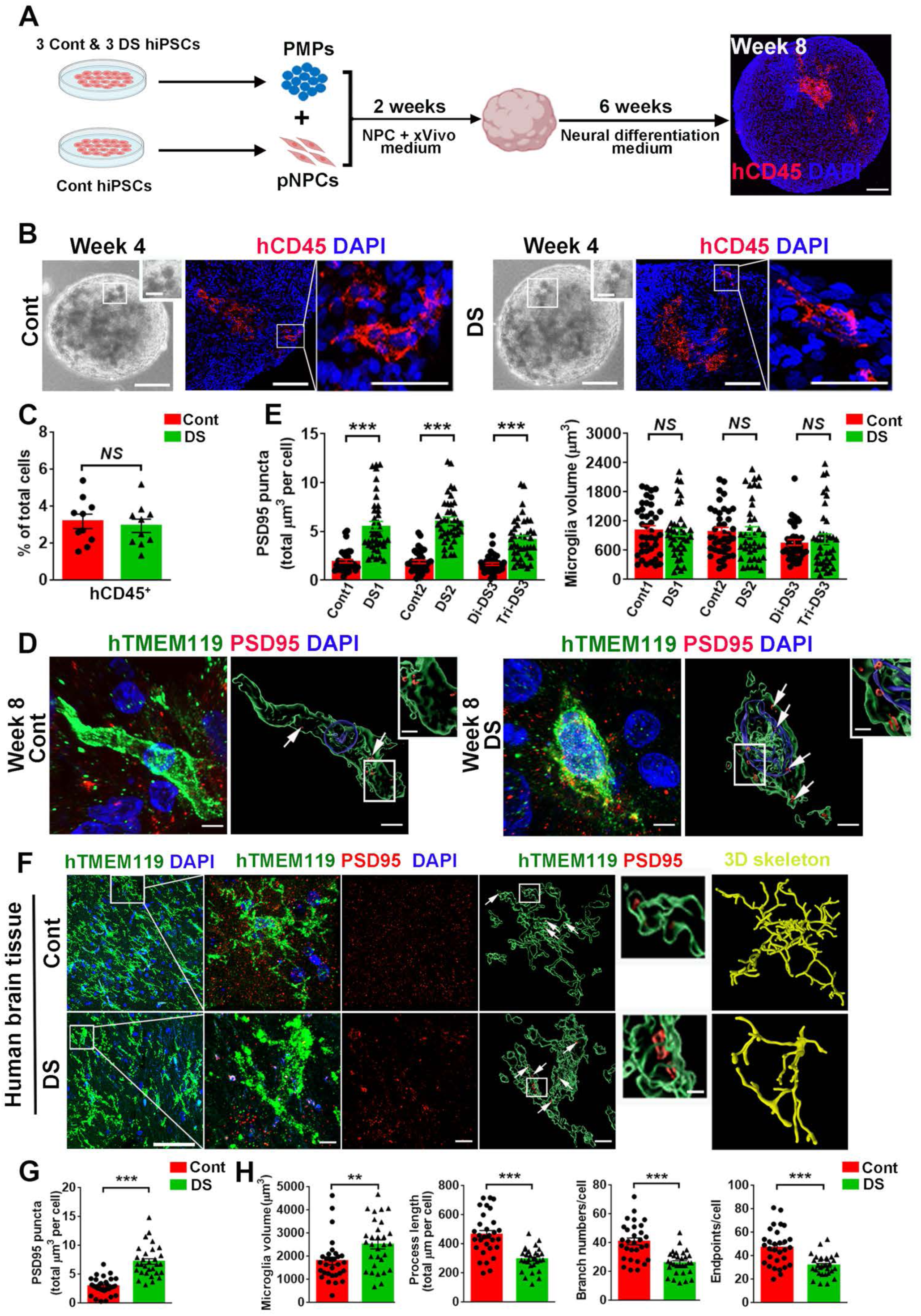
Abnormal development and function of DS microglia in cerebral organoids and human brain tissues. (A) A schematic representation of developing microglia-containing cerebral organoids by co-culture of Cont pNPCs and Cont or DS hiPSC-derived PMPs under 3D conditions. This drawing created using BioRender.com. Scale bar: 100 μm. (B) Representative images of hCD45^+^ microglia in Cont and DS microglia-containing cerebral organoids. (C) Quantification of the percentage of hCD45^+^ cells in total DAPI^+^ cells in organoids at week 8 (n=10, the data were pooled from the three pairs of microglia-containing organoids). Student’s t test, *NS*, not significant. Data are presented as mean ± SEM. (D) Representative raw fluorescent super-resolution and 3D surface rendered images showing colocalization of hTMEM119^+^ and PSD95^+^ staining in week 8 organoids. Scale bars: 5 μm and 1 μm in the original and enlarged images, respectively. (E) Quantification of PSD95^+^ puncta in hCD45^+^ microglia and hTMEM119^+^ microglial volume at week 8 (n = 3, the experiments were repeated 3 times. 3-4 organoids generated from the three pairs of hiPSC lines were used for each experiment, and 4-8 microglia from each organoid were analyzed. Each dot represents one microglia). Student’s t test, \*\*\**P* < 0.001, *NS*, not significant. Data are presented as mean ± SEM. (F) Representative raw fluorescent super-resolution, 3D surface rendering from the raw image, and 3D skeletonization images of hTMEM119 and PSD95 staining in hippocampal slices from a control and a DS individual. Arrows indicate PSD95^+^ puncta. Scale bar: 50 μm, 5 μm and 1 μm in the original and enlarged images, respectively. (G, H) Quantification of PSD95^+^ puncta inside microglia, microglial volume, process length, branch numbers, and endpoints (n=30 microglia per group) of human hippocampal slices. Student’s t test, \*\**P* < 0.01 and \*\*\**P* < 0.001. Data are presented as mean ± SEM.

One of the major functions of microglia is synaptic pruning, through which microglia engulf synapses and shape synaptic development (Paolicelli et al., 2011; Wolf et al., 2017). As shown in Figs. 2D and 2E, using super-resolution imaging analysis, we found a significantly higher number of PSD95^+^ puncta a post-synaptic marker, within hTMEM119^+^ DS microglia, as compared to hTMEM119^+^ Cont microglia, suggesting that DS microglia had enhanced synaptic pruning function. This finding is consistent with a recent study in a mouse model of DS that reported an increased number of excitatory synaptic markers engulfed by microglia in DS mice relative to that in wild-type mice (Pinto et al., 2020). In addition, to further verify the enhanced synaptic pruning function of DS microglia in a trisomic environment, we generate trisomic organoids by mixing the PMPs and primitive neural progenitor cells (pNPCs) derived from the three DS hiPSC lines used in this study. At week 8, we examined neuronal differentiation with MAP2 staining and observed abundant MAP2^+^ neurons in the trisomic organoids (Fig. S2B). We found that like DS microglia developed in a euploid environment, DS microglia in the trisomic organoids also exhibited enhanced synaptic pruning function compared to control microglia (Fig. S2C-S2D). Interestingly, the recent study (Pinto et al., 2020) also reported that microglia in DS mice had larger soma size and branching impairment in comparison to microglia in wild-type mice.

However, in the cerebral organoids, microglia extend fewer processes and do not show morphology as highly ramified as they do *in vivo* (Benito-Kwiecinski and Lancaster, 2020; Ormel et al., 2018; Xu et al., 2021). Consequently, we did not observe any significant morphological differences between DS vs. Cont microglia in cell size, as indicated by the quantification of microglia cell volume (Fig. 2D-E). To better understand the phenotypic changes of DS microglia during early brain development, we analyzed microglia in post-mortem hippocampal tissues from DS and control subjects at ages of less than 1 year (Table S3). Like our brain organoid model, we observed that DS microglia in the hippocampal tissue also had a higher number of PSD95^+^ puncta, as compared to control microglia (Fig. 2F, 2G). Importantly, DS microglia also exhibited a reactive morphology as indicated by increased microglial volume as well as reduced numbers of branches, process lengths, and endpoints (Fig. 2F, 2H), which are consistent with the recent observations (Flores-Aguilar et al., 2020; Pinto et al., 2020). Taken together, these results demonstrate that our organoid model recapitulates the enhanced synaptic pruning function of DS microglia but is not optimal for modeling the morphological changes of DS microglia.

We then transplanted PMPs derived from the three pairs of Cont and DS hiPSCs into the brains of postnatal day 0 (P0) *Rag2*^−/−^ *IL2r*γ ^−/−^ *hCSF1*^KI^ immunodeficient mice to generate human microglial mouse chimeras (Xu et al., 2020). The PMPs were transplanted into the white matter overlying the hippocampus and sites within the hippocampal formation, as described in our recent study (Xu et al., 2020). As shown in Fig. 3A, at 8 weeks post-transplantation, both xenografted Cont and DS microglia labeled by human specific TMEM119 (hTMEM119) were found to migrate throughout the brain along the corpus callosum and disperse in multiple brain regions, including the cerebral cortex and hippocampus. At 3-4 months post-transplantation, hTMEM119^+^ Cont and DS microglia were widely distributed throughout the hippocampus (Fig. 3A). As shown in Fig. S3A, S3C, in both Cont and DS microglia chimeras, the vast majority of hN^+^ cells (> 87%) were hTMEM119^+^, indicating that both Cont and DS PMPs efficiently differentiated into microglia. We next assessed the proliferation of engrafted cells by staining the proliferative marker Ki67. Both Cont and DS microglia were capable of proliferating in the mouse brains at 8 weeks post-transplantation, and no difference was observed between them (Fig. S3A, S3C). As expected, both DS and Cont microglia showed complex processes in the chimeric mouse brains and developed increasingly complex morphology along with the developing mouse brain from 4 weeks to 3 months old (Fig. 3B-C). Interestingly, in the grey matter (GM), 3D skeleton analysis indicated that DS microglia had less intricate morphology than Cont microglia, as indicated by enlarged cell volume, shortened process length, fewer endpoints, and branch numbers (Fig. 3B-C, S3D). As opposed to the GM, microglia morphology is not as complex in the white matter (WM) (Fig. S3B). We found that in the WM, the cell volume of Cont and DS microglia was similar, but the DS microglia showed significantly fewer branch numbers, reduced endpoints, and shorter process length (Fig. S3E).

**Fig 3.**
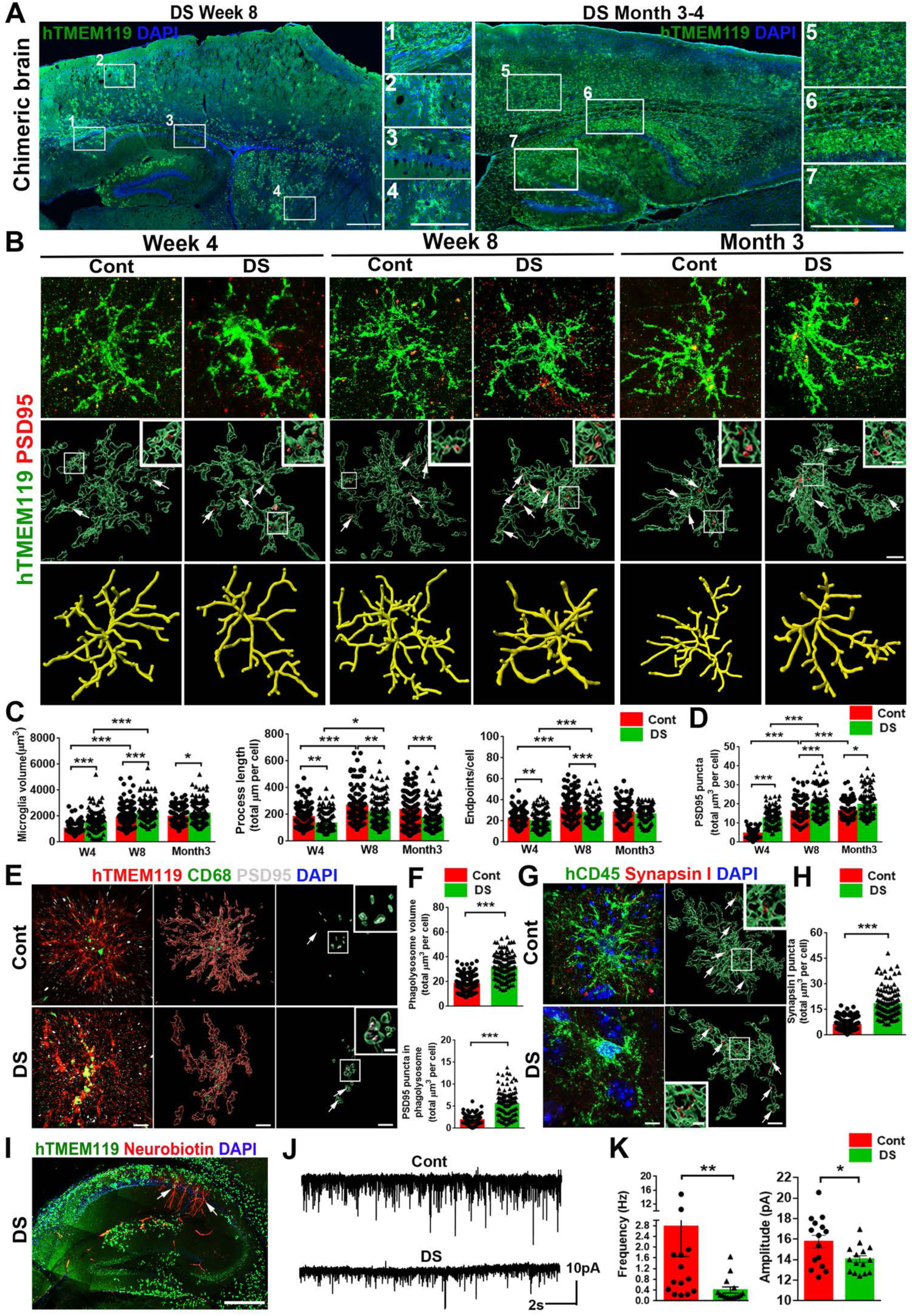
Modeling DS microglial phenotypes in human-mouse microglial chimeras. (A) Representative images from sagittal brain sections showing the distribution of transplanted DS hiPSC-derived microglia at week 8 and 3 to 4 months. Scale bar: 1 mm and 500 μm in the original and enlarged images, respectively. (B) Representative super-resolution, 3D surface rendering, and 3D skeletonization images showing colocalization of hTMEM119^+^ and PSD95^+^ staining at 4 weeks, 8 weeks, and 3 months post-transplantation. Arrows indicate PSD95^+^ puncta. Scale bar: 5 μm and 1 μm in the original and enlarged images, respectively. (C, D) Quantification of microglial volume, process length, endpoints, and PSD95^+^ puncta inside microglia (n=120-160 microglia from 3-5 mice per group). Student’s t test, \**P* < 0.05, \*\**P* < 0.01, and \*\*\**P* < 0.001. Data are presented as mean ± SEM. (E) Representative super-resolution and 3D surface rendered images showing colocalization of hTMEM119, CD68, and PSD95 staining of week 8 chimeras. Arrows indicate PSD95^+^ puncta in the CD68^+^ phagolysosome. Scale bar: 5 μm and 1 μm in the original and enlarged images, respectively. (F) Quantification of CD68^+^ phagolysosome volume and PSD95^+^ puncta inside CD68^+^ phagolysosomes (n=120 microglia from 4 mice per group). Student’s t test, \*\*\**P* < 0.001. Data are presented as mean ± SEM. (G) Representative super-resolution and 3D surface rendered images of hCD45 and synapsin I staining in week 8 of chimeras. Arrows indicate synapsin I^+^ puncta. Scale bar: 5 μm and 1 μm in the original and enlarged images, respectively. (H) Quantification of synapsin I^+^ puncta in hCD45^+^ microglia week 8 of chimeras (n=120 microglia from 4 mice per group). Student’s t test, \*\*\**P* < 0.001, Data are presented as mean ± SEM. (I) A representative image of hTMEM119^+^ microglia and Neurobiotin^+^ recorded neurons in the hippocampus from a 2 to 3 months old DS microglia chimera. Arrows indicate Neurobiotin^+^ recorded neurons. Scale bar: 500 μm. (J) Representative traces of mEPSCs in CA1 hippocampal pyramidal neurons from Cont and DS chimeric mice. (K) Quantification of the frequency and amplitude of mEPSCs in Cont and DS chimeras (n=14-15 neurons from 4-5 mice per group). Student’s t test, \**P* < 0.05 and \*\**P* < 0.01. Data are presented as mean ± SEM.

We previously demonstrated that hiPSC-derived microglia were able to engulf synapses in chimeric mouse brains (Xu et al., 2020). Consistently, we found PSD95^+^ puncta within both Cont and DS microglia, indicating that they were able to prune synapses (Fig. 3B). Of note, the synaptic pruning function of Cont and DS microglia is developmentally regulated, increasing along with the age of animals from 4-week to 8-week to 3 months old. Further 3D reconstructed imaging analysis showed significantly more PSD95^+^ puncta in DS microglia than Cont microglia in GM at all three time points (Fig. 3B, 3D). In addition, to further examine microglial phagocytic function, we triple-stained hTMEM119, PSD95, and CD68, a marker for phagolysosomes. As shown in Fig. 3E, PSD95^+^ puncta were seen inside the CD68^+^ phagolysosomes of both hTMEM119^+^ Cont and DS microglia. DS microglia also showed enhanced phagocytosis, as indicated by a higher volume of CD68^+^ phagolysosome and more PSD95^+^ puncta in CD68^+^ phagolysosomes (Fig. 3F). Moreover, we double-stained human-specific CD45 (hCD45) with synapsin I (a presynaptic marker) and observed more synapsin I^+^ puncta within DS microglia, as compared to Cont microglia (Fig. 3G-H). In the WM, we also observed enhanced phagocytosis function as indicated by an increased volume of CD68^+^ phagolysosomes in DS microglia, compared to Cont microglia in chimeras at 8 weeks post transplantation (Fig. S3F-S3G). To further examine the dendritic spine density of hippocampal pyramidal neurons, Golgi staining was performed.

The results showed that DS chimeras had a lower dendritic spine density than Cont chimeras (Fig. S3H), which is consistent with the findings that DS microglia exhibit excessive pruning of PSD95^+^ synapses. Taken together, these results demonstrate that in the chimeric mouse brain model, DS microglia not only exhibit excessive synaptic pruning but also display less intricate morphology as compared to Cont microglia.

It is challenging to examine the impact of microglial synaptic pruning on synaptic neurotransmission in organoids, as neurons do not exhibit robust post-synaptic currents (PSCs) until after a long period of culturing (Xu et al., 2021). As such, our chimeric mouse model could facilitate the investigation on how DS microglia altered synaptic neurotransmission. We recorded miniature excitatory postsynaptic currents (mEPSCs) in hippocampal slices from Cont and DS microglial chimeric mice at 3 months post-transplantation. This time point was chosen because there was a wide distribution of transplanted Cont and DS microglia in the hippocampus, and a comparable level of chimerization by engrafted microglia was observed especially in the hippocampal CA1 region (Fig. 3A, 3I). Moreover, the recorded CA1 neurons labeled by neurobiotin through recording electrodes were surrounded by engrafted hTMEM119^+^ human microglia (Fig. 3I). We found that compared to Cont, DS microglial chimeric mice had mEPSCs with significantly reduced frequency and amplitude (Fig. 3J- K). These data indicate that excessive synaptic pruning of DS microglia may lead to impaired synaptic neurotransmission.

### Inhibiting the expression of IFNARs rescues the defective DS microglia development in chimeric mice

Trisomy 21 consistently activates type I IFN responses across different cell types, including immune cells (Araya et al., 2019; Sullivan et al., 2016; Waugh et al., 2019). The elevated type I IFN signaling is largely caused by increased gene dosage of type I IFNARs, as the genes encoding the subunits of IFNARs –*IFNAR1* and *IFNAR2* – are both located on Hsa21(Araya et al., 2019; Sullivan et al., 2016; Waugh et al., 2019). Type I IFN signaling regulates brain development and synaptic plasticity (Blank and Prinz, 2017; Ejlerskov et al., 2015; Hosseini et al., 2020). We examine the expression of type I IFN receptors in DS microglia (Fig. 4A). First, we performed qPCR using chimeric mouse brain tissues with human specific *IFNAR1* and *IFNAR2* primers. As shown in Fig. 4B, we found that DS microglial chimeras had higher expression of *IFNAR1* and *IFNAR2* gene transcripts than Cont microglial chimeras at 8 weeks and 4 months post-transplantation. Given that only engrafted human cells expressed the human specific *IFNAR* genes, this finding suggested that DS microglia expressed higher levels of *IFNAR1* and *IFNAR2* than Cont microglia. To further prove this, we performed flow cytometry targeting hCD45^+^ human microglia cells using 4 months old chimeric mouse brain tissues and analyzed the expression of IFNAR1 and IFNAR2 in these hCD45^+^ microglia. Consistently, DS chimeras showed higher levels of IFNAR1 and IFNAR2 expression than Cont chimeras (Fig. 4C-D). Therefore, these results demonstrate that compared to Cont microglia, there is an overexpression of IFNARs in DS microglia.

**Fig 4.**
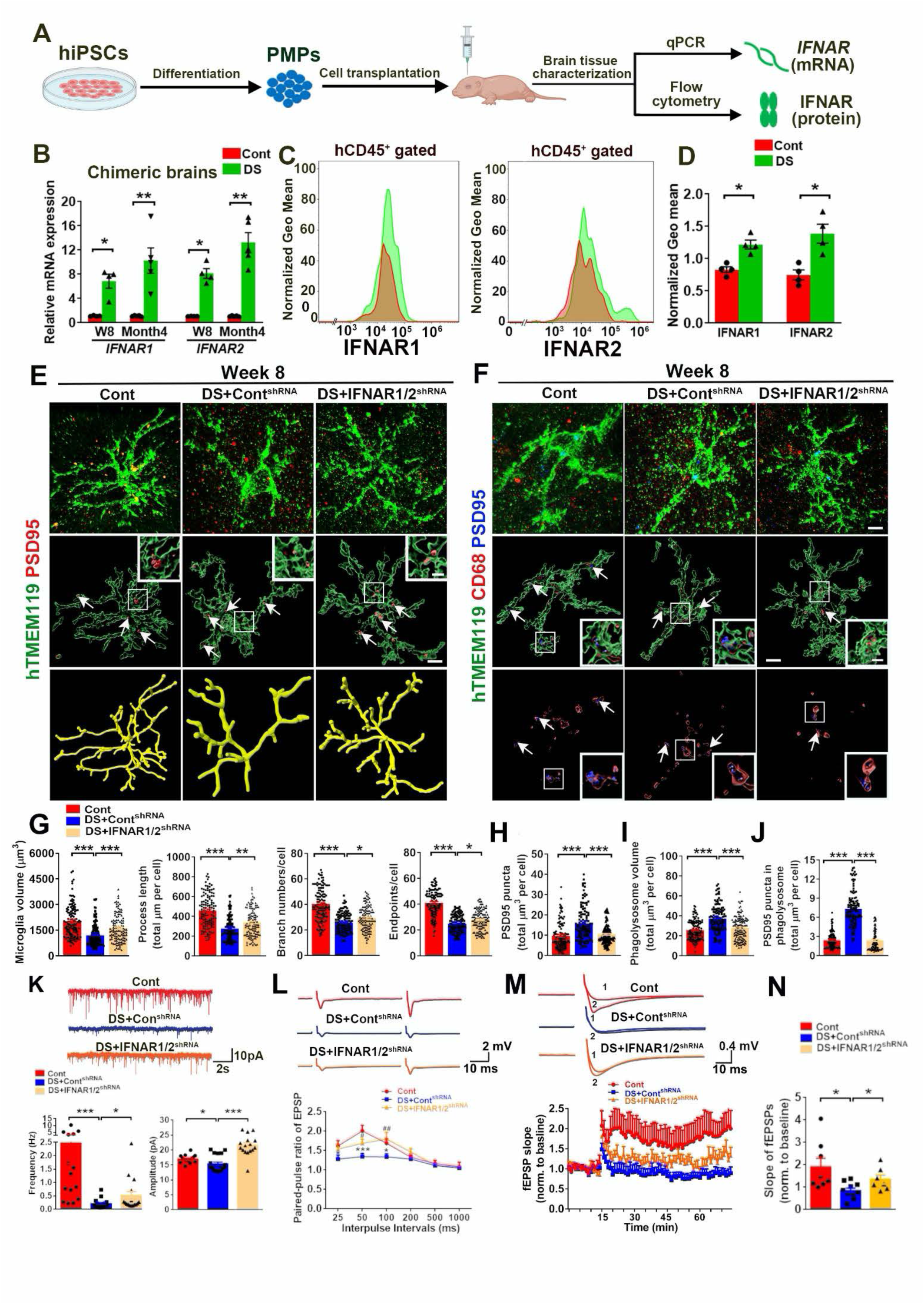
Knockdown of *IFNARs* rescues defective DS microglia in human-mouse microglial chimeras. (A) A schematic diagram showing the experimental design. This drawing created using BioRender.com. (B) qPCR analysis of *IFNAR1* and *IFNAR2* mRNA expression in chimeric mice at week 8 and month 4 (n=4-5 mice per group). Student’s t test, \**P* < 0.05 and \*\**P* < 0.01. Data are presented as mean ± SEM. (C, D) Flow cytometry analysis and quantification of IFNAR1 and IFNAR2 expression in 4 months old Cont and DS chimeric mice (n=4). Student’s t test, \**P* < 0.05. Data are presented as mean ± SEM. (E) Representative images of hTMEM119PSD95 staining in 8-week-old Cont, DS + Cont^shRNA^, and DS +IFNAR1/2^shRNA^ chimeras (n=115-136 microglia from 3-4 mice per group). Arrows indicate PSD95^+^ puncta. Scale bars:5 μm and 1 μm in the original and enlarged images, respectively. (F) Representative images of hTMEM119^+^CD68^+^ PSD95^+^ microglia in 8-week-old Cont, DS + Cont^shRNA^, and DS +IFNAR1/2^shRNA^ chimeras (n=113-135 microglia from 3-4 mice per group). Arrows indicate PSD95^+^ puncta in the CD68^+^ phagolysosome. Scale bars:5 μm and 1 μm in the original and enlarged images, respectively. (G) Quantification of microglial volume, process length, branch numbers, and endpoints (n=115-136 from 3-4 mice per group), One-way ANOVA test. \**P* < 0.05, \*\**P* < 0.01 and \*\*\**P* < 0.00. Data are presented as mean ± SEM. (H) Quantification of PSD95+ puncta in hTMEM119+microglia (n=115-133 microglia from 3-4 mice per group). One-way ANOVA test, \**P* < 0.05, \*\**P* < 0.01 and \*\*\**P* < 0.001. Data are presented as mean ± SEM. (I, J). Quantification of CD68+ phagolysosomes and PSD95+ puncta in CD68+ phagolysosomes (n=113- 136 microglia from 3-4 mice per group). One-way ANOVA test, \**P* < 0.05, \*\**P* < 0.01 and \*\*\**P* < 0.001. Data are presented as mean ± SEM. (K) Representative traces of mEPSCs in CA1 hippocampal pyramidal neurons from 2-3 months old Cont, DS+Cont^shRNA^ and DS+IFNAR1/2^shRNA^ chimeric mice. Quantification of the frequency and amplitude of mEPSCs in Cont and DS chimeras (n=15-18 neurons from 3 mice per group). Student’s t test, \**P* < 0.05 and \*\*\**P* < 0.001. Data are presented as mean ± SEM. (L) Representative traces and quantification of paired pulse ratio of EPSPs in 3-4 months old Cont, DS+Cont^shRNA^ and DS+IFNAR1/2^shRNA^ chimeras (n=9-12 slices from 3-4 mice per group). Asterisk represents Cont versus DS+ Cont^shRNA^, pound sign indicates DS+Cont^shRNA^ versus DS+IFNAR1/2^shRNA^. One-way ANOVA test, \**P* < 0.05, **p < 0.01 and ***p < 0.001. Data are presented as mean ± SEM. (M) Representative traces of baseline (1) and last 10 min (2) fEPSP after 4X 100 Hz LTP induction. Quantification of LTP after LTP induction in 3-4 months old Cont, DS+Cont^shRNA^ and DS+IFNAR1/2^shRNA^ chimeras (n=7-10 slices from 3-4 mice per group). Student’s t test, \**P* < 0.05. Data are presented as mean ± SEM. (N) Quantification of the last 10 min of fEPSP slope after LTP induction in 3-4 months old Cont, DS+Cont^shRNA^ and DS+IFNAR1/2^shRNA^ chimeras (n=7-10 slices from 3-4 mice per group). Student’s t test, \**P* < 0.05. Data are presented as mean ± SEM.

We then hypothesized that the overexpression of IFNARs in DS microglia might be responsible for their altered development and functions. To test this hypothesis, we took the RNAi knockdown approach. As observed phenotypes were highly consistent across the three DS hiPSC lines, we used two DS hiPSC lines (DS2 and the isogenic Tri-DS3) to express IFNAR1/2^shRNA^ or Cont^shRNA^. Overexpression of IFNARs in DS could be detected at iPSC stages, as indicated by the higher expression of *IFNAR1* and *IFNAR2* gene transcripts in DS hiPSCs compared to Cont hiPSCs (Fig. S4A). Transduction of DS hiPSCs with lentivirus carrying IFNAR1/2^shRNA^, but not the Cont^shRNA^, significantly inhibited the expression of *IFNAR1* and *IFNAR2* gene transcripts in DS hiPSCs (Fig. S4B). Then, we derived PMPs from DS hiPSCs treated with IFNAR1/2^shRNA^ (DS+IFNAR1/2^shRNA^) or Cont^shRNA^ (DS+Cont^shRNA^) and subsequently generated mouse chimeras by engrafting these PMPs. At 8 weeks post-transplantation, we analyzed the chimeric brains and found that the expression of *IFNAR1* and *IFNAR2* was downregulated in DS+IFNAR1/2^shRNA^ group when compared to DS+Cont^shRNA^ chimeras, as indicated by qPCR (Fig. S4C). Western blot analysis using an antibody specifically against human IFNAR2 confirmed the lower expression of IFNAR2 protein in DS+IFNAR1/2^shRNA^ microglia than DS+Cont^shRNA^ microglia (Fig. S4D). In both DS+IFNAR1/2^shRNA^ and DS+Cont^shRNA^ microglial chimeras, the vast majority of the hN^+^ cells were hTMEM119^+^, demonstrating that DS+IFNAR1/2^shRNA^ and DS+Cont^shRNA^ PMPs differentiated into microglia with similar efficiency (Fig. S4E-S4F). Interestingly, 3D skeleton analysis indicated that compared to DS+Cont^shRNA^ microglia, DS+IFNAR1/2^shRNA^ microglia in the GM have more ramified morphology as indicated by smaller cell volume, longer processes with an increased number of branches and endpoints (Fig. 4E, 4G). In the WM, we found that compared to DS+Cont^shRNA^, DS+ IFNAR1/2^shRNA^ microglia showed similar microglia volume, branch numbers, and endpoints but increased process length (Fig. S4G, S4I). Additionally, we found fewer PSD95^+^ puncta within DS+IFNAR1/2^shRNA^ microglia, a smaller total volume of CD68^+^ phagolysosome, and fewer PSD95^+^ puncta inside the CD68^+^ phagolysosomes in the DS+IFNAR1/2^shRNA^ microglia than in DS+Cont^shRNA^ microglia in the GM (Fig. 4F, 4H-J). We further explored the effects of inhibiting the overexpression of IFNARs in DS microglia on the hippocampal spine density. We observed that the spine density was higher in DS+IFNAR1/2^shRNA^ chimeric mice, as compared to DS+Cont^shRNA^ chimeras (Fig. S5A), suggesting that the dendritic spine density was rescued by inhibiting the overexpression of IFNAR in DS microglia. All these data suggest decreased synaptic pruning and phagocytic function of DS+IFNAR1/2^shRNA^ microglia at 8 weeks post-transplantation. Similarly, in the WM, the total volume of CD68^+^ phagolysosomes in DS+IFNAR1/2^shRNA^ microglia was smaller than DS+Cont^shRNA^ microglia, indicating a decrease in the phagocytic function (Fig. S4H, S4J). Taken together, these findings demonstrate that inhibiting the expression of IFNARs increases the ramification of DS microglia and rescues their synaptic pruning functions.

To explore the effects of inhibiting IFNARs in DS microglia on synaptic neurotransmission, we first recorded mEPSCs in hippocampal CA1 neurons from DS+Cont^shRNA^ and DS+IFNAR1/2^shRNA^ microglial chimeric mice at 3 months post-transplantation. There was a significant increase in mEPSC frequency and amplitude in DS+IFNAR1/2^shRNA^ microglial chimeric mice, compared to DS+Cont^shRNA^ group (Fig. 4K). These results indicate that overexpression of IFNARs in DS microglia may partly contribute to their effects on impairing synaptic neurotransmission. The changes in both frequency and amplitude of mEPSCs between DS and control microglial chimeras suggest that in addition to the observed postsynaptic alterations (Fig. 3B, 3D,3E-F and Fig. 4E-F, 4H-J), pre-synaptic mechanisms are involved. We thus further recorded pair-pulse ratio (PPR) to examine whether there were changes in synaptic release in the presynaptic sites. As shown in Fig. 4L, we observed pair-pulse facilitation (PPF) in the three groups. The PPR was decreased in DS+Cont^shRNA^ group compared with control microglia transplantation group. Inhibiting the IFNARs expression in DS microglia significantly enhanced the PPR, as indicated by the higher PPR at 50ms and 100ms interpulse intervals in DS+IFNAR1/2^shRNA^ group than in DS+Cont^shRNA^ group (Fig. 4L). To assess long-term synaptic plasticity, we induced long-term potentiation (LTP) using 4 trains of 100 Hz high-frequency stimulation (HFS). As shown in Fig. 4M, the enhancement of the fEPSP slope persisted for 60 min in Cont chimeric mice, but not in DS+Cont^shRNA^ chimeric mice. DS+IFNAR1/2^shRNA^ mice exhibited a significant enhancement of fEPSC slope than in DS+Cont^shRNA^ chimeric mice (Fig. 4N), suggesting that inhibiting the IFNARs expression could partially rescue the long-term synaptic plasticity. In addition, after PPF and LTP recording, we collected all the recorded hippocampal slices and performed histological analysis. All the slices similarly showed a wide distribution of xenografted, hTMEM119^+^ human microglia (Fig. S5B).

Altogether, these data strongly demonstrate that elevated synaptic pruning function of DS microglia results in impaired neurotransmission and synaptic plasticity in the hippocampus of the chimeric mice. Inhibiting the overexpression of IFNARs in DS microglia partially rescues the defects and improves synaptic functions.

### Single-cell RNA sequencing (scRNA-seq) analysis of DS microglial chimeric mouse brains following exposure to pathological tau

We next sought to explore whether our DS microglial mouse chimera could recapitulate the dystrophic microglia pathology found in AD and DSAD human brain tissues (Braak and Del Tredici, 2015; Jimenez et al., 2008; Martini et al., 2020; Meyer-Luehmann and Prinz, 2015; Sanchez-Mejias et al., 2016; Shahidehpour et al., 2020; Streit et al., 2009; Xue and Streit, 2011). Previous *in vitro* studies demonstrated that soluble p-tau in human AD brain tissue extracts induced microglial degeneration (Sanchez-Mejias et al., 2016). We thus prepared S1 soluble protein fractions from DS human brain tissue with abundant tau pathology (Tangle stage 6, Table S4) qualifying for AD, as well as from age- and sex-matched control brain tissues, following the method described previously (Clavaguera et al., 2013; Clavaguera et al., 2009; Jimenez et al., 2014; Sanchez-Mejias et al., 2016). Western blot analysis showed that p-tau was not detectable in control brain tissue-derived soluble fractions by PHF-1 antibody that recognizes p-tau at 396/404 sites (Otvos et al., 1994). In contrast, p-tau was highly abundant in DSAD brain tissue-derived soluble tau fractions (Fig. S6A). In addition, both control brain and DSAD brain tissue-derived soluble fractions had undetectable levels of Aβ content, whereas Aβ could be detected in the homogenate of DSAD cortical tissue (Fig. S6A). We also measured the concentration of total tau in soluble S1 fractions by ELISA and found that both groups had a similar level of total tau (Table S5). Chimeric mice engrafted with microglia derived from two DS lines (DS2 and Tri-DS3) further received injection of either Cont or DSAD at the age of 8 to 10 weeks, as previously described (Boluda et al., 2015; Iba et al., 2013; Peeraer et al., 2015). At two weeks post- injection, AT8^+^ p-tau was found to be distributed widely in the mouse brains that received DSAD tau, which is consistent with previous reports (Boluda et al., 2015; Nimmo et al., 2020). In the control tau injection group, very few AT8^+^ p-tau was detected (Fig. S6B). To characterize the responses of DS microglia to p-tau, we examined phagocytosis of p-tau by double-staining p-tau with human microglia at 1-2 months post tau injection. As shown in Fig.5 A, 5B and Fig. S6C, S6D, in both GM and WM, hTMEM119^+^ DS microglia engulfed a large number of AT8^+^ or PHF-1^+^ p-tau in the DSAD Tau injection group. In the Cont tau injection group, few p-tau was seen in DS microglia. These results demonstrate the successful injection of soluble tau and robust responses of DS microglia to p-tau in chimeric mice.

**Fig 5.**
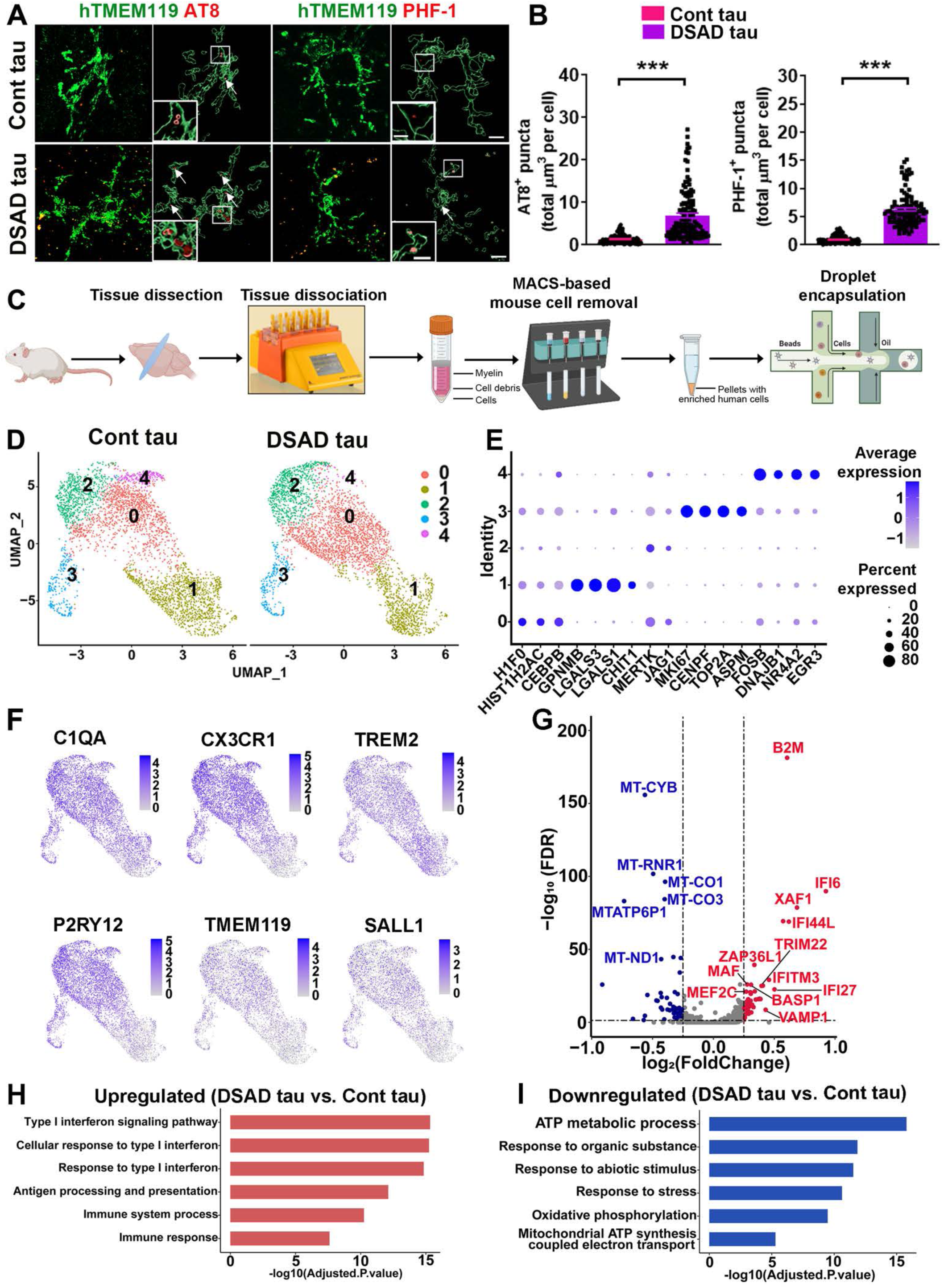
scRNA-seq analysis of DS microglial chimeric brains receiving injection of Cont or pathological DSAD tau. (A) Representative images of hTMEM119, AT8, and PHF-1 staining in 4 to 5-month-old chimeric mice receiving injection of Cont or DSAD tau at the age of 8 weeks. Arrows indicate AT8^+^ or PHF-1^+^ p-tau. Scale bars: 7 μm and 3 μm in the original and enlarged images, respectively. (B) Quantification of AT8^+^ and PHF-1^+^ p-tau in microglia (n=110-127 from 3-4 mice per group). Student’s t test, \*\*\**P* < 0.001. Data are presented as mean ± SEM. (C) A schematic diagram showing the design of the scRNA-seq experiment. DS microglia were isolated from 4 months old chimeric mice that received an injection of Cont or DSAD tau at 8 weeks. This drawing created using BioRender.com. (D) A UMAP plot showing independent subclusters (clusters 0-4) from Cont tau and DSAD tau groups. (E) A dot plot showing the representative conserved markers from each subcluster. (F) UMAP plots with dots (representing cells) colored by the expression levels of human microglial genes. (G) A volcano plot illustrating downregulated (blue) and upregulated (red) DEGs in DS microglia. (H, I) GO enrichment analyses of the upregulated and downregulated DEGs in DS microglia.

To determine the responses of DS microglia to p-tau at the transcriptomic level, we performed scRNA-seq. At two months following tau injection into the chimeric mice engrafted with DS2 and Tri- DS3 hiPSC-derived microglia, brain regions, including the cerebral cortex, hippocampus, corpus callosum, and olfactory bulb, were collected for scRNA-seq. To capture ample numbers of DS iPSC-derived microglia for scRNA-seq, we enriched DS microglia by using a magnetic cell sorting-based mouse cell depletion kit, as previously described (Hasselmann et al., 2019b). After cell sorting, the flow through was collected and centrifuged, and then the cell suspension was directly subjected to droplet-based scRNA-seq isolation (Fig. 5C). A total of 7,790 human microglial cells were recovered after quality control, of which 3,919 DS microglia were from the Cont tau group and 3,871 DS microglia were from the DSAD tau group. The median gene counts were 1,041 and 1,158 per cell for Cont tau and DSAD tau groups, respectively (Fig. S6E). We identified 6 clusters, in which cluster 5 contained a very small cell number (< 0.4% of cells), expressed hematopoietic progenitor cell marker *CD34,* rather than microglial markers (Fig. S6E), and thus was not included for further analysis. UMAP (Uniform Manifold Approximation and Projection) and clustering of DS microglia from Cont tau and DSAD tau groups demonstrated the overall similarities in clustering of the two groups (0-4) (Fig. 5D). Clusters can be distinguished by their expression of enriched markers (Table S6), which included 5 subclusters: cluster 0 (*H1F0, HIST1H2AC,* and *CEBPB*), cluster 1 (*GPNMB, LGALS3, LGALS1,* and *CHIT1*), cluster 2 (*MERTK* and *JAG1*), cluster 3 (*MKI67, CENPF, TOP2A,* and *ASPM*), and cluster 4 (*FOSB, DNAJB1, NR4A2,* and *EGR3*) (Fig. 5E). All clusters highly expressed microglial markers, including *C1QA, CX3CR1, TREM2, TMEM119, P2RY12,* and *SALL1* (Fig. 5F). Using a cutoff of ± 0.25 log fold change and an FDR of < 0.05, we identified 122 DEGs that included 56 upregulated and 66 downregulated genes (Fig. 5G, Table S7) in DSAD tau group, as compared with the Cont tau group. To explore the function of these DEGs, we performed Gene Ontology (GO) and KEGG enrichment analyses by using g: Profiler. Interestingly, many significantly enriched DEGs that were upregulated in DSAD tau group were associated with the type I IFN signaling pathway (e.g. *IFI6, XAF1, IFI44L, IFI27,* and *IFITM3*) (Ioannidis et al., 2012) and cell senescence pathway (e.g. *B2M, ZFP36L1,* and *XAF1*) (Heo et al., 2016; Loh et al., 2020; Smith et al., 2015) (Fig. 5H, S7A). In subclusters (0 and 3), we also found upregulated type I IFN signaling (Fig. S7B). Furthermore, a considerable amount of downregulated DEGs in the DSAD tau group are involved in mitochondrial functions and ATP metabolic processes (e.g., *LDHA, ALDOA, MT-ATP6,* and *HSPA1B*), suggesting dysfunctional mitochondria and reduced production of ATP. This was also seen in subclusters (e.g., 0 and 1) (Fig. 5I, S7C). Cluster 4 accounted for a very small proportion in the DSAD tau group (0.64%), whereas in the Cont tau group, cluster 4 accounted for 6.66% (Fig.S6E). Thus, a small number of downregulated DEGs in the cluster 4 of DSAD tau group were found, which were not enriched for any significant pathways in our GO analysis of cluster 4 (Fig.S7C).

To explore the heterogeneity of these DS microglia, we compared the conserved gene markers that define each cluster with both genes that define disease-associated microglia (DAM; Table S8) (Butovsky and Weiner, 2018; Chen and Colonna, 2021; Deczkowska et al., 2018; Keren-Shaul et al., 2017; Krasemann et al., 2017; Sobue et al., 2021) and cellular senescence genes (Table S9) (Avelar et al., 2020; Zhao et al., 2016). When comparing the expression of DAM genes with the conserved markers of each cluster, we found that DAM genes were expressed by all clusters. Notably, the conserved markers of cluster 1 have the highest number of genes that overlapped with DAM genes (Fig. 6A). We found that 21 of the total upregulated DEGs detected from each cluster in DSAD tau group were DAM genes. The dot plot shown in Fig. 6B demonstrated that most of the identified DAM DEGs had higher expression levels in all clusters of DSAD tau group than those of Cont tau group. For cellular senescence genes, we found that many of these genes were expressed in all the clusters, while cluster 3 expressed the highest number of senescence genes (Fig. 6C). We found that 8 of the total upregulated DEGs detected from each cluster in DSAD tau group were senescence genes. Similarly, the dot plot shown in Fig. 6D demonstrated that most of the identified senescence DEGs had higher expression levels in all clusters of DSAD tau group than those of Cont tau group. Notably, all the 8 senescence DEGs were upregulated in clusters 1 and 3 in DSAD tau group.

**Fig 6.**
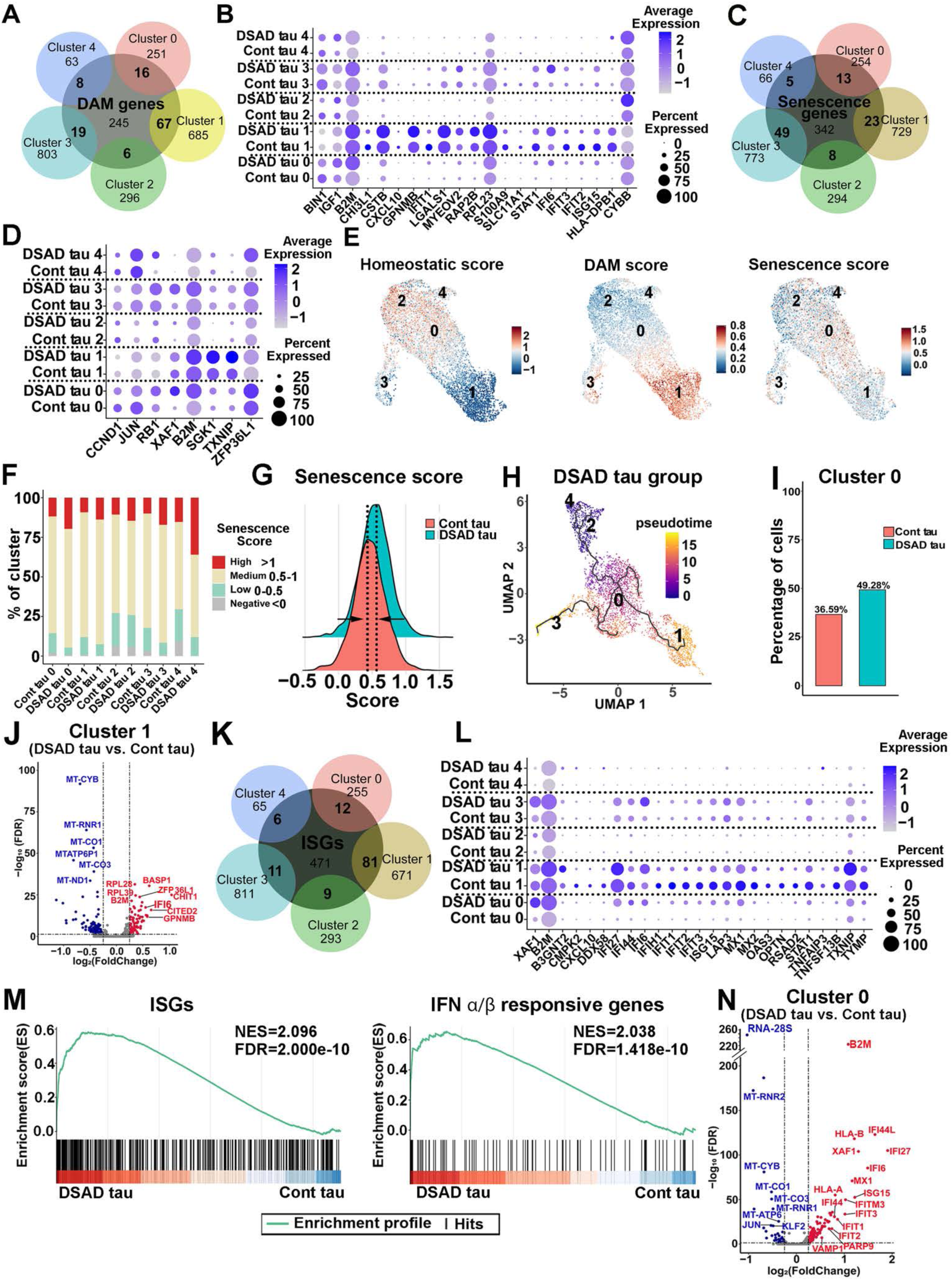
Pathological tau induces DAM, senescence, and ISG signatures in DS microglia. (A) A Venn diagram showing the overlap between DAM genes and the gene markers defined from each cluster. (B) A dot plot representing the expression of DAM DEGs in each cluster from the Cont and DSAD tau groups. (C) A Venn diagram showing the overlap between senescence genes and gene markers defined from each cluster. (D) Dot plot showing the DEGs expression of the senescence genes signature in every subcluster from DSAD tau and Cont tau groups. (E) Feature plots showing the scoring of homeostatic, DAM, and senescence signatures in DS microglia. (F) Bar plots of the percentage of cells with a negative (< 0), low (0-0.5), medium (0.5-1), or high (> 1) senescence score in the custom senescence signature in each cluster. (G) A ridge plot showing the senescence score in DSAD tau and Cont tau groups. (H) UMAP representation of the trajectory of DS microglia in response to DSAD tau. Cells are colored by pseudotime. (I) Cell ratios of cluster 0 in the Cont and DSAD tau groups. (J) A Volcano plot illustrating the downregulated (blue) and upregulated (red) DEGs in cluster 1. (K) A Venn diagram showing the overlap between ISGs and the gene markers defined from each cluster. (L) A Dot plot displaying the ISG DEGs in each cluster from the Cont tau and DSAD tau groups. (M) GSEA plots showing enrichment of ISGs and IFNα/β responsive genes in DSAD tau and Cont tau groups (NES: normalized enrichment score, FDR: false discovery rate). (N) A Volcano plot showing the downregulated (blue) and upregulated (red) DEGs in cluster 0.

To further explore the association between DAM and senescent phenotype in DS microglia, we then probed the dataset with homeostatic genes (*TMEM119, P2RY12, P2RY13, CX3CR1, SELPLG, and BIN1)*, DAM genes (Table S8), and custom senescence signature genes (*ATM, AXL, B2M, ZFP36L1, XAF1, CDKN2A, CDKN1A, CDKN2D, CASP8, IL1B, GLB1, and SERPINE1).* As shown in Fig. 6E, cluster 2 expressed high levels of homeostatic genes and thus was annotated as homeostatic microglia, while cluster 1 was annotated as DAM. Very interestingly, significant numbers of cells displaying senescence signature were seen in all clusters, particularly in cluster 0. As compared to Cont tau group, more cells in each subcluster in the DSAD tau group displayed greater association with senescence signature (Fig. 6F). Moreover, DSAD tau group overall showed a shift towards a higher senescence signature (Fig. 6G). Further trajectory analysis within DSAD tau group revealed a phenotypical change of DS microglia from cluster 2 (homeostatic) toward the cluster 3 (senescent state) and cluster 1 (DAM state), passing through cluster 0, an intermediate stage (Fig. 6H). Notably, cluster 0 was enriched in DSAD group (Fig. 6I). A recent study reported that DAM exhibited senescent phenotypes (Hu et al., 2021). Next, we analyzed DEGs within cluster 1, as cluster 1 had the highest DAM score. As shown in Fig. 6J (Table S11), the volcano plot showed that *B2M,* a pro-aging factor in neurodegenerative disease (Smith et al., 2015), was one of the top upregulated DEGs in the DSAD tau group. Moreover, *ZAP36L1*, whose expression was shown to promote cellular senescence (Galloway et al., 2016; Loh et al., 2020), was expressed at a much higher level in the cluster 1 of DSAD tau group than that of Cont tau group (Fig. 6H). In addition, the expression of various mitochondria genes (e.g. *MT-CYB, MT-RNR1, MT-CO1,* and *MT-CO3*) (Lunnon et al., 2017) was decreased in cluster 1 of DSAD tau group, which also suggested cellular senescence in DS microglia, because senescent cells often show impaired cell metabolism (Lopez-Otin et al., 2013; Vasileiou et al., 2019). Taken together, our scRNA-seq results depict that DSAD tau promotes overall senescence of DS microglia in all clusters and induces a DAM-like population of microglial cluster, specifically cluster 1. This DAM cluster 1 in DSAD tau group also shows enhanced cellular senescence.

Since we identified several elevated type I IFN signaling genes in DSAD tau group by analyzing overall DEGs (Fig. 5G), we further examined the expression of interferon-stimulated genes (ISGs, Table S10) (Hubel et al., 2019; Rusinova et al., 2013). The Venn diagram shown in Fig. 6K demonstrated that ISGs were expressed by all subclusters. The DAM cluster 1 expressed the highest number of ISGs. 25 of the total upregulated DEGs detected from each cluster in DSAD tau group were ISGs (Fig. 6L). The pathway enrichment analysis also indicated increased activation of type I IFN pathway in the DSAD tau group when compared to the Cont tau group (Fig. S7B). We performed gene set enrichment analysis (GSEA) and interestingly identified notable enrichment of IFN stimulated genes and IFNα/β responsive genes in the DSAD tau group (Fig. 6M). As cluster 0 was an intermediate state (Fig. 6H), this cluster likely represented early responses of DS microglia to DSAD tau. Moreover, given the fact that cluster 0 was enriched in DSAD tau group and showed a higher senescence score (Fig. 6E), we further analyzed DEGs within cluster 0 between DSAD tau and Cont tau groups. Intriguingly, the volcano plot indicated that many type I IFN signaling-related genes (e.g., *IFI44L, XAF1, IFI27, IFI6, IFI44, IFIT3,* and *IFITM3*) were significantly upregulated in cluster 0 of DSAD tau group (Fig. 6N, Table S11). Therefore, these data demonstrate that increased cellular senescence of DS microglia subclusters in DSAD tau group largely overlaps with the elevated type 1 IFN signaling, suggesting that the pathological tau might work through activating type 1 IFN signaling to induce senescence of DS microglia.

### DS microglial senescence induced by pathological tau can be rescued via inhibiting microglial *IFNARs*

To experimentally validate the senescent phenotype of DS microglia as indicated by scRNA-seq data, we first compared the morphology of hN^+^ DS microglia in Cont tau and DSAD tau groups, because dystrophic morphology is a prominent characteristic of senescent microglia (Ransohoff, 2016; Shahidehpour et al., 2020; Streit et al., 2009; Streit et al., 2004; Xue and Streit, 2011). By double-staining Iba-1 and hN, we found that the percentage of hN^+^/Iba-1^+^ cells among total Iba-1^+^ cells was similar between the two groups (Fig.7A, 7C). hN^+^/Iba-1^+^ DS microglia in DSAD tau group, but not in the Cont tau group, displayed dystrophic morphology, such as beading with shortened processes and fragmentation (Fig. 7A). Quantitative analysis showed shortened process length, enlarged soma size, and an increased ratio between soma size and process length in the DSAD tau group (Fig. 7D). These morphological changes have been observed in dystrophic microglia in human DSAD brain tissue (Flores-Aguilar et al., 2020; Martini et al., 2020). Concomitantly, we found that in the same chimeric mouse brains that received DSAD tau injection, the hN-negative/Iba-1^+^ host mouse microglia displayed hypertrophic morphology with shortened processes, decreased branch numbers, and endpoints, rather than dystrophic morphology (Fig. S8A-S8B). This suggests that in response to pathological tau, mouse microglia may have reactions that are different from human microglia. We next stained chimeric mouse brain tissue with hCD45 and ferritin, expression of which has been strongly linked to microglial senescence (Lopes et al., 2008; Simmons et al., 2007; Verina et al., 2011). We found that there was an increased number of ferritin^+^/hCD45^+^ DS microglia in the DSAD tau group compared to the Cont tau group (Fig. 7B, 7E). Furthermore, scRNA-seq analysis showed that multiple inflammatory cytokines and chemokines, such as *IL1B*, *CCL2*, and *CCL4*, were expressed at lower levels in DS microglia in the DSAD tau group than those in the Cont tau group (Fig. 7F). Consistently, qPCR analysis using human specific primers for *IL1B* and *TNFA* that respectively encode IL1β and TNFα also showed decreased expression of these inflammatory cytokines in DS microglia in DSAD tau group (Fig. 7G). Taken together, these results demonstrate that pathological tau induces cellular senescence of DS human microglia rather than causing massive activation of human microglia in the chimeric mouse brain.

**Fig 7.**
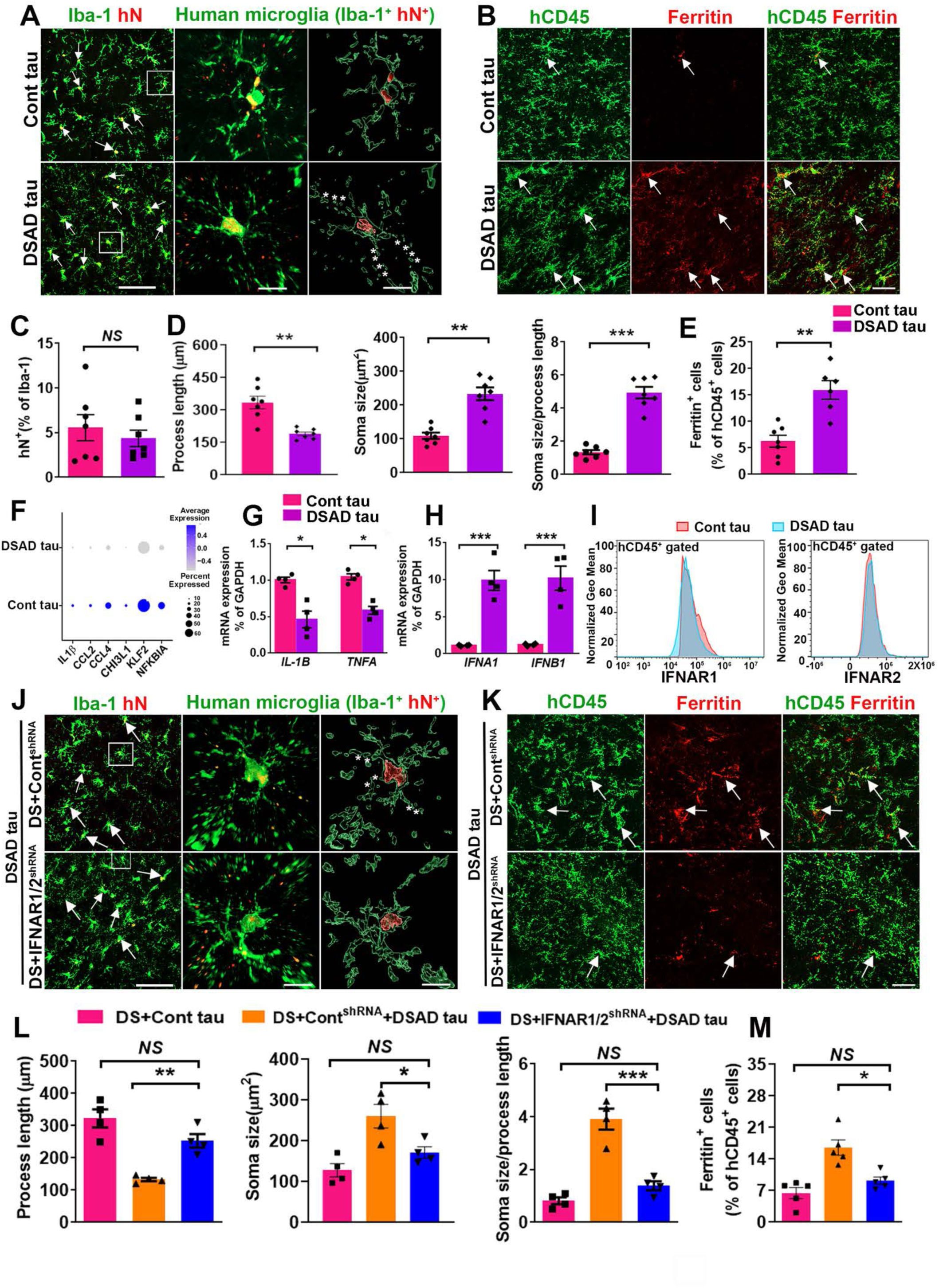
Knockdown of IFNARs rescues pathological tau-induced senescence in DS microglia. (A) Representative images of Iba^+^/hN^+^ human microglia in Cont tau and DSAD tau groups. Arrows indicate Iba^+^/hN^+^ human microglia. Asterisks indicate fragmented processes. Scale bars: 50 μm and 10 μm. (B) Representative images showing colocalization of hCD45^+^ and Ferritin^+^ staining in Cont and DSAD tau groups. Arrows indicate Ferritin^+^ and/or hCD45^+^ staining. Scale bar: 50 μm. (C) Quantification of the percentage of hN^+^ in Iba-1^+^ cells in Cont and DSAD tau groups (n=7 mice/group). Student’s t test, *NS*, not significant. Data are presented as mean ± SEM. (D) Quantification of the process length, soma size, soma size/process length cells (n=7 mice per group). Student’s t test, \**P* < 0.05, \*\**P* < 0.01, \*\*\**P* < 0.001, *NS*, not significant. Data are presented as mean ± SEM. (E) Quantification of the percentage of Ferritin in hCD45^+^ cells (n=6-7 mice per group). Student’s t test, \*\**P* < 0.01. Data are presented as mean ± SEM. (F) A Dot plot representing the expression of the inflammation-related genes *IL1β*, *CCL2*, *CCL4, CHI3L1, KLF2, NFKBIA* identified from scRNA-seq. (G) qPCR analysis of *IL-1B and TNFA* mRNA expression (n=4 mice per group). Student’s t test, \**P* < 0.05. Data represented as mean ± SEM. (H) qPCR analysis of *IFNA1* and *IFNB1* mRNA expression (n=4 mice per group). Student’s t test, \*\*\**P* < 0.001. Data represented as mean ± SEM. (I) Flow cytometry analysis showing the expression of IFNAR1 and IFNAR2 in Cont tau and DSAD tau group (n=2 mice per group). (J) Representative images of human microglia (Iba^+^hN^+^) in DS+Cont^shRNA^+DSAD tau and DS+IFNAR1/2^shRNA^+DSAD tau chimeric mice. Arrows indicate Iba^+^/hN^+^ human microglia. Asterisks indicate fragmented processes. Scale bar: 50 μm and 10 μm. (K) Representative image showing colocalization of hCD45^+^ and Ferritin^+^ staining in DS+Cont^shRNA^+DSAD tau and DS+IFNAR1/2^shRNA^+DSAD tau chimeric mice. Arrows indicate Ferritin^+^ and/or hCD45^+^ staining. Scale bar: 50 μm. (L) Quantification of the process length, soma size, soma size/process length (n=4 mice per group). One-way ANOVA test, \*\**P* < 0.01 and \*\*\**P* < 0.001, *NS*, not significant. Data are presented as mean ± SEM. (M) Quantification of the percentage of Ferritin in hCD45^+^ cells (n=5 mice per group). One-way ANOVA test, \**P* < 0.05, *NS*, not significant. Data are presented as mean ± SEM.

These findings also indicate that our chimeric mouse brain model can faithfully recapitulate microglial pathology that is seen in human AD and DSAD brain tissues (Boluda et al., 2015; Flores-Aguilar et al., 2020; Martini et al., 2020; Streit et al., 2009; Streit et al., 2020; Streit et al., 2004; Xue and Streit, 2011). To further explore the impact of pathological tau on Cont microglia, we injected either Cont or DSAD tau into control microglial chimeric mice at 8 weeks post-transplantation and then stained the brain tissues with hCD45 and ferritin at two months and six months after tau injection. As shown in Fig. S8E, we found no ferritin^+^/hCD45^+^ control microglia in the Cont tau group and very few ferritin^+^/hCD45^+^ control microglia in the DSAD tau group at two months post tau injection. Interestingly, there were many more ferritin^+^/hCD45^+^ control microglia at six months post DSAD tau injection, compared with that at 2 months after DSAD tau injection (Fig. S8E). In line with previous reports (Romero-Molina et al., 2018; Sanchez-Mejias et al., 2016), soluble p-tau is highly toxic for human microglia, including both control and DS microglia. Furthermore, in DS microglial chimeric mice, we observed ferritin^+^/hCD45^+^ DS microglia as early as 1 to 2 months post-tau injection, suggesting that DS trisomy genotypes significantly contribute to the phenotypes induced by p-tau treatment.

ScRNA-seq data prompted us to hypothesize that elevated type I IFN signaling could mediate the pathological tau-induced senescence in DS microglia. We then assessed the expression of *IFNA1* and *IFNB1* genes, which respectively encodes IFNα and IFNβ, the ligands of IFNARs. We found that the *IFNA1* and *IFNB1* gene transcripts were significantly upregulated in the DSAD tau group compared to the Cont group (Fig. 7H). We further evaluated the expression of *IFNARs* using human specific *IFNAR1* and *IFNAR2* primers and found no differences in the expression of *IFNARs* between Cont Tau and DSAD Tau groups (Fig. S8C). In addition, flow cytometry analysis also showed that the expression of IFNARs was similar in both groups (Fig. 7I). These results suggest that pathological tau increases the production of type I IFNs but is not able to further increase the already overexpressed IFNARs in DS microglia (relative to Cont microglia as shown in Fig.7I, S8C). This is consistent with the concept that IFN signaling pathways are usually upregulated by increasing the production of ligands rather than IFN receptors (Platanias, 2005).

Previous studies in mice on type I IFN signaling yielded contradictory results, as some showed that blocking type I IFN signaling could limit disease progression in various models of AD (Baruch et al., 2014; Taylor et al., 2014; Taylor et al., 2018), while others demonstrated that a lack of type I IFN signaling caused spontaneous neurodegeneration (Ejlerskov et al., 2015). These conflicting observations underscore the importance of distinguishing cell type-specific roles of IFNAR in the disease. In addition, given IFNAR1 and *2* are triplicated in DS and overexpressed in DS microglia (Fig. 4B, 4C), we proposed that cell-type specific inhibition of IFNARs in DS microglia might prevent their senescence in response to pathological tau. To test this, we took the RNAi knockdown approach again and used the two DS hiPSC lines (DS2 and Tri-DS3) expressing IFNAR1/2^shRNA^ or Cont^shRNA^. We found that the expression of *IFNA1* and *IFNB1* remained higher in the DS IFNAR1/2^shRNA^ + DSAD tau group compared to the DS IFNAR1/2^shRNA^ + Cont AD tau group (Fig. S8D). Then, we characterized the morphology of DS microglia with Iba1 and hN staining. As compared to DS microglia in the DS Cont^shRNA^+DSAD tau, DS microglia in the DS IFNAR1/2^shRNA^+DSAD tau showed less fragmented processes (Fig. 7L), longer process length, smaller soma size, and decreased soma size/process length ratio (Fig. 7L). DS microglia in the DS IFNAR1/2^shRNA^+DSAD tau group showed similar process length and soma size in comparison with those in the DS+Cont tau group (Fig. 7L). Moreover, there were fewer Ferritin^+^/hCD45^+^ microglia in the DS IFNAR1/2^shRNA^+DSAD tau group than the DS Cont^shRNA^+DSAD tau group (Fig. 7K, 7M). Taken together, these results indicate that inhibiting the expression of IFNARs in DS microglia rescues their senescent phenotypes following DSAD tau injection.

## Discussion

We have previously demonstrated that the complementary combination of human iPSC-derived cerebral organoid and chimeric mouse brain systems can precisely model DS-related phenotypes and help to understand the underlying molecular mechanisms (Xu et al., 2019). In this study, using our new microglia-containing cerebral organoid and human-mouse chimeric brain models, we reveal that DS microglia exhibit abnormal alterations at morphological (e.g. shortened processes and fewer branches) and functional (e.g. enhanced synaptic pruning and phagocytic functions) levels during brain development. Moreover, we discover that human brain tissue-derived soluble pathological tau promotes cellular senescence of DS microglia in chimeric mouse brains. Importantly, inhibiting the expression of *IFNARs* in DS microglia can not only reverse the morphological and functional alterations of DS microglia but also prevent senescence of DS microglia in response to pathological tau.

Cerebral organoids have proven to be valuable in modeling developmental defects in various types of neural cells. We find that DS microglia show enhanced synaptic pruning function in cerebral organoids, consistent with previous studies in a mouse model of DS and DS human brain tissue (Flores-Aguilar et al., 2020; Pinto et al., 2020). Previous studies also report that DS microglia show morphological signs of activation (Flores-Aguilar et al., 2020; Pinto et al., 2020). However, microglia in organoid models (Abud et al., 2017; Ormel et al., 2018) do not develop the highly ramified morphology that is comparable to the morphology in the brain. Similarly, in our cerebral organoids, both DS vs. Cont microglia did not exhibit highly branched and ramified morphology. Thus, we did not observe any significant morphological differences between DS vs. Cont microglia. We think that this might be due to the intrinsic limitations of organoids, such as the continued immaturity of neural cells – an issue noted with all current organoid technologies. In the early developing brain, microglia mature to a ramified phenotype along with the maturation of other neural populations, and ramification of microglia is largely controlled by neuron and macroglia (astroglia and oligodendroglia)-derived factors, such as IL34 and CSF1, and by neuron–microglia interactions (Kierdorf et al., 2013; Mosser et al., 2017; Schulz et al., 2012). Immature neural cells in the organoids may not be able to provide an environment that is enough to support microglial ramification. Thus, the current form of the cerebral organoids is not able to fully recapitulate DS microglia phenotypes. As such, we further model the defects of DS microglia using our *in vivo* human microglial chimeric mouse model, in which engrafted human microglia actively interact with neurons in a brain development-dependent manner (Xu et al., 2020). We find that the human microglia chimeric mice not only faithfully recapitulate the defective functions of DS microglia but also their morphological changes. Very interestingly, as compared to control microglial chimeras, DS microglial chimeras showed mEPSCs with decreased amplitude and frequency. These findings corroborate the similar observations in Dp (16) mouse model of DS, suggesting that over-pruning of synapses by DS microglia inhibits synaptic neurotransmission and contributes to the cognitive deficits in DS.

Our knowledge on the pathophysiology of DS microglia is primarily gained from studies in transgenic mouse models of DS and postmortem human DS brain tissues. However, these strategies have significant limitations in studying underlying molecular mechanisms, because functional human DS brain tissues are rarely available, and the mouse models demonstrate incomplete/inaccurate expression of Hsa21 genes. Hsa21 contains more than 500 genes (Moyer et al., 2020), whereas, for example, in Dp(16) mouse model that recently was used to study DS microglia (Pinto et al., 2020), only 113 genes are triplicated. Owing to the much better genetic tractability of the hiPSC models as compared to the mouse models of DS (Moyer et al., 2020), we previously demonstrated the application of DS hiPSC models in studying the genetic underpinning of DS-related phenotypes (Chen et al., 2014; Xu et al., 2019). In this study, we focus on type I IFN signaling, because the roles of type I IFN signaling are not only limited to antiviral and immunomodulation, but are also involved in regulating homeostatic processes in the CNS (Blank and Prinz, 2017). It is known that under healthy conditions, baseline levels of constant type I IFN signaling play critical roles in brain development and functions, such as synaptic plasticity (Blank and Prinz, 2017; Ejlerskov et al., 2015; Hosseini et al., 2020). Moreover, trisomy 21 consistently activates type I IFN responses across different cell types, including immune cells (Araya et al., 2019; Sullivan et al., 2016; Waugh et al., 2019). The elevated type I IFN signaling is mainly caused by increased gene dosage of type I IFN receptors, as the genes encoding the subunits of type I IFN receptor, *IFNAR1,* and *IFNAR2*, are both located on Hsa21(Araya et al., 2019; Sullivan et al., 2016; Waugh et al., 2019). Our results show that DS microglia express higher levels of IFNAR1 and IFNAR2 than Cont microglia at both transcript and protein levels during development. shRNA-mediated inhibition of the *IFNAR* expression improves the morphological complexity of DS microglia, corrects its synaptic pruning functions, and rescues the hippocampal synaptic neurotransmission and plasticity, suggesting that type I IFN signaling could be targeted to improve microglial functions in DS brain development.

Previous studies on microglial responses in AD have largely relied on APP-based rodent models. Those studies demonstrate that microglial dysfunction in AD is primarily associated with overactivation, neuroinflammation, and cytotoxicity of microglia (Hansen et al., 2018; Heppner et al., 2015; Sarlus and Heneka, 2017). Similarly, exposing xenografted human microglia to pathological Aβ in chimeric mouse brain models also largely induces activation of the microglia, transitioning from homeostatic state to cytokine response and activated response states (Hasselmann et al., 2019a; Mancuso et al., 2019). Although the amyloid cascade/neuroinflammation theory of AD has been widely accepted, several caveats have arisen. The microglial responses observed in the hippocampus of APP-based mouse models differ from those observed in AD and DSAD patients (Braak and Del Tredici, 2015; Jimenez et al., 2008; Meyer-Luehmann and Prinz, 2015; Sanchez-Mejias et al., 2016; Shahidehpour et al., 2020; Streit et al., 2009; Xue and Streit, 2011). More importantly, non-steroidal anti-inflammatory drugs have demonstrated no clear ability to delay onset or reverse cognitive dysfunction in AD patients (Arvanitakis et al., 2008; Green et al., 2009; Group et al., 2007; Group et al., 2008). Based on postmortem neuropathological examinations of human AD and DSAD brain tissues, a paradigm-shifting tau/microglial senescence hypothesis has emerged, in which human microglia respond to pathological tau by exhibiting accelerated senescence and dystrophic phenotypes, rather than massive activation as previously thought (Guerrero et al., 2021; Navarro et al., 2018; Sanchez-Mejias et al., 2016; Streit et al., 2020; Streit et al., 2004). Rodent models are not able to faithfully recapitulate phenotypes of senescent human microglia. Senescent rodent microglia only exhibit small decreases in the complexity of their processes, but in humans, decreased complexity of microglia is much more advanced and produces dystrophic processes, which include cytoplasmic fragmentation (Streit et al., 2020; Xu et al., 2008). In addition, transcriptomic studies show that human and rodent microglia age differently under normal and diseased conditions (Friedman et al., 2018; Galatro et al., 2017). As such, the supporting evidence for tau/microglial senescence hypothesis from live, functional brain tissue that contains human microglia as well as knowledge on underlying molecular mechanism have so far been out of reach. Both clinical manifestations and biological evidence demonstrate accelerated aging of the brain and the immune system in individuals with DS (Cuadrado and Barrena, 1996; Horvath et al., 2015; Lott and Head, 2005; Teipel and Hampel, 2006). Our recent work has also shown higher numbers of dystrophic/senescent microglia in the brains of people with DS (Martini et al., 2020). Moreover, in human microglial chimeric mouse brains, maturation and aging of donor-derived human microglia appears to be accelerated in the much faster-developing mouse brain relative to human brain (Hasselmann et al., 2019a; Jiang et al., 2020; Mancuso et al., 2019; Svoboda et al., 2019; Xu et al., 2020). In support of this, our scRNA-seq analysis demonstrates that DS microglia show expression of senescent genes at 4 months post-transplantation even in the absence of pathological tau (Fig. 6D, 6F). Therefore, the combination of DS and hiPSC microglial chimeric mouse model presents an unprecedented opportunity to examine the novel tau/microglial senescence hypothesis and reveal aging-related responses of human microglia to pathological tau.

For the following reasons, our study provides first *in vivo* evidence demonstrating that human microglia respond to pathological tau with accelerated senescence. First, scRNA-seq data clearly show that DS microglia in DSAD tau group exhibit enhanced senescence compared to DS microglia in Cont tau group. Senescence scoring (Fig. 6F,6G) and KEGG enrichment analysis of DEGs (Fig. S7A) consistently indicate enhanced microglial senescence in DSAD tau group. Some senescence-associated genes, such as B2M, XAF1, and ZFP36L1, are significantly upregulated in multiple clusters in DSAD tau group, particularly the intermediate, transition cluster 0 (Fig. 6N). Many DEGs that are downregulated in DSAD tau group are involved in mitochondrial functions and cell metabolism, indicating decreased metabolic processes in DS microglia in DSAD tau group. Decreased metabolic processes also suggest cellular senescence (Lopez-Otin et al., 2013; Vasileiou et al., 2019). Notably, a recent study using an optimized cell purification method to isolate microglia from frozen human AD brain tissue for RNA-seq analysis also shows that human AD microglia exhibit accelerated aging (Srinivasan et al., 2020). Moreover, our scRNA-seq and qPCR analyses show that DS microglia in DSAD tau group express much lower levels of proinflammatory cytokines and chemokines than DS microglia in the Cont tau group, suggesting that pathological tau may not induce massive activation of DS microglia. Interestingly, as opposed to human DS microglia, the host mouse microglia in the same chimeric mouse brains in DSAD tau group exhibit hypertrophic morphology, suggestive of activation.

Second, histological evidence also points to enhanced senescence in DSAD tau group. As compared to DS microglia in Cont tau group, DS microglia in DSAD tau group show increased expression of ferritin, a robust biomarker of cellular senescence in microglia (Lopes et al., 2008; Simmons et al., 2007; Verina et al., 2011), and display dystrophic morphology with shortened and fragmented processes, characteristics of senescent microglia. Lastly, our mechanistic studies also reveal enhanced microglial senescence in DSAD tau group. Our scRNA-seq analysis of DS microglia chimeric mouse brains show that type I IFN signaling is significantly upregulated in DS microglia in DSAD tau group, as compared to Cont tau group. Previous studies in human AD brain tissues report increased production of type I IFNs (DiPatre and Gelman, 1997; Overmyer et al., 1999; Sheng et al., 1997; Taylor et al., 2014; Taylor et al., 2018). In our chimeric brains, we also find that injection of pathological tau can induce upregulation of type I IFNs. The receptors for type I IFNs are expressed in microglia in the brain tissue from AD patients (Yamada and Yamanaka, 1995). Consistently, our flow cytometry analysis demonstrates the upregulated expression of IFNARs in xeno-transplanted DS microglia, although their expression levels were not further altered by pathological tau. Importantly, elevated type I IFN signaling has been shown to promote cellular senescence and aging (Frisch and MacFawn, 2020; Yu et al., 2015). By inhibiting *IFNARs* in xenografted microglia, we demonstrate that inhibition of *IFNARs* in DS microglia prevents their senescence after DSAD tau injection. Moreover, pathological tau can also induce senescence of control microglia in the chimeric mouse brain (Fig. S8E). We propose that the elevated type 1 IFN signaling largely mediates the senescent phenotypes triggered by p-tau in human microglia, and overexpression of IFNARs due to trisomy 21 further accelerates the occurrence of the p-tau-induced senescence in DS microglia. A previous study (Bussian et al., 2018) has demonstrated a causal link between the accumulation of tau-pathology-induced senescent cells and cognition-associated neuronal loss. Clearance of senescent glial cells, including senescent microglia, prevents tau-dependent pathology and cognitive decline, suggesting that senescent glial cells play a critical role in both the initiation and progression of tau-mediated neurodegeneration. Our findings suggest that new therapeutic strategies targeting *IFNARs* could prevent human microglial senescence to potentially slow the progression of AD in DS.

Accumulation of Aβ in individuals with DS starts as early as childhood and lasts lifelong (Lott and Head, 2005, 2019). Previous studies have characterized the responses of human microglia to pathological Aβ in chimeric mouse brain models and consistently reported the transition to activated microglia state (Hasselmann et al., 2019a; Mancuso et al., 2019). Soluble Aβ proteins can induce microglia to prune synapses in early AD, which may underly the synaptic deficits occur in early AD and mild cognitive impairment before onset of plaques (Hong et al., 2016). Interestingly, a recent study further reported that Aβ pathology induces activation of type 1 IFN signaling in microglia, which drives microglia activation and microglia-mediated synaptic loss at early stages of β-amyloidosis (Roy et al., 2022). In our study, we find that DS microglia exhibit enhanced synaptic pruning functions in the developing chimeric mouse brains, due to their overexpression of IFNARs and elevated type -1 IFN signaling. Thus, the dysfunctional DS microglia in the developing brain likely contribute to the progression to AD and dementia later in life in people with DS. In addition, as compared to the recent studies (Hong et al., 2016; Roy et al., 2022), our microglia chimeric mice challenged by pathological tau is more of modeling AD at later stages when p-tau proteins are produced or brain regions where p-tau is preferentially accumulated over Aβ, such as the hippocampus (Braak and Del Tredici, 2015; Sanchez-Mejias et al., 2016). Recent RNA-seq analysis of human AD microglia reports aging profiles of AD microglia, which is not a direct response to amyloid pathology (Srinivasan et al., 2020). Here we find that pathological tau can directly induce aging of human microglia in vivo in chimeric mouse brains. Nevertheless, we support the notion that parenchymal deposition of Aβ and microglia activation is likely necessary for triggering and driving tau pathology (Edwards, 2019; Hopp et al., 2018; Ising et al., 2019; Long and Holtzman, 2019; Pascoal et al., 2021). In parallel with Aβ-pathology-induced microglial activation, microglia show rapid proliferative response (Condello et al., 2015). This early and sustained microglial proliferation may indirectly promote senescence (Hu et al., 2021). Recent RNA-seq analysis of human AD microglia also suggests that aging profiles of AD microglia are not a direct response to amyloid pathology (Srinivasan et al., 2020). Therefore, pathological Aβ and tau may collude to promote microglial senescence. Importantly, in future studies, our human microglial chimeric mouse brain model will provide new opportunities to investigate how pathological tau-induced human microglial senescence further contributes to tau spread, neurodegeneration, and dementia.

## Limitations of the Study

There are limitations of using this chimeric mouse model for disease modeling. First, there is a lack of peripheral adaptive immune system in the chimeric mice, due to the use of immunodeficient mice. We recently discussed potential approaches to developing chimeric mice with various types of functional neural cells as well as brain innate immune system and peripheral adaptive immune system derived from the same human donor (Jiang et al., 2020). Second, it is important to use behavioral tests to evaluate cognitive functions of the chimeric mice and how the manipulation of the type 1 IFN signaling may affect their behavioral phenotypes. We performed behavioral tests, including open field, novel object recognition, and elevated plus maze tests. However, there was no significant difference among the tested groups (Fig. S9A-C). We use *Rag2/IL2r*γ-deficient mice that also express human CSF1, which is crucial for the survival of xenografted human microglia (Hasselmann et al., 2019b). As previously reported, Rag2 is necessary for proper retinal development and *Rag2*^-/-^ mice are blind (Alvarez-Lindo et al., 2019; Han et al., 2013). Thus, these *Rag2*^-/-^ immunodeficient mice used in our study are likely not ideal for behavioral tests that require visual input. This issue could be potentially circumvented by using other immunodeficient mouse strains expressing human IL-34 or CSF-1 that can support the survival of engrafted human microglia (Mathews et al., 2019; Svoboda et al., 2019).

Moreover, DS is associated with a significantly increased risk of autism spectrum disorder (ASD) (de Graaf et al., 2015; Wiseman et al., 2015). Accumulating evidence has indicated that defects in the synaptic pruning mediated by microglia are closely linked to the development of ASD (Lukens and Eyo, 2022). Performing behavioral tests concerning ASD-related social interactions may help reveal differences in cognitive functions between control and DS microglial chimeras.

## Acknowledgements

This work was in part supported by grants from the NIH (R01NS102382, R01NS122108, and R01AG073779 to P.J.). We thank the UCI-ADRC, which is funded by NIH/NIA Grant P30AG066519 and the Brightfocus Foundation (BFF17-0008), for providing us with DSAD and control human brain tissues. PHF-1 antibody was a gift from Dr. Peter Davies at Albert Einstein College of Medicine, NY, USA. We thank Dr. Xiaobing Zhang from Florida State University and Dr. Zhengquan Tang from Anhui University in China for their suggestions on PPF and LTP recordings. We are also thankful to Mr. Haipeng Xue from Florida International University for the assistance with FISH experiment and Ms. Maharaib Syed from Rutgers University for the assistance with immunohistochemistry.

## Author Contributions

M.J. and P.J. designed experiments and interpreted data; M.J. carried out most of experiments with technical assistance from R.X. and M.M.A; Z.M. and R.P.H. performed the gene expression analyses and helped with RNA-seq data interpretation; S.Z. and P.X. helped with flow cytometry experiments; L.W., M.B., and Z.P.P. performed electrophysiological recordings; A.C.M. and E.H. prepared human brain tissue extracts; A.J. and K.W. helped with scRNA-seq sample preparation; Y.L. helped with characterization of hiPSC lines and hiPSC-derived PMPs; E.H. provided critical suggestions to the overall research direction. P.J. directed the project and wrote the manuscript together with M.J. and input from all co-authors.

## Competing Financial Interests

The authors declare no competing financial interests.

## Methods

### hiPSC lines generation, culture, and quality control

A total of six hiPSC lines were used in this study: three healthy control (Cont) and DS iPSC cell lines (Table S1). The DS hiPSC lines include two DS hiPSC lines (DS1, female; and DS2, male) and isogenic Di-DS3 and Tri-DS3 hiPSCs that were generated from a single female patient. The hiPSC lines were fully characterized and completely de-identified (Chen et al., 2014; Xu et al., 2019). All hiPSCs were cultured on dishes coated with hESC-qualified Matrigel (Corning) in mTeSR plus media (STEMCELL Technologies) under a feeder-free condition. The hiPSCs were passaged with ReLeSR media (STEMCELL Technologies) once per week.

### Differentiation and culture of PMPs and pNPCs

PMPs were generated from the three pairs of Cont and DS hiPSC cell lines using a previously established protocol (Haenseler et al., 2017). The yolk sac embryoid bodies (YS-EBs) were generated by treating the YS-EBs with mTeSR 1 media (STEMCELL Technologies) supplemented with bone morphogenetic protein 4 (BMP4, 50 ng/ml), vascular endothelial growth factor (VEGF, 50 ng/ml), and stem cell factor (SCF, 20 ng/ml) for 6 days. To stimulate myeloid differentiation, the YS-EBs were plated on dishes with X-VIVO 15 medium (Lonza) supplemented with interleukin-3 (IL-3, 25 ng/ml) and macrophage colony-stimulating factor (M-CSF, 100 ng/ml). At 4-6 weeks after plating, human PMPs emerged into the supernatant and were continuously produced for more than 3 months. Human pNPCs were generated from Cont2 hiPSCs (Chen et al., 2016; Xu et al., 2019). The pNPCs were cultured in a medium, which is composed of a 1:1 mixture of Neurobasal (Thermo Fisher Scientific) and DMEM/F12 (Hyclone), supplemented with 1x N2, 1x B27-RA (Thermo Fisher Scientific), FGF2 (20 ng/ml, Peprotech), CHIR99021 (3 mM, Biogems), human leukemia inhibitory factor (hLIF, 10 ng/ml, Millipore), SB431542 (2 mM), and ROCK inhibitor Y-27632 (10 mM, Tocris). The pNPCs were passaged with TrypLE Express (Thermo Fisher Scientific) once per week. pNPCs within 6 passages were used for organoid generation.

### Brain organoid culture

Each brain organoid was generated from a total of 10,000 cells (7,000 pNPCs and 3,000 PMPs), in each well of ultra-low-attachment 96-well plates in the presence of ROCK inhibitor Y-27632 (10 mM) (Xu et al., 2021). The culture medium was composed of a 1:1 mixture of PMP medium and NPC medium (1:1 mixture of Neurobasal and DMEM/F12, supplemented with 1x N2, 1x B27-RA, and FGF2 (20 ng/ml, Peprotech) for three days (day 3). Then, the organoids were transferred to ultra-low-attachment 6-well plates and cultured with 1:1 mixture of PMP medium and NPC medium for another 11 days (2 weeks). The medium was replenished every two days, and the cell culture plates were kept on an orbital shaker at a speed of 80 rpm/min starting from day 8. To promote neural and microglial differentiation, organoids were cultured in differentiation media, comprised of a 1:1 mixture of Neurobasal and DMEM/F12, supplemented with 1x N2 (Thermo Fisher Scientific), BDNF (20 ng/ml, Peprotech), GDNF (20 ng/ml, Peprotech), dibutyryl-cyclic AMP (1mM, Sigma), ascorbic acid (200 nM, Sigma), IL-34 (100 ng/ml, Peprotech), and Granulocyte-macrophage colony-stimulating factor (GM-CSF, 10 ng/ml, Peprotech) from day 15 onwards. As shown in our recent study (Xu et al., 2021), cells are highly proliferative in the early stages of organoid development. Ventricular zone-like structures containing proliferative neural stem cells are seen in organoids even after long-term culture, whereas PMPs differentiate into microglia that are not actively proliferating. Thus, the different proliferation rates between pNPCs vs. PMPs as well as protracted proliferation of human neural stem cells likely lead to a lower ratio of microglia cells in organoids than the starting 7:3 ratio. The medium was replenished every other day. After 4 weeks, the organoids were used for further experimentation.

### In vitro differentiation of PMPs to microglia

PMPs were differentiated in the medium composed of DMEM/ F12 supplemented with N2, 2 mM Glutamax, 100 U/mL penicillin and 100 mg/mL streptomycin, 100 ng/mL M-CSF (Peprotech), 100 ng/mL IL-34 (Peprotech), and 10 ng/mL GM-CSF (Peprotech) for two weeks (Haenseler et al., 2017). The medium was changed once a week. After two weeks, cells were collected for Western blotting.

### Animals and cell transplantation

All animal work was performed without gender bias with the approval of the Rutgers University Institutional Animal Care and Use Committee. PMPs were collected from the supernatant and suspended at a concentration of 100,000 cells/µl in PBS. Cells were then injected into the brains of P0 Rag2^−/−^hCSF1 immunodeficient mice (C;129S4-*Rag2^tm1.1Flv^ Csf1tm1^(CSF1)Flv^ Il2rg^tm1.1Flv^/J*, The Jackson Laboratory). The transplantation sites were bilateral from the midline = ±1.0 mm, posterior from bregma = −2.0 mm, and dorsoventral depths = −1.5 and −1.2 mm (Xu et al., 2020). All pups were placed in ice for 4-5 mins to anesthetize. The pups were then injected with 0.5 μl of cells into each site (four sites total), using a digital stereotaxic device (David KOPF Instruments) that was equipped with a neonatal mouse adapter (Stoelting). The pups were weaned at three weeks and kept for further experimentation at different time points.

### Preparation of soluble S1 fractions

Soluble S1 fractions from human samples (provided by the University of California Alzheimer’s Disease Research Center (UCI-ADRC) and the Institute for Memory Impairments and Neurological Disorders) were prepared as described before (Sanchez-Mejias et al., 2016). The human tissues were homogenized in TBS (20 mM Tris-HCl, 140 mM NaCl, pH 7.5) containing protease and phosphatase inhibitors (Roche). Homogenates were ultracentrifuged (4 °C for 60 min) at 100,000×*g* (Optima MAX Preparative Ultracentrifuge, Beckman Coulter). Supernatants, S1 fractions, were aliquoted and stored at −80 °C.

### Tau quantification by ELISA

The total amount of soluble tau in S1 fractions was determined by using an ELISA kit (human Tau, Invitrogen), according to the manufacturer’s protocol. The ELISA experiments were repeated in three independent experiments using triplicate replicas.

### Intracerebral adult brain injection

All adult brain injections were performed using a Kopf stereotaxic apparatus (David Kopf, Tujunga, CA). Stereotaxic surgery was performed on two months old transplanted Rag2^−/−^hCSF1 immunodeficient mice. The mice were aseptically injected with human brain extracts (Cont tau or DSAD tau) in the dorsal hippocampus and the overlying cortex (bregma: −2.5 mm; lateral: +2 mm; depth: −2.4 mm and −1.8 mm from the skull). A dose of tau at 3.2 µg, which was previously shown to be able to induce tau pathology in tau transgenic mice (Boluda et al., 2015; Iba et al., 2013; Peeraer et al., 2015), was injected into each human microglial chimeric mouse. Concentrations of tau per injection site were 0.8 μg/µl each site for both DSAD tau and Cont tau. The mice were injected with 1 μl of Cont or DSAD tau into each site (four sites total). After the injection, the mice were kept for further experiments.

### Flow cytometric analysis

Single-cell suspensions from the chimeric brains were washed and suspended in PBS with 1% BSA and 1 mg/ml of anti-FcR to block FcR binding. After 10 min of incubation on ice, the appropriate primary Abs, unconjugated or conjugated to different fluorescent markers, were added to the cells at a concentration of 1-10 μg/ml and incubated for 30 mins on ice. Then, the cells were washed twice in PBS+1% BSA and fixed in PBS+1% paraformaldehyde. Flow cytometric analysis was performed on a FACS Calibur or Cytek Aurora cytometer, and results were analyzed with FlowJo software (TreeStar).

### RNA isolation and quantitative reverse transcription PCR

Total RNA was extracted using TRIzol reagent (Thermo Fisher Scientific, 15596026), and 600 µg RNA was reverse transcribed into complementary DNA (cDNA) using TaqMan™ Reverse Transcription Reagents (Thermo Fisher Scientific; N8080234). Total DNA was prepared with Superscript III First- Strand kit (Invitrogen). Real-time PCR was performed on the ABI 7500 Real-Time PCR System using the TaqMan Fast Advanced Master Mix (Thermo Fisher Scientific). All primers are listed in Table S12. The 2^−ΔΔ^*^Ct^* method was used to calculate relative gene expression after normalization to the *GAPDH* internal control.

### Bulk RNA-seq

PMPs generated from the three pairs of Cont and DS hiPSC lines were used for RNA extraction and RNA sequencing sample preparation. Total RNA was prepared with an RNAeasy kit (QIAGEN) (Chen et al., 2014) and libraries were constructed by using 600 ng of total RNA from each sample and utilizing a TruSeqV2 kit from Illumina (Illumina, San Diego, CA) following the manufacturer’s suggested protocol. The libraries were subjected to 75 bp paired read sequencing using a NextSeq500 Illumina sequencer to generate approximately 30 to 35 million paired-reads per sample. Fastq files were generated using the Bcl2Fastq software, version 1.8.4. The genome sequence was then indexed using the *rsem- prepare-reference* command. Each fastq file was trimmed using the fastp program (v0.12.2) and then aligned to the human genome (GRCh38/hg38) using HISAT2 (v.2.2.0). Gene counts were extracted from bam files using Rsubread/featureCounts in R with a UCSC transcript map. To analyze the transcripts, CPM > 1 was set as a cutoff to filter transcripts. Fold change > 2 was set as criteria to filter differential expressed genes (DEGs). The raw gene counts were processed with R package edgeR for differential expression analysis, exact Test hypothesis testing was used for each pairwise analysis.

Differentially expressed genes were defined with |log_2_FC| > 1 between control and DS groups, and p- value < 0.05. Gene ontology enrichment analysis was performed with DAVID version 6.8, upregulated and downregulated genes were enriched separately. Data shown in the paper were from terms enriched in *GOTERM_BP_DIRECT*.

### Tissue dissociation for scRNA-seq

After perfusion with cold PBS, whole brains were dissected and stored briefly in 1x DPBS. The tissue dissociation was then performed utilizing the Adult Brain Dissociation Kit (Miltenyi) according to the manufacturer’s instructions. Following tissue dissociation, the tissue was dissected into 1xmm^3^ pieces and placed into the C-tubes equipped with enzymes. These samples were further dissociated in gentleMACS OctoDissociator with heaters (Miltenyi) using the established preprogrammed protocol. Following enzymatic digestion, samples were isolated using a 70 μm cell strainer and pelleted by centrifugation. Myelin and debris byproducts were removed by debris removal solution, overlaid with 6mL of 1X DPBS, and spun at 3000xg for 10 min at 4°C. The supernatant was discarded, and the cell pellet was processed for Magnetic Isolation.

### Magnetic Isolation of human microglia for scRNA-seq

Dissociated cell pellets were resuspended in 160 µL FACS buffer (0.5% BSA in 1X DPBS) + 40uL Mouse cell removal beads (Miltenyi) and incubated at 4°C for 15 min. The resulting samples were then isolated using LS columns and the MidiMACs separator (Miltenyi), the human cells were collected in the flow through. Using centrifugation (10 min, 400xg), the cells were pelleted and resuspended to 1,000 cells per microliter in FACS buffer for scRNA-seq library preparation, according to the previous study (Hasselmann et al., 2019a).

### ScRNA-seq

Following magnetic isolation, we used Chromium™ Single Cell 3′ Library and Gel Bead Kit v3.1, Chromium™ Single Cell Chip G Kit, and Chromium™ i7 Multiplex Kit, 96 rxns for capture and library preparation. The libraries were analyzed on Agilent 4200 TapeStation System using High Sensitivity D1000 ScreenTape Assay and quantified using KAPA qPCR. Libraries were then normalized to 10 nM before being pooled together. Next, the pooled library was clustered and sequenced on Illumina NovaSeq 6000 S4 flowcell for 150bp paired-end sequencing, Read 1 and Read 2 are sequenced from both ends of the fragment. For each individual library, the sequencing data from four unique indexes were combined before further analysis.

### Raw data pre-processing

The sequencing data was aligned with the pooled mouse (mm10, Ensembl 93) and human (hg19, Ensembl 87) reference genomes (10x Genomics pre-built human and mouse reference genome v3.0.0) and interpreted via barcodes analyzed with Cell Ranger software (10x Genomics, v.6.0.2). The resulting matrices of gene count × barcodes were coded by individual sample identifiers and loaded into Seurat (v.4.0.3) software in R/Bioconductor. An initial analysis revealed a distinct cluster of human- expressing cells. Cells with gene number <200 or >10,000, and mitochondria gene >10% were excluded. Leaving a total number of 47,826 cells from four samples. Cells were summarized for all genes by species and those with >75% of reads aligning with hg19 were selected as human. Raw hg19 counts from these cells were loaded into a new Seurat object, normalized and clustered.

### Data processing and quality control

The separated fastq files were re-aligned to the human hg19 reference genome using the same version of Cell Ranger as described above. Seurat R package v4.0.4 was used for quality control and dimensional reduction analysis. Gene-barcode matrices were merged after inputting into R as Seurat objects. Cells with gene number <200 or >4,000, and mitochondria gene >10% were excluded. Leaving a total number of 7,790 cells.

### Dimensional reduction and differential expression analysis

Data were normalized by RPM following log transformation and top 2,000 highly variable genes were selected for scaling and principal component analysis (PCA). The top 15 principal components were used for downstream uniform manifold approximation and projection (UMAP) visualization and clustering. Louvain algorithm with resolution 0.2 was used to cluster cells, which resulted in 6 distinct cell clusters. Cluster 5 was removed due to the size of the cluster (15 cells). Differential expression analysis was performed by using Wilcoxon rank sum test embedded in *FindMarkers* function from Seurat. A gene was considered to be differentially expressed if it was detected in at least 25% of one group and with at least 0.25 log fold change between two groups and the significant level of Benjamini– Hochberg (BH) adjusted p-value < 0.05.

### Gene ontology (GO) and gene set enrichment analysis (GSEA)

GO analysis used the g:Profiler website (https://biit.cs.ut.ee/gprofiler/gost). All genes were calculated between Control and DSAD group before logFC, and adjusted p-value filtration were ranked by logFC value. Pre-defined gene sets were used to test their distribution on the ranked gene list, GSEA analysis and visualization were performed by *gseaplot* function from clusterProfiler v4.0.5 R package.

### Pseudotime trajectory inference analysis

Single cell pseudotime trajectory was predicted by Monocle 3 v1.0.0. The gene-cell matrix was exported from Seurat object as input into Monocle3 to generate cds object, along with the metadata information from Seurat. Cells were re-clustered by Monocle3 using the default parameters. To learn trajectory graph and order cells in pseudotime, clustering information from Seurat object was projected into the cds object. Cells in the cluster with enriched homeostatic genes expression scores were defined as root cells.

### shRNA knockdown

*IFNAR1* shRNA (sc-35637-V), *IFNAR2* shRNA (sc-40091-V), and non-targeting control shRNA (sc- 108080) lentiviral particles were purchased from Santa Cruz Biotechnology. The *IFNAR1* and *IFNAR2* shRNA lentivirus particles carried three different shRNAs specifically targeted *IFNAR1/IFNAR2* gene expression. DS2 and Tri-DS3 hiPSCs infected with lentivirus carrying the control shRNA (DS2 or Tri- DS3+ Cont^shRNA^) or IFNAR1/2 shRNA (DS2 or Tri-DS3 + IFNAR1/2^shRNA^) were used in this study. Lentivirus was mixed with polybrene (5 mg/ml) in mTeSR plus and applied to undifferentiated hiPSCs overnight. The following day, the transduction medium was removed and replaced with fresh media.

After three days of culture, a puromycin (0.75 mg/ml) selection was used for two weeks to select for transducing hiPSCs. Stable hiPSCs were then used for PMP differentiation. Knockdown efficiency was confirmed by examining IFNAR1 and IFNAR2 expression at both mRNA and protein levels.

### Electrophysiology

#### Whole cell patch-clamp recording

Brain slice mEPSCs recordings was carried out following an established protocol (Liu et al., 2017). Mice were anesthetized with Euthasol, and brains were quickly removed into ice cold (0 °C) oxygenated ACSF cutting solution (in mM): 50 sucrose, 2.5 KCl, 0.625 CaCl_2_, 1.2 MgCl_2_, 1.25 NaH_2_PO_4_, 25 NaHCO_3_, and 2.5 glucose, pH to 7.3 with NaOH. 300 µm sagittal section slices were cut using a vibratome (VT 1200S; Leica). After 1 hour of recovery (30 °C) in artificial cerebrospinal fluid (ACSF) (in mM)125 NaCl, 2.5 KCl, 2.5 CaCl_2_, 1.2 MgCl_2_, 1.25 NaH_2_PO_4_, 56NaHCO_3_, and 10 glucose. For miniature excitatory postsynaptic currents (mEPSC) recording, a Cs2^+^-based solution was used, which consisted of (in mM): 40 CsCl, 3.5 KCl, 10 HEPES, 0.05 EGTA, 90 K-gluconate, 1.8 NaCl, 1.7 MgCl_2_, 2 ATP-magnesium, 0.4 GTP-sodium, and 10 phosphocreatine. mEPSCs were recorded at a holding potential of −70 mV in the presence of 50 μM PTX and 1 μM tetrodotoxin (TTX; VWR, Catalog#89160– 628). Electrophysiological data were analyzed using Clampfit 10.5 (Molecular Devices, USA). Brain slices after recordings were used for post hoc immunostaining with hTMEM119 to verify colocalization of xeno-grafted human microglia with Neurobiotin fluorescence-positive neurons labeled via recording electrodes.

### Extracellular hippocampal slice recording

The extracellular recording of hippocampal slices was performed as described previously (Castillo et al., 2002). Mice brains were quickly immersed in cold (4°C) oxygenated cutting solution containing (in mM): 206 Sucrose, 11 D-Glucose, 2.5 KCl, 1 NaH_2_PO_4_, 10 MgCl_2_, 2 CaCl_2,_ and 26 NaHCO_3_. Transverse 400 μm bilateral hippocampal sections were cut and transferred to an oxygenated ACSF contained (in mM): 125 NaCl, 2.5 KCl, 2.5 CaCl_2_, 1.2 MgCl_2_, 1.25 NaH2PO_4_, 26 NaHCO_3_, and 2.5 glucose. After at least half hour 34°C of recovery, slices were transferred at room temperature and incubated for 1 hour. Field EPSPs were recorded with glass pipette (2-3MΩ). The stimulation electrode was placed in dorsal CA1b stratum radiatum equidistant from the CA1 stratum pyramidale. Single-pulse baseline stimulation was applied at 0.05 Hz with baseline intensity set to 40–50% of the maximum population spike-free fEPSP amplitude. LTP was induced by 4X100Hz stimulation with 200 ms interval between bursts.

### Western Blotting

Western blotting was performed as described previously (Jin et al., 2018). Lysates from cells, S1 fractions, and human brain tissues were prepared using a RIPA buffer and the protein contents were measured using a Pierce^TM^ BCA Protein Assay Kit (Thermo Scientific). Proteins were separated on 12% SDS-PAGE gradient gel and transferred onto a nitrocellulose membrane. The membrane was blocked with 5% non-fat milk for one hour at room temperature. After blocking, the membrane was incubated with primary antibody overnight at 4 °C overnight. Secondary antibodies conjugated to HRP were used in addition to ECL reagents (Pierce ECL Plus Western Blotting Substrate, Thermo scientific) for immunodetection. All the S1 fractions were derived from DSAD and Cont human brain tissues (Table S4). For quantification of band intensity, blots from three independent experiments for the molecule of interest were used. Signals were measured by ImageJ software and represented as relative intensity versus control. GAPDH was used as an internal control to normalize band intensity.

### Tissue immunostaining, image acquisition, and analysis

Mouse brains and organoids were fixed with 4% paraformaldehyde. The organoids were placed in 25% sucrose and mouse brains were placed in 20% and later in 30% sucrose for dehydration. Following dehydration, organoids and brain tissues were immersed in OCT and frozen for sectioning. The frozen tissues were cryo-sectioned with 30-μm thickness for immunofluorescence staining. The organoids and tissues were blocked with a blocking solution (5% goat or donkey serum in PBS with 0.2% or 0.8% Triton X-100) at room temperature (RT) for 1 hour. The primary antibodies were diluted in the same blocking solution and incubated with the organoids or tissues at 4 °C overnight (all the primary antibodies are listed in Supplementary Table 12). The sections were washed with PBS and incubated with secondary antibodies for 1 hour at RT. After washing with PBS, the slides were mounted with anti- fade Fluoromount-G medium containing 1, 40,6-diamidino-2-phenylindole dihydrochloride (DAPI) (Southern Biotechnology).

All images were captured with a Zeiss 800 confocal microscope. Large scale images in Fig. 3A, 3G and S4B were obtained by confocal tile scan by the Zen software (Zeiss). To obtain a 3D reconstruction, images were processed by the Zen software (Zeiss). To visualize synaptic pruning and phagocytic function, super-resolution images in Figs. 2-7, Supplementary Figs. 2-3, and 6 were acquired by Zeiss Airyscan super-resolution microscope at 63X with 0.2mm z-steps. To generate 3D- surface rendered images, super-resolution images were processed by Imaris software (Bitplane 9.5). To visualize the engulfment of PSD95^+^, Synapsin I^+^, AT8^+^ and PHF-1^+^ p-tau within microglia, CD68^+^ phagolysosome inside microglia, fluorescence found outside of the microglia were subtracted from the image via the mask function in Imaris software. For hTMEM119^+^ human microglia and Iba-1^+^/ hN- negative mouse microglia morphology analysis, the Filament function was used to generate filaments for each cell in the images, and dendrites were automatically rendered based on the hTMEM119 and Iba1 signal. The number of positive cells from each section was counted after a Z projection. The dystrophic (Iba-1^+^hN^+^) microglial morphological changes after tau injection were assessed by calculating the soma size and process length using the software Fiji with MorpholibJ plugin and integrated library as described before (Flores-Aguilar et al., 2020; Legland et al., 2016). Soma size was measured with a closing morphological filter connecting dark pixels and process length was analyzed with Skeletonize function.

### Behavioral tests

We examined three groups of mice: (1) control chimeric mice that received transplantation of Cont 2 and Cont 3 cells; (2) DS + Cont^shRNA^ chimeric mice that received transplantation of DS2+Cont^shRNA^ or Tri-DS3+Cont^shRNA^ cells; (3) DS + IFNAR1/2^shRNA^ chimeric mice that received transplantation of DS2+ IFNAR1/2^shRNA^ or Tri-DS3+ IFNAR1/2^shRNA^ cells; All behavior tests were performed in a randomized order by two investigator double-blinded to treatment.

### Open field test

Mice were placed into a clear chamber (40 long x 40 wide x 40 cm high) under dim ambient light conditions. The activity was monitored by overhead video camera and analyzed by Anymaze software (Stoelting Co., IL) for 5 min in a single trial. Four squares were determined as the center and the other twelve squares along the center as the periphery. The collected results included total distance traveled and the total number of entries into the center of the field.

### Elevated plus maze

The elevated plus maze was performed in a grey, non-reflective base plate cross (arms 35 cm long x 5 cm wide) elevated 50 cm above the floor. Two opposite arms were enclosed by the grey walls (35 cm long x 15 cm high), and the two other arms were open. A mouse was placed in the center of the apparatus facing an enclosed arm and allowed to explore the apparatus for 5 min. The total traveling distance and the time spent on the open arms were recorded.

### Novel object recognition

As previously described (Xu et al., 2019), two sample objects in one clear box were used to examine learning and memory with 24 hours delays. Before testing, mice were habituated in a clear chamber (40 long x 40 wide x 40 cm high) for 10 minutes on 2 consecutive days under ambient light conditions. The activities were monitored by an overhead video camera and analyzed by an Anymaze software (Stoelting Co., IL). First, two identical objects, termed as ‘familiar’, were placed into the chamber, and the mouse was placed at the mid-point of the wall opposite the objects. After 10 minutes to explore the objects, the mouse was returned to the home cage. After 24 hours, one of the ‘familiar’ objects used for the memory acquisition was replaced with a ‘new’ object similar to the ‘familiar’ one. The mouse was again placed in the chamber for 3 minutes to explore the objects; The total traveling distance and the time spent investigating the new objects was assessed. The preference to the novel object was calculated as Time exploring new object / (Time exploring new object + Time exploring familiar object).

### Golgi staining

The mice were anesthetized, and whole brains were collected for Golgi staining using the FD Rapid Golgi Stain Kit (FD Neuro Technologies, Columbia, MD, USA; PK401), according to the manufacturer’s introduction. Z-stack images were acquired with a widefield microscope (Olympus BX63), and spine density was quantified using ImageJ software (NIH).

### Statistical analysis

All data are represented as mean ± SEM. When only two independent groups were compared, significance was determined by using two-tailed unpaired t test with Welch’s correction. When three or more groups were compared, one-way ANOVA with Bonferroni post-hoc test was used. A p-value of < 0.05 was considered significant. All the analyses were done in GraphPad Prism v.9. All experiments were independently performed at least three times with similar results.

## Data availability

The GEO accession number of bulk RNA sequencing and the single-cell RNA sequencing data reported in this study is GSE189227.

---

**Fig S1.**
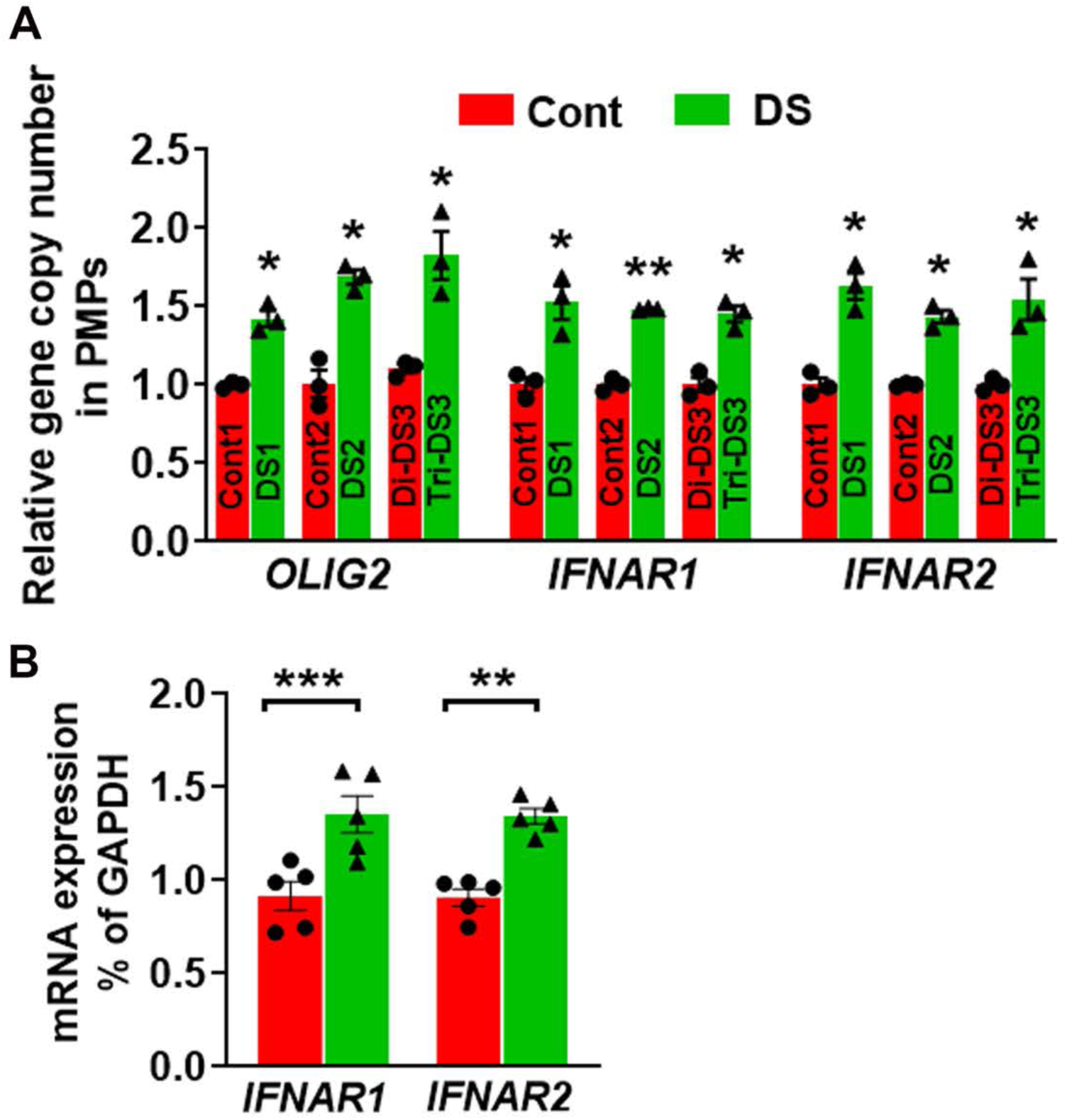
Related to **Fig 1**. Gene expression of hiPSC-derived PMPs. (A) qPCR results of genomic DNA showing the relative *OLIG2, IFNAR1,* and *IFNAR2* DNA copy numbers in Cont (Cont1, Cont2, and Di-DS3) and DS (DS1, DS2, and Tri-DS3) PMPs (n=3, each experiment was repeated three times). Student’s t test, \*\*\**P* < 0.001. Data are presented as mean ± SEM. (B) qPCR analysis of *IFNAR1* and *IFNAR2* mRNA expression in three pairs of Cont and DS hiPSC derived PMPs. (n=5, the data were pooled from the three pairs of Cont and DS hiPSC-derived PMPs, Student’s t test, \*\*\**P* <0.001. Data are presented as mean ± SEM.

**Fig S2.**
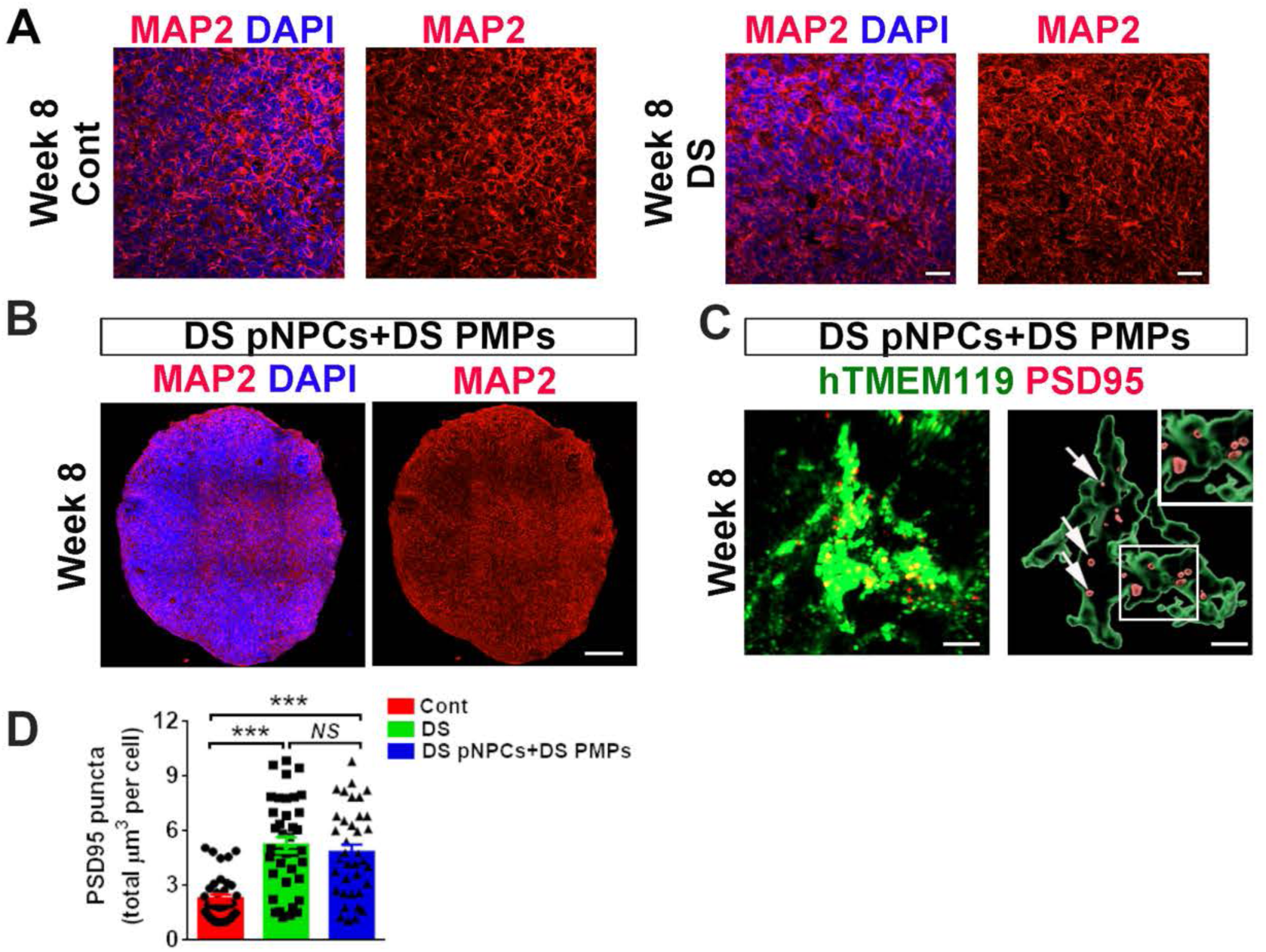
Related to **Fig 2**. Abnormal development of DS microglia in cerebral organoids. (A) Representative images of MAP^+^ neurons in Cont and DS microglia-containing cerebral organoids, Scale bar: 20 μm. (B) Representative images of MAP^+^ neurons in DS pNPCs with DS microglia-containing cerebral organoids at week 8. Scale bar: 20 μm. (C) Quantification of pooled data from Cont, DS, and DS pNPCs with DS PMPs groups at week 8 (n=3 for each experiment, 4 to 6 organoids from each line are used). One-way ANOVA test, \*\*\**P* < 0.001, *NS*, not significant. Data are presented as mean ± SEM.

**Fig S3.**
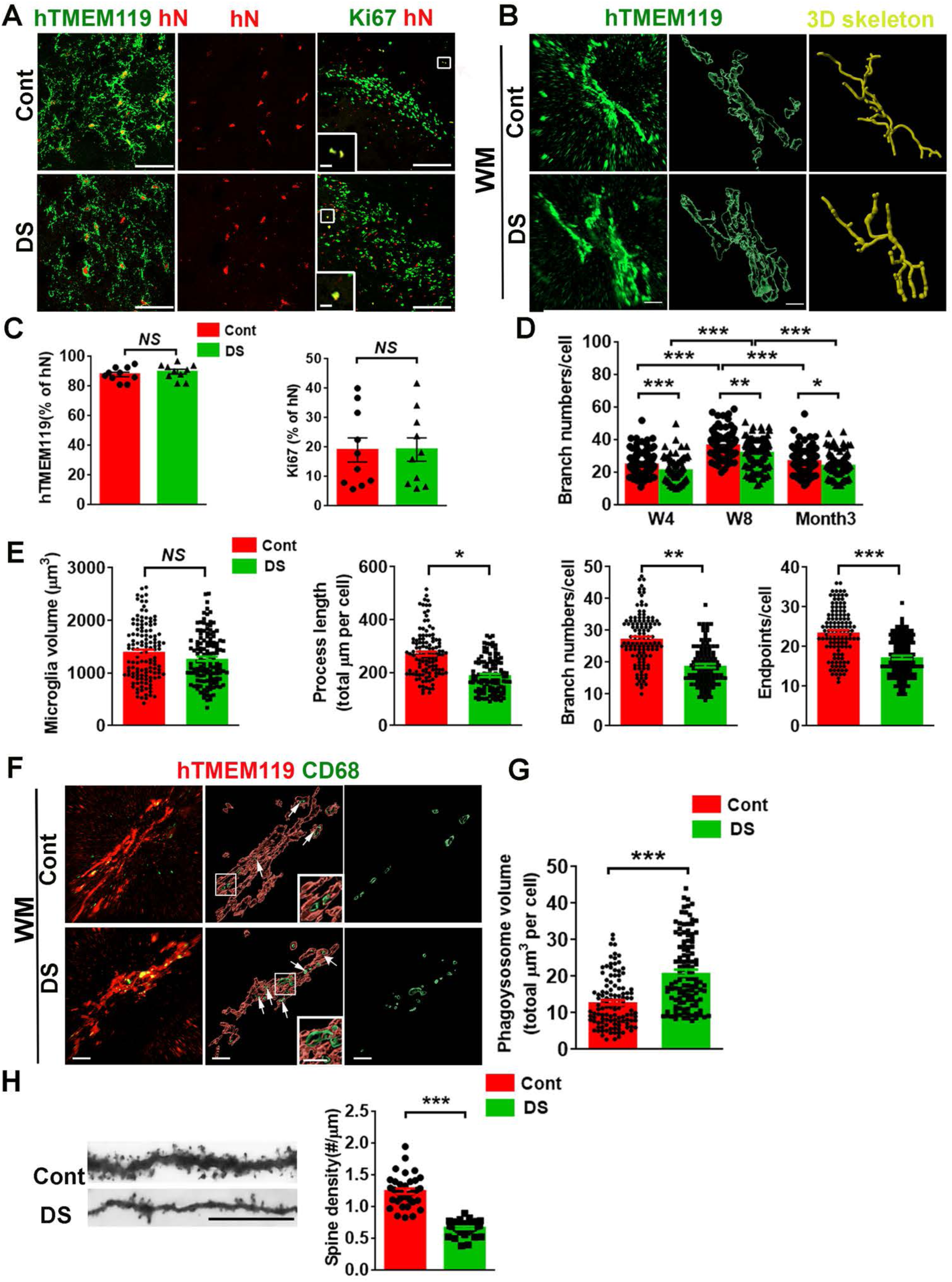
Related to **Fig 3**. Abnormal development of DS microglia in chimeric mice. (A) Representative images of hTMEM119^+^hN^+^ and Ki67^+^hN^+^ in Cont and DS chimeric mice. Scale bar: 50 μm and 10 μm in the original and enlarged images, respectively. (B) Representative raw fluorescent super-resolution, 3D surface rendered, and 3D skeletonization images of hTMEM119 staining of week 8 chimeras. Scale bar: 7 μm. (C) Quantification of the percentage of hTMEM119 and Ki67 in hN^+^ cells from Cont and DS chimeric mice (n=10 mice per group), Student’s t test, \**P* < 0.05 and \*\*\**P* < 0.001. Data are presented as mean ± SEM. (D) Quantification of branch numbers (n=120-160 from 3-5 mice per group). Student’s t test, \**P* < 0.05, \*\*\**P* < 0.001. Data are presented as mean ± SEM. (E) Quantification of microglial volumes, process length, branch numbers and endpoints (n=116-136 from 3-4 mice per group). Student’s t test, \**P* < 0.05, \*\**P* < 0.01 and \*\*\**P* < 0.001, *NS*, not significant. Data are presented as mean ± SEM. (F) Representative super-resolution and 3D surface rendered images showing hTMEM119 and CD68 staining of week 8 chimeras. Arrows indicate CD68^+^ phagolysosome. Scale bar: 5 μm and 3 μm in the original and enlarged images, respectively. (G) Quantification of CD68+phagolysosome volume in microglia (n=115-133 from 3-4 mice per group), Student’s t test, \*\*\**P* < 0.001. Data are presented as mean ± SEM. (H) Golgi staining and quantification of dendritic spines in 2-3 months old chimeric hippocampus (n=34 dendrites per group). Scale bar, 10 μm. Student’s t test, \*\*\**P* < 0.001. Data are presented as mean ± SEM.

**Fig S4.**
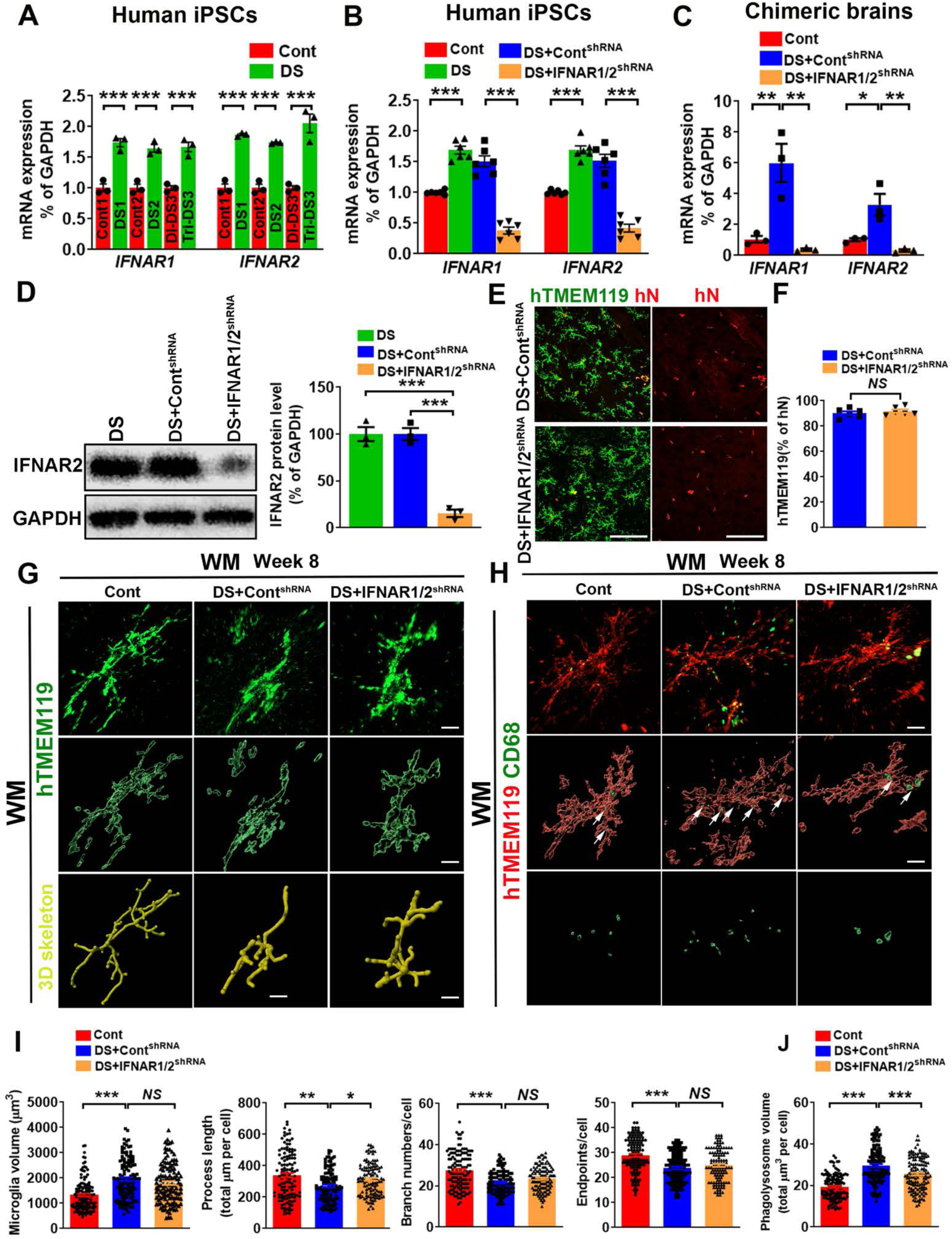
Related to **Fig 4**. Characterization of *IFNARs* knockdown in chimeras. (A) qPCR analysis of *IFNAR1* and *IFNAR2* mRNA expression in three pairs of Cont and DS hiPSC lines. (n=3, each experiment was repeated three times), Student’s t test, \*\*\**P* <0.001. Data are presented as mean ± SEM. (B) qPCR analysis of *IFNAR1* and *IFNAR2* mRNA expression in two DS hiPSC lines expressing IFNAR1/2^shRNA^ or Cont^shRNA^, (n=5, each experiment was repeated five times), Student’s t test, \*\*\**P* <0.001. Data are presented as mean ± SEM. (C) qPCR analysis of *IFNAR1*and *IFNAR2* mRNA expression in 3-4-month-old chimeric mice, Cont, DS + Cont^shRNA^ and DS + IFNAR1/2^shRNA^ chimeras (n=3 mice per group). Student’s t test, \**P* < 0.05 and \*\**P* < 0.01. Data are presented as mean ± SEM. (D) Western blotting analysis of IFNAR2 expression in DS, DS + Cont^shRNA^ and DS + IFNAR1/2^shRNA^ microglia cells (n=3, each experiment was repeated three times). One-way ANOVA test. \*\*\**P* <0.001. Data are presented as mean ± SEM. (E) Representative images of hTMEM119 and hN in DS + Cont^shRNA^ and DS + IFNAR1/2^shRNA^ in chimeric mice. Scale bar: 50 μm. (F) Quantification of the percentage of hTMEM119 in hN^+^ cells in DS + Cont^shRNA^ and DS + IFNAR1/2^shRNA^ chimeric mice (n=7 mice per group). Student’s t test, *NS*, not significant. Data are presented as mean ± SEM. (G) Representative raw fluorescent super-resolution, 3D surface rendered, and 3D skeletonization images of hTMEM119 staining in Cont, and DS + Cont^shRNA^ and DS + IFNAR1/2^shRNA^ chimeric mice. Scale bar: 5 μm. (H) Representative super-resolution and 3D surface rendered images showing hTMEM119 and CD68 staining in Cont, and DS + Cont^shRNA^ and DS + IFNAR1/2^shRNA^ chimeric mice. Arrows indicate PSD95^+^ puncta in the CD68^+^ phagolysosome. Scale bar: 5 μm. (I) Quantification of microglial volume, process length, branch numbers and endpoints (n=118-136 from 3-4 mice per group), One-way ANOVA test. \**P* < 0.05, \*\**P* < 0.01 and \*\*\**P* < 0.001. Data are presented as mean ± SEM. (J) Quantification of CD68^+^ phagolysosome volume (n=128-136 from 3-4 mice per group). One-way ANOVA test, \**P* < 0.05 and \*\*\**P* < 0.001. Data are presented as mean ± SEM.

**Fig S5.**
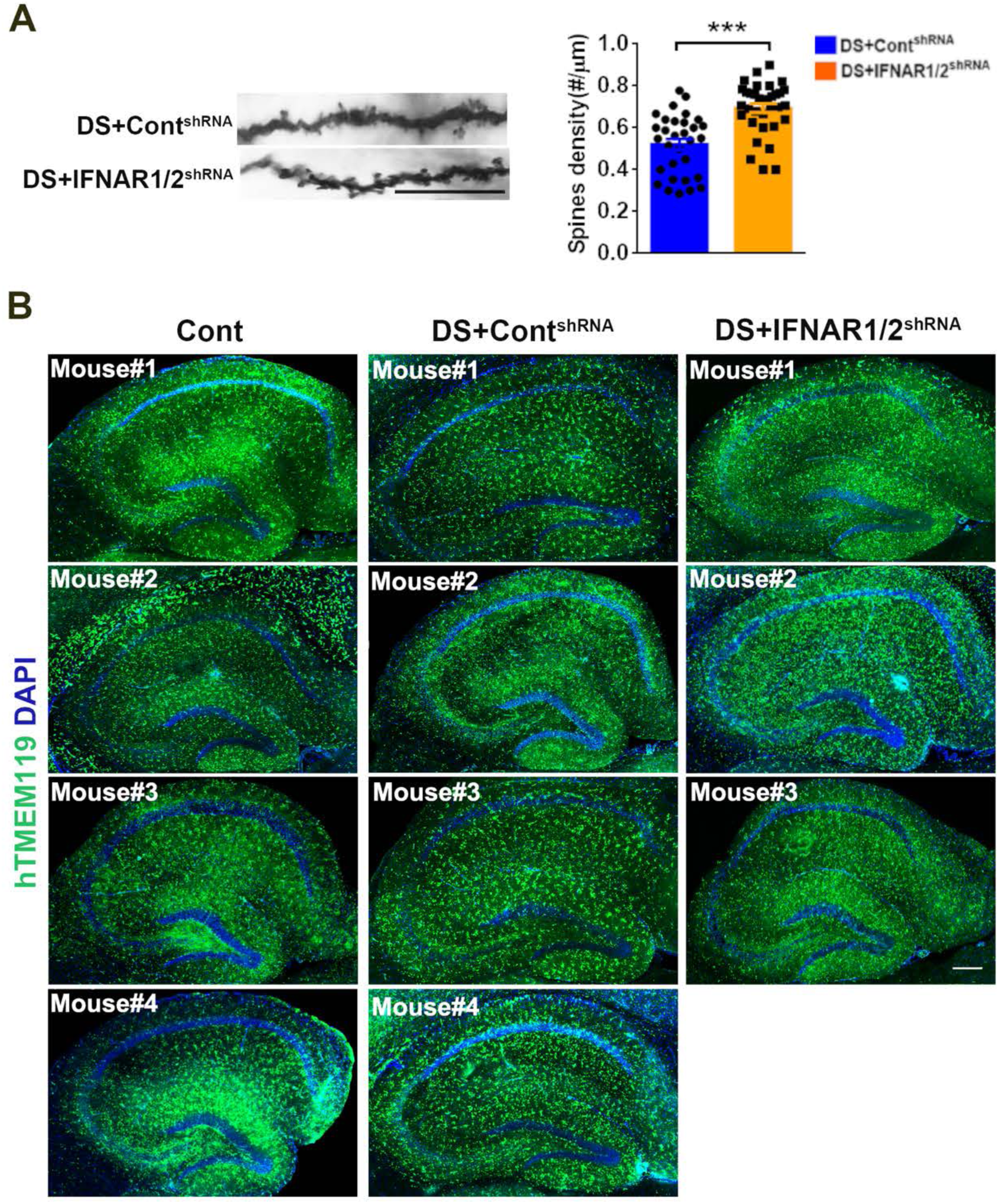
Related to **Fig 4**. Characterization of *IFNARs* knockdown in chimeras. (A) Golgi staining and quantification of dendritic spines in 2-3 months old DS + Cont^shRNA^ and DS + IFNAR1/2^shRNA^ chimeric hippocampus (n=31-34 dendrites per group). Scale bar, 10 μm. Student’s t test, \*\*\**P* < 0.001. Data are presented as mean ± SEM. (B) Representative images of LTP recording slices from transverse sagittal brain sections showing the distribution of transplanted Cont, DS + Cont^shRNA^ and DS + IFNAR1/2^shRNA^ hiPSC-derived microglia at month 3-4 old chimeric mice. Scale bar: 200 μm.

**Fig S6.**
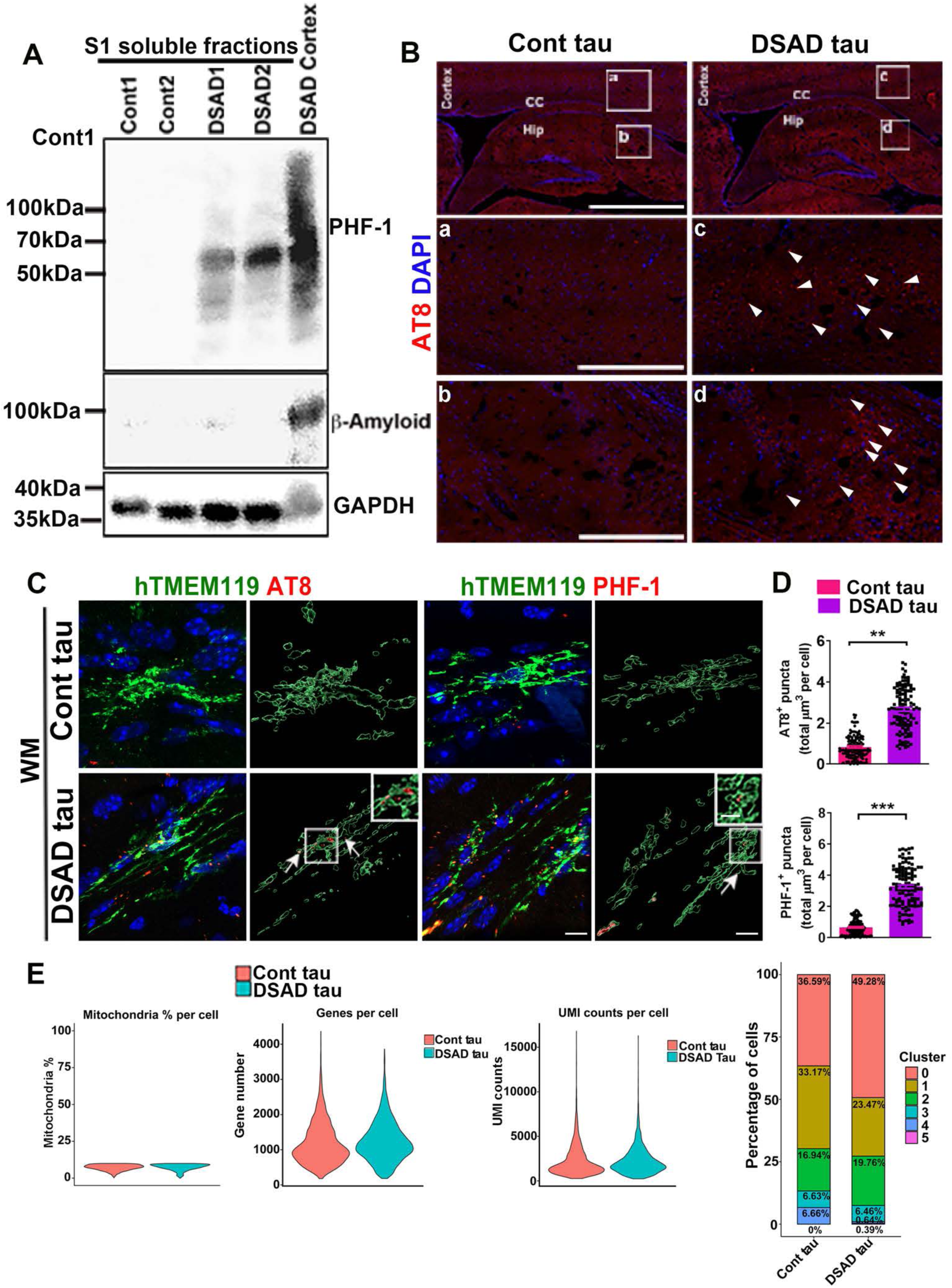
Related to **Fig 5**. Quality control of human brain tissue-derived soluble tau and scRNA-seq analysis. (A) Western blotting analysis showing PHF-1 and Aβ expression in the S1 soluble fraction from Cont and DSAD brain tissues. (B) Representative images of sagittal brain sections showing the distribution of AT8 at two weeks post-injection. Arrowheads indicate AT8^+^ p-tau. Scale bar: 500 μm and 50 μm in the original and enlarged images, respectively. (C) Representative images of hTMEM119, AT8, and PHF-1 staining in 4-5-month-old chimeric mice. Arrows indicate AT8^+^ or PHF-1^+^ p-tau. Scale bars: 5 μm and 3 μm in the original and enlarged images, respectively. (D) Quantification of AT8^+^ and PHF-1^+^ in DS microglia (n=120 from 4 mice per group). Student’s t test, \*\**P* < 0.01 and \*\*\**P* < 0.001. Data are presented as mean ± SEM. (E) Violin plots representing the mitochondrial gene ratio, the number of genes per cell, and the count of unique molecular identifiers (UMIs) in Cont and DSAD tau groups. (F) The percentage of each cluster in Cont and DSAD tau groups.

**Fig S7.**
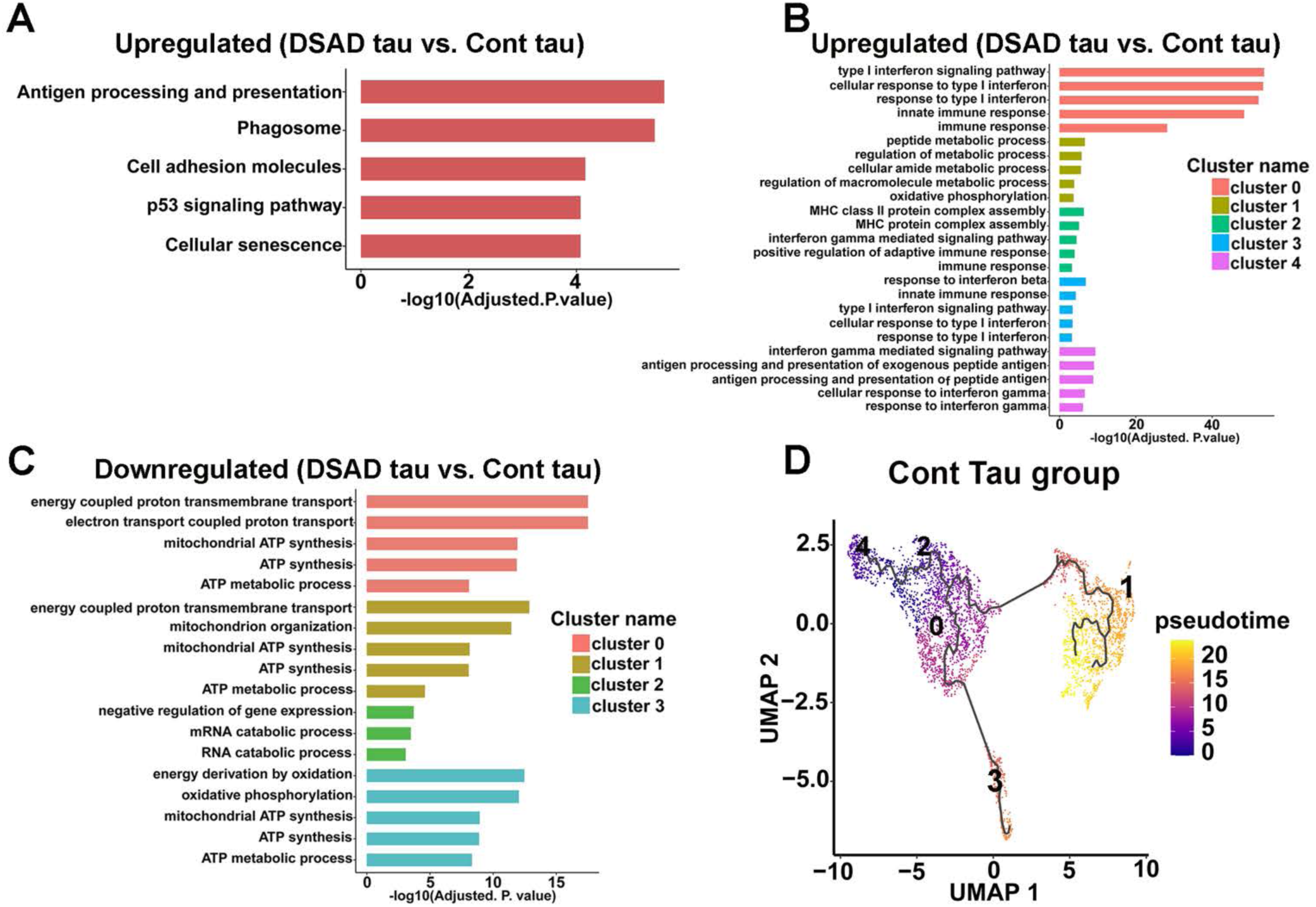
Related to **Fig 6**. scRNA-seq analysis of DS microglia exposed to pathological tau. (A) KEGG enrichment analyses of the upregulated DEGs in DS microglia. (B, C) GO analyses of upregulated and downregulated DEGs from each cluster. (D) UMAP representation of the trajectory of DS microglia in response to Cont tau, cells are colored by pseudotime.

**Fig S8.**
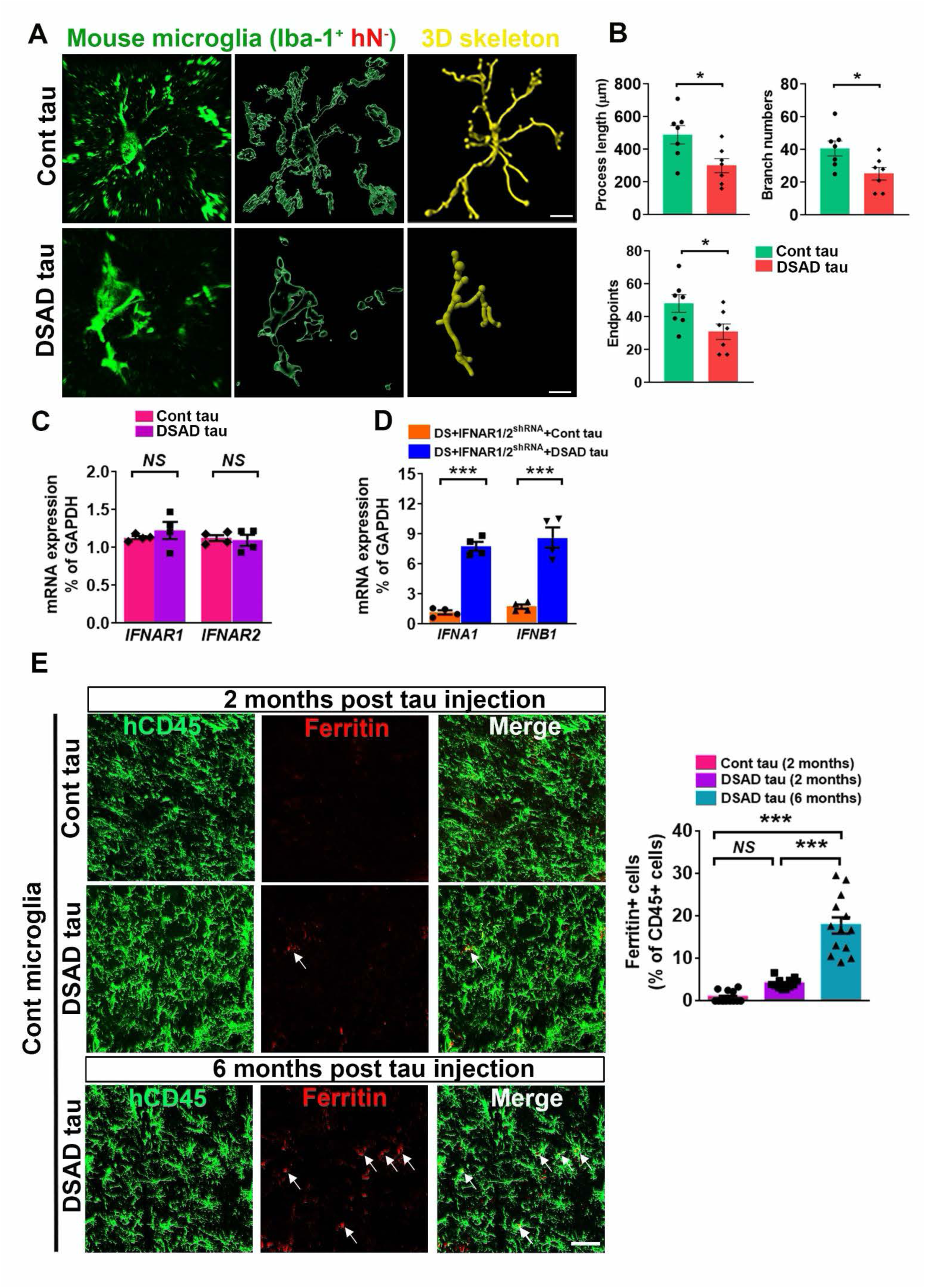
Related to **Fig 7**. Pathological tau induces hypertrophy of mouse microglia. (A) Representative raw fluorescent super-resolution, 3D surface rendered, and 3D skeletonization images of mouse microglia (Iba^+^hN^-^) in Cont tau and DSAD tau chimeric mice. Scale bar: 5 μm. (B) Quantification of the process length, branch numbers, and endpoints in mouse microglia (Iba^+^hN^-^) (n=7 mice per group), Student’s t test, \**P* < 0.05, \*\**P* < 0.01. Data are presented as mean ± SEM. (C) qPCR analysis of *IFNAR1* and *IFNAR2* mRNA expression (n=4 mice per group). Student’s t test, *NS*, not significant. Data are presented as mean ± SEM. (D) qPCR analysis of *IFNA1* and *IFNB1* mRNA expression (n=4 mice per group). Student’s t test, \*\*\**P* < 0.001. Data are presented as mean ± SEM. (E) Representative images showing colocalization of hCD45^+^ and Ferritin^+^ staining in Cont chimeras at two and six months after tau injection. Arrows indicate Ferritin^+^ and/or hCD45^+^ staining. Scale bar: 50 μm. Quantification of the percentage of Ferritin in hCD45^+^ cells (n =13 fields from 2 mice per group). Student’s t test, \*\*\**P* < 0.001 and *NS*, not significant. Data are presented as mean ± SEM.

**Fig S9.**
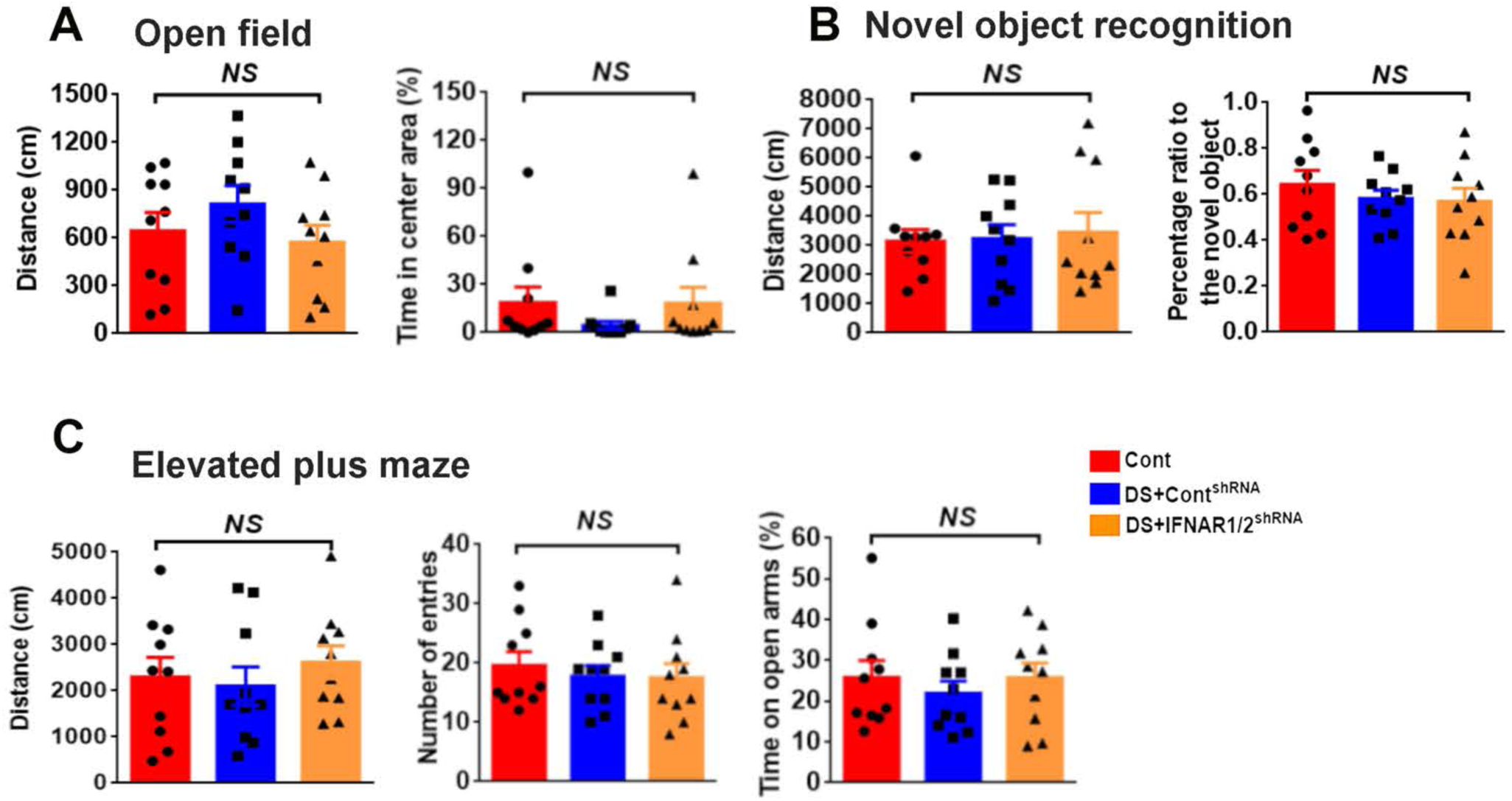
Related to Discussion. Behavior tests in chimeric mice. (A) Quantification of traveling distance and percentage of time in the center of open-field test in 3-4 months old Cont, DS + Cont^shRNA^, and DS + IFNAR1/2^shRNA^ chimeric mice (n=10 mice for each group). One-way ANOVA test. *NS*, no significance. (B) Quantification of traveling distance and preference ratio to the novel object in the three groups of 3- 4 months old chimeric mice (n=10 mice for each group). One-way ANOVA test. NS, no significance. (C) Quantification of the traveling distance, the number of entries, and the time mice spend on open arms (n=10 mice for each group) in the Elevated plus maze test from 3-4 months old chimeric mice. One-way ANOVA test. *NS*, no significance.

## Notes

### Competing Interest Statement

The authors have declared no competing interest.

### Summary of Updates

Added new experiments, including electrophysiological recordings and behavioral testings

## References

Abud, E.M., Ramirez, R.N., Martinez, E.S., Healy, L.M., Nguyen, C.H.H., Newman, S.A., Yeromin, A.V., Scarfone, V.M., Marsh, S.E., Fimbres, C., et al. (2017). iPSC-Derived Human Microglia-like Cells to Study Neurological Diseases. Neuron 94, 278–293 e279.

Alvarez-Lindo, N., Baleriola, J., de Los Rios, V., Suarez, T., and de la Rosa, E.J. (2019). RAG-2 deficiency results in fewer phosphorylated histone H2AX foci, but increased retinal ganglion cell death and altered axonal growth. Sci Rep 9, 18486.

Araya, P., Waugh, K.A., Sullivan, K.D., Nunez, N.G., Roselli, E., Smith, K.P., Granrath, R.E., Rachubinski, A.L., Enriquez Estrada, B., Butcher, E.T., et al. (2019). Trisomy 21 dysregulates T cell lineages toward an autoimmunity-prone state associated with interferon hyperactivity. Proc Natl Acad Sci U S A 116, 24231–24241.

Arvanitakis, Z., Grodstein, F., Bienias, J.L., Schneider, J.A., Wilson, R.S., Kelly, J.F., Evans, D.A., and Bennett, D.A. (2008). Relation of NSAIDs to incident AD, change in cognitive function, and AD pathology. Neurology 70, 2219–2225.

Avelar, R.A., Ortega, J.G., Tacutu, R., Tyler, E.J., Bennett, D., Binetti, P., Budovsky, A., Chatsirisupachai, K., Johnson, E., Murray, A., et al. (2020). A multidimensional systems biology analysis of cellular senescence in aging and disease. Genome Biol 21, 91.

Bar, E., and Barak, B. (2019). Microglia roles in synaptic plasticity and myelination in homeostatic conditions and neurodevelopmental disorders. Glia 67, 2125–2141.

Baruch, K., Deczkowska, A., David, E., Castellano, J.M., Miller, O., Kertser, A., Berkutzki, T., Barnett-Itzhaki, Z., Bezalel, D., Wyss-Coray, T., et al. (2014). Aging. Aging-induced type I interferon response at the choroid plexus negatively affects brain function. Science 346, 89–93.

Belichenko, P.V., Kleschevnikov, A.M., Becker, A., Wagner, G.E., Lysenko, L.V., Yu, Y.E., and Mobley, W.C. (2015). Down Syndrome Cognitive Phenotypes Modeled in Mice Trisomic for All HSA 21 Homologues. PloS one 10, e0134861.

Benito-Kwiecinski, S., and Lancaster, M.A. (2020). Brain Organoids: Human Neurodevelopment in a Dish. Cold Spring Harb Perspect Biol 12.

Bennett, M.L., Bennett, F.C., Liddelow, S.A., Ajami, B., Zamanian, J.L., Fernhoff, N.B., Mulinyawe, S.B., Bohlen, C.J., Adil, A., Tucker, A., et al. (2016). New tools for studying microglia in the mouse and human CNS. Proc Natl Acad Sci U S A 113, E1738–1746.

Blank, T., and Prinz, M. (2017). Type I interferon pathway in CNS homeostasis and neurological disorders. Glia 65, 1397–1406.

Boluda, S., Iba, M., Zhang, B., Raible, K.M., Lee, V.M., and Trojanowski, J.Q. (2015). Differential induction and spread of tau pathology in young PS19 tau transgenic mice following intracerebral injections of pathological tau from Alzheimer’s disease or corticobasal degeneration brains. Acta Neuropathol 129, 221–237.

Borowski, A.B., Boesteanu, A.C., Mueller, Y.M., Carafides, C., Topham, D.J., Altman, J.D., Jennings, S.R., and Katsikis, P.D. (2007). Memory CD8+ T cells require CD28 costimulation. J Immunol 179, 6494–6503.

Braak, H., and Del Tredici, K. (2015). The preclinical phase of the pathological process underlying sporadic Alzheimer’s disease. Brain 138, 2814–2833.

Brownjohn, P.W., Smith, J., Solanki, R., Lohmann, E., Houlden, H., Hardy, J., Dietmann, S., and Livesey, F.J. (2018). Functional Studies of Missense TREM2 Mutations in Human Stem Cell-Derived Microglia. Stem cell reports 10, 1294–1307.

Bussian, T.J., Aziz, A., Meyer, C.F., Swenson, B.L., van Deursen, J.M., and Baker, D.J. (2018). Clearance of senescent glial cells prevents tau-dependent pathology and cognitive decline. Nature 562, 578–582.

Butovsky, O., and Weiner, H.L. (2018). Microglial signatures and their role in health and disease. Nat Rev Neurosci 19, 622–635.

Carson, M.J., Reilly, C.R., Sutcliffe, J.G., and Lo, D. (1998). Mature microglia resemble immature antigen-presenting cells. Glia 22, 72–85.

Casano, A.M., and Peri, F. (2015). Microglia: multitasking specialists of the brain. Dev Cell 32, 469–477.

Castillo, P.E., Schoch, S., Schmitz, F., Sudhof, T.C., and Malenka, R.C. (2002). RIM1alpha is required for presynaptic long-term potentiation. Nature 415, 327–330.

Chen, C., Jiang, P., Xue, H., Peterson, S.E., Tran, H.T., McCann, A.E., Parast, M.M., Li, S., Pleasure, D.E., Laurent, L.C., et al. (2014). Role of astroglia in Down’s syndrome revealed by patient-derived human-induced pluripotent stem cells. Nat Commun 5, 4430.

Chen, Y., and Colonna, M. (2021). Microglia in Alzheimer’s disease at single-cell level. Are there common patterns in humans and mice? J Exp Med 218.

Choong, X.Y., Tosh, J.L., Pulford, L.J., and Fisher, E.M. (2015). Dissecting Alzheimer disease in Down syndrome using mouse models. Front Behav Neurosci 9, 268.

Clavaguera, F., Akatsu, H., Fraser, G., Crowther, R.A., Frank, S., Hench, J., Probst, A., Winkler, D.T., Reichwald, J., Staufenbiel, M., et al. (2013). Brain homogenates from human tauopathies induce tau inclusions in mouse brain. Proc Natl Acad Sci U S A 110, 9535–9540.

Clavaguera, F., Bolmont, T., Crowther, R.A., Abramowski, D., Frank, S., Probst, A., Fraser, G., Stalder, A.K., Beibel, M., Staufenbiel, M., et al. (2009). Transmission and spreading of tauopathy in transgenic mouse brain. Nat Cell Biol 11, 909–913.

Condello, C., Yuan, P., Schain, A., and Grutzendler, J. (2015). Microglia constitute a barrier that prevents neurotoxic protofibrillar Abeta42 hotspots around plaques. Nat Commun 6, 6176.

Cuadrado, E., and Barrena, M.J. (1996). Immune dysfunction in Down’s syndrome: primary immune deficiency or early senescence of the immune system? Clin Immunol Immunopathol 78, 209–214.

Das, I., and Reeves, R.H. (2011). The use of mouse models to understand and improve cognitive deficits in Down syndrome. Dis Model Mech 4, 596–606.

de Graaf, G., Buckley, F., and Skotko, B.G. (2015). Estimates of the live births, natural losses, and elective terminations with Down syndrome in the United States. Am J Med Genet A 167A, 756–767.

Deczkowska, A., Keren-Shaul, H., Weiner, A., Colonna, M., Schwartz, M., and Amit, I. (2018). Disease-Associated Microglia: A Universal Immune Sensor of Neurodegeneration. Cell 173, 1073–1081.

DiPatre, P.L., and Gelman, B.B. (1997). Microglial cell activation in aging and Alzheimer disease: partial linkage with neurofibrillary tangle burden in the hippocampus. J Neuropathol Exp Neurol 56, 143–149.

Doran, E., Keator, D., Head, E., Phelan, M.J., Kim, R., Totoiu, M., Barrio, J.R., Small, G.W., Potkin, S.G., and Lott, I.T. (2017). Down Syndrome, Partial Trisomy 21, and Absence of Alzheimer’s Disease: The Role of APP. J Alzheimers Dis 56, 459–470.

Douvaras, P., Sun, B., Wang, M., Kruglikov, I., Lallos, G., Zimmer, M., Terrenoire, C., Zhang, B., Gandy, S., Schadt, E., et al. (2017). Directed Differentiation of Human Pluripotent Stem Cells to Microglia. Stem cell reports 8, 1516–1524.

Edwards, F.A. (2019). A Unifying Hypothesis for Alzheimer’s Disease: From Plaques to Neurodegeneration. Trends Neurosci 42, 310–322.

Ejlerskov, P., Hultberg, J.G., Wang, J., Carlsson, R., Ambjorn, M., Kuss, M., Liu, Y., Porcu, G., Kolkova, K., Friis Rundsten, C., et al. (2015). Lack of Neuronal IFN-beta-IFNAR Causes Lewy Body- and Parkinson’s Disease-like Dementia. Cell 163, 324–339.

Flores-Aguilar, L., Iulita, M.F., Kovecses, O., Torres, M.D., Levi, S.M., Zhang, Y., Askenazi, M., Wisniewski, T., Busciglio, J., and Cuello, A.C. (2020). Evolution of neuroinflammation across the lifespan of individuals with Down syndrome. Brain 143, 3653–3671.

Friedman, B.A., Srinivasan, K., Ayalon, G., Meilandt, W.J., Lin, H., Huntley, M.A., Cao, Y., Lee, S.H., Haddick, P.C.G., Ngu, H., et al. (2018). Diverse Brain Myeloid Expression Profiles Reveal Distinct Microglial Activation States and Aspects of Alzheimer’s Disease Not Evident in Mouse Models. Cell Rep 22, 832–847.

Frisch, S.M., and MacFawn, I.P. (2020). Type I interferons and related pathways in cell senescence. Aging Cell 19, e13234.

Galatro, T.F., Holtman, I.R., Lerario, A.M., Vainchtein, I.D., Brouwer, N., Sola, P.R., Veras, M.M., Pereira, T.F., Leite, R.E.P., Moller, T., et al. (2017). Transcriptomic analysis of purified human cortical microglia reveals age-associated changes. Nat Neurosci 20, 1162–1171.

Galloway, A., Saveliev, A., Lukasiak, S., Hodson, D.J., Bolland, D., Balmanno, K., Ahlfors, H., Monzon- Casanova, E., Mannurita, S.C., Bell, L.S., et al. (2016). RNA-binding proteins ZFP36L1 and ZFP36L2 promote cell quiescence. Science 352, 453–459.

Geirsdottir, L., David, E., Keren-Shaul, H., Weiner, A., Bohlen, S.C., Neuber, J., Balic, A., Giladi, A., Sheban, F., Dutertre, C.A., et al. (2019). Cross-Species Single-Cell Analysis Reveals Divergence of the Primate Microglia Program. Cell 179, 1609–1622 e1616.

Gosselin, D., Skola, D., Coufal, N.G., Holtman, I.R., Schlachetzki, J.C.M., Sajti, E., Jaeger, B.N., O’Connor, C., Fitzpatrick, C., Pasillas, M.P., et al. (2017). An environment-dependent transcriptional network specifies human microglia identity. Science 356.

Green, R.C., Schneider, L.S., Amato, D.A., Beelen, A.P., Wilcock, G., Swabb, E.A., Zavitz, K.H., and Tarenflurbil Phase 3 Study, G. (2009). Effect of tarenflurbil on cognitive decline and activities of daily living in patients with mild Alzheimer disease: a randomized controlled trial. JAMA 302, 2557–2564.

Group, A.R., Lyketsos, C.G., Breitner, J.C., Green, R.C., Martin, B.K., Meinert, C., Piantadosi, S., and Sabbagh, M. (2007). Naproxen and celecoxib do not prevent AD in early results from a randomized controlled trial. Neurology 68, 1800–1808.

Group, A.R., Martin, B.K., Szekely, C., Brandt, J., Piantadosi, S., Breitner, J.C., Craft, S., Evans, D., Green, R., and Mullan, M. (2008). Cognitive function over time in the Alzheimer’s Disease Anti- inflammatory Prevention Trial (ADAPT): results of a randomized, controlled trial of naproxen and celecoxib. Arch Neurol 65, 896–905.

Guerrero, A., De Strooper, B., and Arancibia-Carcamo, I.L. (2021). Cellular senescence at the crossroads of inflammation and Alzheimer’s disease. Trends Neurosci 44, 714–727.

Haenseler, W., Sansom, S.N., Buchrieser, J., Newey, S.E., Moore, C.S., Nicholls, F.J., Chintawar, S., Schnell, C., Antel, J.P., Allen, N.D., et al. (2017). A Highly Efficient Human Pluripotent Stem Cell Microglia Model Displays a Neuronal-Co-culture-Specific Expression Profile and Inflammatory Response. Stem cell reports 8, 1727–1742.

Han, X., Chen, M., Wang, F., Windrem, M., Wang, S., Shanz, S., Xu, Q., Oberheim, N.A., Bekar, L., Betstadt, S., et al. (2013). Forebrain engraftment by human glial progenitor cells enhances synaptic plasticity and learning in adult mice. Cell Stem Cell 12, 342–353.

Hansen, D.V., Hanson, J.E., and Sheng, M. (2018). Microglia in Alzheimer’s disease. J Cell Biol 217, 459–472.

Hasselmann, J., Coburn, M.A., England, W., Figueroa Velez, D.X., Kiani Shabestari, S., Tu, C.H., McQuade, A., Kolahdouzan, M., Echeverria, K., Claes, C., et al. (2019a). Development of a Chimeric Model to Study and Manipulate Human Microglia In Vivo. Neuron 103, 1016–1033 e1010.

Hasselmann, J., Coburn, M.A., England, W., Figueroa Velez, D.X., Kiani Shabestari, S., Tu, C.H., McQuade, A., Kolahdouzan, M., Echeverria, K., Claes, C., et al. (2019b). Development of a Chimeric Model to Study and Manipulate Human Microglia In Vivo. Neuron.

Head, E., Lott, I.T., Hof, P.R., Bouras, C., Su, J.H., Kim, R., Haier, R., and Cotman, C.W. (2003). Parallel compensatory and pathological events associated with tau pathology in middle aged individuals with Down syndrome. J Neuropathol Exp Neurol 62, 917–926.

Heo, J.I., Kim, W., Choi, K.J., Bae, S., Jeong, J.H., and Kim, K.S. (2016). XIAP-associating factor 1, a transcriptional target of BRD7, contributes to endothelial cell senescence. Oncotarget 7, 5118–5130.

Heppner, F.L., Ransohoff, R.M., and Becher, B. (2015). Immune attack: the role of inflammation in Alzheimer disease. Nat Rev Neurosci 16, 358–372.

Holtman, I.R., Skola, D., and Glass, C.K. (2017). Transcriptional control of microglia phenotypes in health and disease. J Clin Invest 127, 3220–3229.

Hong, S., Beja-Glasser, V.F., Nfonoyim, B.M., Frouin, A., Li, S.M., Ramakrishnan, S., Merry, K.M., Shi, Q.Q., Rosenthal, A., Barres, B.A., et al. (2016). Complement and microglia mediate early synapse loss in Alzheimer mouse models. Science 352, 712–716.

Hopp, S.C., Lin, Y., Oakley, D., Roe, A.D., DeVos, S.L., Hanlon, D., and Hyman, B.T. (2018). The role of microglia in processing and spreading of bioactive tau seeds in Alzheimer’s disease. J Neuroinflammation 15, 269.

Horvath, S., Garagnani, P., Bacalini, M.G., Pirazzini, C., Salvioli, S., Gentilini, D., Di Blasio, A.M., Giuliani, C., Tung, S., Vinters, H.V., et al. (2015). Accelerated epigenetic aging in Down syndrome. Aging Cell 14, 491–495.

Hosseini, S., Michaelsen-Preusse, K., Grigoryan, G., Chhatbar, C., Kalinke, U., and Korte, M. (2020). Type I Interferon Receptor Signaling in Astrocytes Regulates Hippocampal Synaptic Plasticity and Cognitive Function of the Healthy CNS. Cell Rep 31, 107666.

Hu, Y., Fryatt, G.L., Ghorbani, M., Obst, J., Menassa, D.A., Martin-Estebane, M., Muntslag, T.A.O., Olmos-Alonso, A., Guerrero-Carrasco, M., Thomas, D., et al. (2021). Replicative senescence dictates the emergence of disease-associated microglia and contributes to Abeta pathology. Cell Rep 35, 109228.

Hubel, P., Urban, C., Bergant, V., Schneider, W.M., Knauer, B., Stukalov, A., Scaturro, P., Mann, A., Brunotte, L., Hoffmann, H.H., et al. (2019). A protein-interaction network of interferon-stimulated genes extends the innate immune system landscape. Nat Immunol 20, 493–502.

Hunter, M.P., Nelson, M., Kurzer, M., Wang, X., Kryscio, R.J., Head, E., Pinna, G., and O’Bryan, J.P. (2011). Intersectin 1 contributes to phenotypes in vivo: implications for Down’s syndrome. Neuroreport 22, 767–772.

Iba, M., Guo, J.L., McBride, J.D., Zhang, B., Trojanowski, J.Q., and Lee, V.M. (2013). Synthetic tau fibrils mediate transmission of neurofibrillary tangles in a transgenic mouse model of Alzheimer’s- like tauopathy. J Neurosci 33, 1024–1037.

Ioannidis, I., McNally, B., Willette, M., Peeples, M.E., Chaussabel, D., Durbin, J.E., Ramilo, O., Mejias, A., and Flano, E. (2012). Plasticity and virus specificity of the airway epithelial cell immune response during respiratory virus infection. J Virol 86, 5422–5436.

Ising, C., Venegas, C., Zhang, S., Scheiblich, H., Schmidt, S.V., Vieira-Saecker, A., Schwartz, S., Albasset, S., McManus, R.M., Tejera, D., et al. (2019). NLRP3 inflammasome activation drives tau pathology. Nature 575, 669–673.

Jiang, P., Turkalj, L., and Xu, R. (2020). High-Fidelity Modeling of Human Microglia with Pluripotent Stem Cells. Cell Stem Cell 26, 629–631.

Jimenez, S., Baglietto-Vargas, D., Caballero, C., Moreno-Gonzalez, I., Torres, M., Sanchez-Varo, R., Ruano, D., Vizuete, M., Gutierrez, A., and Vitorica, J. (2008). Inflammatory response in the hippocampus of PS1M146L/APP751SL mouse model of Alzheimer’s disease: age-dependent switch in the microglial phenotype from alternative to classic. J Neurosci 28, 11650–11661.

Jimenez, S., Navarro, V., Moyano, J., Sanchez-Mico, M., Torres, M., Davila, J.C., Vizuete, M., Gutierrez, A., and Vitorica, J. (2014). Disruption of amyloid plaques integrity affects the soluble oligomers content from Alzheimer disease brains. PloS one 9, e114041.

Jin, M.M., Wang, F., Qi, D., Liu, W.W., Gu, C., Mao, C.J., Yang, Y.P., Zhao, Z., Hu, L.F., and Liu, C.F. (2018). A Critical Role of Autophagy in Regulating Microglia Polarization in Neurodegeneration. Front Aging Neurosci 10, 378.

Keating, D.J., Chen, C., and Pritchard, M.A. (2006). Alzheimer’s disease and endocytic dysfunction: clues from the Down syndrome-related proteins, DSCR1 and ITSN1. Ageing Res Rev 5, 388–401.

Keren-Shaul, H., Spinrad, A., Weiner, A., Matcovitch-Natan, O., Dvir-Szternfeld, R., Ulland, T.K., David, E., Baruch, K., Lara-Astaiso, D., Toth, B., et al. (2017). A Unique Microglia Type Associated with Restricting Development of Alzheimer’s Disease. Cell 169, 1276–1290 e1217.

Kierdorf, K., Erny, D., Goldmann, T., Sander, V., Schulz, C., Perdiguero, E.G., Wieghofer, P., Heinrich, A., Riemke, P., Holscher, C., et al. (2013). Microglia emerge from erythromyeloid precursors via Pu.1- and Irf8-dependent pathways. Nat Neurosci 16, 273–280.

Krasemann, S., Madore, C., Cialic, R., Baufeld, C., Calcagno, N., El Fatimy, R., Beckers, L., O’Loughlin, E., Xu, Y., Fanek, Z., et al. (2017). The TREM2-APOE Pathway Drives the Transcriptional Phenotype of Dysfunctional Microglia in Neurodegenerative Diseases. Immunity 47, 566–581 e569.

Legland, D., Arganda-Carreras, I., and Andrey, P. (2016). MorphoLibJ: integrated library and plugins for mathematical morphology with ImageJ. Bioinformatics 32, 3532–3534.

Li, Q., and Barres, B.A. (2018). Microglia and macrophages in brain homeostasis and disease. Nat Rev Immunol 18, 225–242.

Ling, K.H., Hewitt, C.A., Tan, K.L., Cheah, P.S., Vidyadaran, S., Lai, M.I., Lee, H.C., Simpson, K., Hyde, L., Pritchard, M.A., et al. (2014). Functional transcriptome analysis of the postnatal brain of the Ts1Cje mouse model for Down syndrome reveals global disruption of interferon-related molecular networks. BMC Genomics 15, 624.

Loh, X.Y., Sun, Q.Y., Ding, L.W., Mayakonda, A., Venkatachalam, N., Yeo, M.S., Silva, T.C., Xiao, J.F., Doan, N.B., Said, J.W., et al. (2020). RNA-Binding Protein ZFP36L1 Suppresses Hypoxia and Cell- Cycle Signaling. Cancer Res 80, 219–233.

Long, J.M., and Holtzman, D.M. (2019). Alzheimer Disease: An Update on Pathobiology and Treatment Strategies. Cell 179, 312–339.

Lopes, K.O., Sparks, D.L., and Streit, W.J. (2008). Microglial dystrophy in the aged and Alzheimer’s disease brain is associated with ferritin immunoreactivity. Glia 56, 1048–1060.

Lopez-Otin, C., Blasco, M.A., Partridge, L., Serrano, M., and Kroemer, G. (2013). The hallmarks of aging. Cell 153, 1194–1217.

Lott, I.T., and Head, E. (2005). Alzheimer disease and Down syndrome: factors in pathogenesis. Neurobiol Aging 26, 383–389.

Lott, I.T., and Head, E. (2019). Dementia in Down syndrome: unique insights for Alzheimer disease research. Nature reviews Neurology 15, 135–147.

Lukens, J.R., and Eyo, U.B. (2022). Microglia and Neurodevelopmental Disorders. Annu Rev Neurosci.

Lunnon, K., Keohane, A., Pidsley, R., Newhouse, S., Riddoch-Contreras, J., Thubron, E.B., Devall, M., Soininen, H., Kloszewska, I., Mecocci, P., et al. (2017). Mitochondrial genes are altered in blood early in Alzheimer’s disease. Neurobiol Aging 53, 36–47.

Mancuso, R., Van Den Daele, J., Fattorelli, N., Wolfs, L., Balusu, S., Burton, O., Liston, A., Sierksma, A., Fourne, Y., Poovathingal, S., et al. (2019). Stem-cell-derived human microglia transplanted in mouse brain to study human disease. Nat Neurosci 22, 2111–2116.

Martini, A.C., Helman, A.M., McCarty, K.L., Lott, I.T., Doran, E., Schmitt, F.A., and Head, E. (2020). Distribution of microglial phenotypes as a function of age and Alzheimer’s disease neuropathology in the brains of people with Down syndrome. Alzheimers Dement (Amst) 12, e12113.

Mathews, S., Branch Woods, A., Katano, I., Makarov, E., Thomas, M.B., Gendelman, H.E., Poluektova, L.Y., Ito, M., and Gorantla, S. (2019). Human Interleukin-34 facilitates microglia-like cell differentiation and persistent HIV-1 infection in humanized mice. Mol Neurodegener 14, 12.

Meyer-Luehmann, M., and Prinz, M. (2015). Myeloid cells in Alzheimer’s disease: culprits, victims or innocent bystanders? Trends Neurosci 38, 659–668.

Morris, G.P., Clark, I.A., Zinn, R., and Vissel, B. (2013). Microglia: a new frontier for synaptic plasticity, learning and memory, and neurodegenerative disease research. Neurobiol Learn Mem 105, 40–53.

Mosser, C.A., Baptista, S., Arnoux, I., and Audinat, E. (2017). Microglia in CNS development: Shaping the brain for the future. Prog Neurobiol 149–150, 1-20.

Moyer, A.J., Gardiner, K., and Reeves, R.H. (2020). All Creatures Great and Small: New Approaches for Understanding Down Syndrome Genetics. Trends Genet.

Muffat, J., Li, Y., Yuan, B., Mitalipova, M., Omer, A., Corcoran, S., Bakiasi, G., Tsai, L.H., Aubourg, P., Ransohoff, R.M., et al. (2016). Efficient derivation of microglia-like cells from human pluripotent stem cells. Nat Med 22, 1358–1367.

Navarro, V., Sanchez-Mejias, E., Jimenez, S., Munoz-Castro, C., Sanchez-Varo, R., Davila, J.C., Vizuete, M., Gutierrez, A., and Vitorica, J. (2018). Microglia in Alzheimer’s Disease: Activated, Dysfunctional or Degenerative. Front Aging Neurosci 10, 140.

Nimmo, J., Johnston, D.A., Dodart, J.C., MacGregor-Sharp, M.T., Weller, R.O., Nicoll, J.A.R., Verma, A., and Carare, R.O. (2020). Peri-arterial pathways for clearance of alpha-Synuclein and tau from the brain: Implications for the pathogenesis of dementias and for immunotherapy. Alzheimers Dement (Amst) 12, e12070.

Ormel, P.R., Vieira de Sa, R., van Bodegraven, E.J., Karst, H., Harschnitz, O., Sneeboer, M.A.M., Johansen, L.E., van Dijk, R.E., Scheefhals, N., Berdenis van Berlekom, A., et al. (2018). Microglia innately develop within cerebral organoids. Nat Commun 9, 4167.

Otvos, L., Jr., Feiner, L., Lang, E., Szendrei, G.I., Goedert, M., and Lee, V.M. (1994). Monoclonal antibody PHF-1 recognizes tau protein phosphorylated at serine residues 396 and 404. J Neurosci Res 39, 669–673.

Overmyer, M., Helisalmi, S., Soininen, H., Laakso, M., Riekkinen, P., Sr., and Alafuzoff, I. (1999). Reactive microglia in aging and dementia: an immunohistochemical study of postmortem human brain tissue. Acta Neuropathol 97, 383–392.

Pandya, H., Shen, M.J., Ichikawa, D.M., Sedlock, A.B., Choi, Y., Johnson, K.R., Kim, G., Brown, M.A., Elkahloun, A.G., Maric, D., et al. (2017). Differentiation of human and murine induced pluripotent stem cells to microglia-like cells. Nat Neurosci 20, 753–759.

Paolicelli, R.C., Bolasco, G., Pagani, F., Maggi, L., Scianni, M., Panzanelli, P., Giustetto, M., Ferreira, T.A., Guiducci, E., Dumas, L., et al. (2011). Synaptic pruning by microglia is necessary for normal brain development. Science 333, 1456–1458.

Parker, S.E., Mai, C.T., Canfield, M.A., Rickard, R., Wang, Y., Meyer, R.E., Anderson, P., Mason, C.A., Collins, J.S., Kirby, R.S., et al. (2010). Updated National Birth Prevalence estimates for selected birth defects in the United States, 2004-2006. Birth Defects Res A Clin Mol Teratol 88, 1008–1016.

Parkhurst, C.N., Yang, G., Ninan, I., Savas, J.N., Yates, J.R., 3rd, Lafaille, J.J., Hempstead, B.L., Littman, D.R., and Gan, W.B. (2013). Microglia promote learning-dependent synapse formation through brain-derived neurotrophic factor. Cell 155, 1596–1609.

Pascoal, T.A., Benedet, A.L., Ashton, N.J., Kang, M.S., Therriault, J., Chamoun, M., Savard, M., Lussier, F.Z., Tissot, C., Karikari, T.K., et al. (2021). Microglial activation and tau propagate jointly across Braak stages. Nat Med 27, 1592–1599.

Patel, A., Yamashita, N., Ascano, M., Bodmer, D., Boehm, E., Bodkin-Clarke, C., Ryu, Y.K., and Kuruvilla, R. (2015). RCAN1 links impaired neurotrophin trafficking to aberrant development of the sympathetic nervous system in Down syndrome. Nat Commun 6, 10119.

Peeraer, E., Bottelbergs, A., Van Kolen, K., Stancu, I.C., Vasconcelos, B., Mahieu, M., Duytschaever, H., Ver Donck, L., Torremans, A., Sluydts, E., et al. (2015). Intracerebral injection of preformed synthetic tau fibrils initiates widespread tauopathy and neuronal loss in the brains of tau transgenic mice. Neurobiol Dis 73, 83–95.

Pinto, B., Morelli, G., Rastogi, M., Savardi, A., Fumagalli, A., Petretto, A., Bartolucci, M., Varea, E., Catelani, T., Contestabile, A., et al. (2020). Rescuing Over-activated Microglia Restores Cognitive Performance in Juvenile Animals of the Dp(16) Mouse Model of Down Syndrome. Neuron 108, 887–904 e812.

Platanias, L.C. (2005). Mechanisms of type-I- and type-II-interferon-mediated signalling. Nat Rev Immunol 5, 375–386.

Prasher, V.P., Farrer, M.J., Kessling, A.M., Fisher, E.M., West, R.J., Barber, P.C., and Butler, A.C. (1998). Molecular mapping of Alzheimer-type dementia in Down’s syndrome. Ann Neurol 43, 380–383.

Ransohoff, R.M. (2016). How neuroinflammation contributes to neurodegeneration. Science 353, 777–783.

Romero-Molina, C., Navarro, V., Sanchez-Varo, R., Jimenez, S., Fernandez-Valenzuela, J.J., Sanchez- Mico, M.V., Munoz-Castro, C., Gutierrez, A., Vitorica, J., and Vizuete, M. (2018). Distinct Microglial Responses in Two Transgenic Murine Models of TAU Pathology. Front Cell Neurosci 12, 421.

Roy, E.R., Chiu, G., Li, S., Propson, N.E., Kanchi, R., Wang, B., Coarfa, C., Zheng, H., and Cao, W. (2022). Concerted type I interferon signaling in microglia and neural cells promotes memory impairment associated with amyloid beta plaques. Immunity.

Rusinova, I., Forster, S., Yu, S., Kannan, A., Masse, M., Cumming, H., Chapman, R., and Hertzog, P.J. (2013). Interferome v2.0: an updated database of annotated interferon-regulated genes. Nucleic Acids Res 41, D1040–1046.

Sanchez-Mejias, E., Navarro, V., Jimenez, S., Sanchez-Mico, M., Sanchez-Varo, R., Nunez-Diaz, C., Trujillo-Estrada, L., Davila, J.C., Vizuete, M., Gutierrez, A., et al. (2016). Soluble phospho-tau from Alzheimer’s disease hippocampus drives microglial degeneration. Acta Neuropathol 132, 897–916.

Sarlus, H., and Heneka, M.T. (2017). Microglia in Alzheimer’s disease. J Clin Invest 127, 3240–3249.

Schulz, C., Gomez Perdiguero, E., Chorro, L., Szabo-Rogers, H., Cagnard, N., Kierdorf, K., Prinz, M., Wu, B., Jacobsen, S.E., Pollard, J.W., et al. (2012). A lineage of myeloid cells independent of Myb and hematopoietic stem cells. Science 336, 86–90.

Shahidehpour, R.K., Higdon, R.E., Crawford, N.G., Neltner, J.H., Ighodaro, E.T., Patel, E., Price, D., Nelson, P.T., and Bachstetter, A. (2020). Dystrophic microglia are a disease associated microglia morphology in the human brain. bioRxiv, 2020.2007.2030.228999.

Shahidehpour, R.K., Higdon, R.E., Crawford, N.G., Neltner, J.H., Ighodaro, E.T., Patel, E., Price, D., Nelson, P.T., and Bachstetter, A.D. (2021). Dystrophic microglia are associated with neurodegenerative disease and not healthy aging in the human brain. Neurobiol Aging 99, 19–27.

Sheng, J.G., Mrak, R.E., and Griffin, W.S. (1997). Glial-neuronal interactions in Alzheimer disease: progressive association of IL-1alpha+ microglia and S100beta+ astrocytes with neurofibrillary tangle stages. J Neuropathol Exp Neurol 56, 285–290.

Simmons, D.A., Casale, M., Alcon, B., Pham, N., Narayan, N., and Lynch, G. (2007). Ferritin accumulation in dystrophic microglia is an early event in the development of Huntington’s disease. Glia 55, 1074–1084.

Smith, A.M., and Dragunow, M. (2014). The human side of microglia. Trends Neurosci 37, 125–135.

Smith, L.K., He, Y., Park, J.S., Bieri, G., Snethlage, C.E., Lin, K., Gontier, G., Wabl, R., Plambeck, K.E., Udeochu, J., et al. (2015). beta2-microglobulin is a systemic pro-aging factor that impairs cognitive function and neurogenesis. Nat Med 21, 932–937.

Sobue, A., Komine, O., Hara, Y., Endo, F., Mizoguchi, H., Watanabe, S., Murayama, S., Saito, T., Saido, T.C., Sahara, N., et al. (2021). Microglial gene signature reveals loss of homeostatic microglia associated with neurodegeneration of Alzheimer’s disease. Acta Neuropathol Commun 9, 1.

Spellman, C., Ahmed, M.M., Dubach, D., and Gardiner, K.J. (2013). Expression of trisomic proteins in Down syndrome model systems. Gene 512, 219–225.

Srinivasan, K., Friedman, B.A., Etxeberria, A., Huntley, M.A., van der Brug, M.P., Foreman, O., Paw, J.S., Modrusan, Z., Beach, T.G., Serrano, G.E., et al. (2020). Alzheimer’s Patient Microglia Exhibit Enhanced Aging and Unique Transcriptional Activation. Cell Rep 31, 107843.

Streit, W.J., Braak, H., Xue, Q.S., and Bechmann, I. (2009). Dystrophic (senescent) rather than activated microglial cells are associated with tau pathology and likely precede neurodegeneration in Alzheimer’s disease. Acta Neuropathol 118, 475–485.

Streit, W.J., Khoshbouei, H., and Bechmann, I. (2020). Dystrophic microglia in late-onset Alzheimer’s disease. Glia 68, 845–854.

Streit, W.J., Sammons, N.W., Kuhns, A.J., and Sparks, D.L. (2004). Dystrophic microglia in the aging human brain. Glia 45, 208–212.

Suizu, F., Hiramuki, Y., Okumura, F., Matsuda, M., Okumura, A.J., Hirata, N., Narita, M., Kohno, T., Yokota, J., Bohgaki, M., et al. (2009). The E3 ligase TTC3 facilitates ubiquitination and degradation of phosphorylated Akt. Dev Cell 17, 800–810.

Sullivan, K.D., Lewis, H.C., Hill, A.A., Pandey, A., Jackson, L.P., Cabral, J.M., Smith, K.P., Liggett, L.A., Gomez, E.B., Galbraith, M.D., et al. (2016). Trisomy 21 consistently activates the interferon response. Elife 5.

Sultan, M., Piccini, I., Balzereit, D., Herwig, R., Saran, N.G., Lehrach, H., Reeves, R.H., and Yaspo, M.L. (2007). Gene expression variation in Down’s syndrome mice allows prioritization of candidate genes. Genome Biol 8, R91.

Suzuki, N., Suzuki, S., Millar, D.G., Unno, M., Hara, H., Calzascia, T., Yamasaki, S., Yokosuka, T., Chen, N.J., Elford, A.R., et al. (2006). A critical role for the innate immune signaling molecule IRAK-4 in T cell activation. Science 311, 1927–1932.

Svoboda, D.S., Barrasa, M.I., Shu, J., Rietjens, R., Zhang, S., Mitalipova, M., Berube, P., Fu, D., Shultz, L.D., Bell, G.W., et al. (2019). Human iPSC-derived microglia assume a primary microglia-like state after transplantation into the neonatal mouse brain. Proc Natl Acad Sci U S A 116, 25293–25303.

Taylor, J.M., Minter, M.R., Newman, A.G., Zhang, M., Adlard, P.A., and Crack, P.J. (2014). Type-1 interferon signaling mediates neuro-inflammatory events in models of Alzheimer’s disease. Neurobiol Aging 35, 1012–1023.

Taylor, J.M., Moore, Z., Minter, M.R., and Crack, P.J. (2018). Type-I interferon pathway in neuroinflammation and neurodegeneration: focus on Alzheimer’s disease. J Neural Transm (Vienna) 125, 797–807.

Teipel, S.J., and Hampel, H. (2006). Neuroanatomy of Down syndrome in vivo: a model of preclinical Alzheimer’s disease. Behav Genet 36, 405–415.

Vasileiou, P.V.S., Evangelou, K., Vlasis, K., Fildisis, G., Panayiotidis, M.I., Chronopoulos, E., Passias, P.G., Kouloukoussa, M., Gorgoulis, V.G., and Havaki, S. (2019). Mitochondrial Homeostasis and Cellular Senescence. Cells 8.

Verina, T., Kiihl, S.F., Schneider, J.S., and Guilarte, T.R. (2011). Manganese exposure induces microglia activation and dystrophy in the substantia nigra of non-human primates. Neurotoxicology 32, 215–226.

Wang, C., Yue, H., Hu, Z., Shen, Y., Ma, J., Li, J., Wang, X.D., Wang, L., Sun, B., Shi, P., et al. (2020). Microglia mediate forgetting via complement-dependent synaptic elimination. Science 367, 688–694.

Wang, J., Sanmamed, M.F., Datar, I., Su, T.T., Ji, L., Sun, J., Chen, L., Chen, Y., Zhu, G., Yin, W., et al. (2019). Fibrinogen-like Protein 1 Is a Major Immune Inhibitory Ligand of LAG-3. Cell 176, 334–347 e312.

Waugh, K.A., Araya, P., Pandey, A., Jordan, K.R., Smith, K.P., Granrath, R.E., Khanal, S., Butcher, E.T., Estrada, B.E., Rachubinski, A.L., et al. (2019). Mass Cytometry Reveals Global Immune Remodeling with Multi-lineage Hypersensitivity to Type I Interferon in Down Syndrome. Cell Rep 29, 1893–1908 e1894.

Wiseman, F.K., Al-Janabi, T., Hardy, J., Karmiloff-Smith, A., Nizetic, D., Tybulewicz, V.L., Fisher, E.M., and Strydom, A. (2015). A genetic cause of Alzheimer disease: mechanistic insights from Down syndrome. Nat Rev Neurosci 16, 564–574.

Wolf, S.A., Boddeke, H.W., and Kettenmann, H. (2017). Microglia in Physiology and Disease. Annu Rev Physiol 79, 619–643.

Xu, H., Chen, M., Manivannan, A., Lois, N., and Forrester, J.V. (2008). Age-dependent accumulation of lipofuscin in perivascular and subretinal microglia in experimental mice. Aging Cell 7, 58–68.

Xu, R., Boreland, A.J., Li, X., Erickson, C., Jin, M., Atkins, C., Pang, Z.P., Daniels, B.P., and Jiang, P. (2021). Developing human pluripotent stem cell-based cerebral organoids with a controllable microglia ratio for modeling brain development and pathology. Stem Cell Reports 16, 1923–1937.

Xu, R., Brawner, A.T., Li, S., Liu, J.J., Kim, H., Xue, H., Pang, Z.P., Kim, W.Y., Hart, R.P., Liu, Y., et al. (2019). OLIG2 Drives Abnormal Neurodevelopmental Phenotypes in Human iPSC-Based Organoid and Chimeric Mouse Models of Down Syndrome. Cell Stem Cell 24, 908–926 e908.

Xu, R., Li, X., Boreland, A.J., Posyton, A., Kwan, K., Hart, R.P., and Jiang, P. (2020). Human iPSC- derived mature microglia retain their identity and functionally integrate in the chimeric mouse brain. Nat Commun 11, 1577.

Xue, Q.S., and Streit, W.J. (2011). Microglial pathology in Down syndrome. Acta Neuropathol 122, 455–466.

Yamada, T., and Yamanaka, I. (1995). Microglial localization of alpha-interferon receptor in human brain tissues. Neurosci Lett 189, 73–76.

Yu, Q., Katlinskaya, Y.V., Carbone, C.J., Zhao, B., Katlinski, K.V., Zheng, H., Guha, M., Li, N., Chen, Q., Yang, T., et al. (2015). DNA-damage-induced type I interferon promotes senescence and inhibits stem cell function. Cell Rep 11, 785–797.

Zhao, M., Chen, L., and Qu, H. (2016). CSGene: a literature-based database for cell senescence genes and its application to identify critical cell aging pathways and associated diseases. Cell Death Dis 7, e2053.

